# Proton-Activated Chloride Channel 1 (PACC1) is essential for innate host defense against bacterial sepsis

**DOI:** 10.64898/2025.12.31.697179

**Authors:** Lucien P Garo, Kevin Brueck, Sarah Walachowski, Archana Jayaraman, Marcel Strueve, Shuang Xu, Hulbert Yang, Matthew Helmkamp, Seung Hoan Choi, Christoph Reinhardt, Markus Bosmann

## Abstract

Bacterial sepsis remains a devastating clinical problem. Here, we describe a protective role for the recently discovered acid-sensitive, proton-activated chloride channel, PACC1 (PAC/ASOR/TMEM206), during sepsis. Initially, we found PACC1 was enriched in healthy human and mouse mononuclear phagocytes, particularly macrophages, and differentially regulated by inflammatory stimuli, suggesting PACC1 involvement in innate immunity. To further investigate, we generated *de novo Pacc1* knockout (^-/-^) mice, which presented without major immunologic abnormalities at baseline. Compared to wildtype (WT), *Pacc1^-/-^* myeloid cells showed normal phagocytic uptake of acid-insensitive *Escherichia coli BioParticles*, but impaired development of the acidifying phagolysosome using acid-sensitive *E. coli* BioParticles. Transcriptomic profiling of *Pacc1*^-/-^ macrophages revealed dysregulated phagolysosomal and cytokine networks (e.g., interferons). Because phagolysosomal bacterial clearance is essential to resolve infection, we challenged *Pacc1^-/-^* mice with intraperitoneal gram-negative *E. coli* sepsis. *Pacc1^-/-^* mice displayed increased bacterial burden, immune cell infiltration, inflammation, and lethality. In contrast, phagocytosis-independent *E. coli* lipopolysaccharide (LPS)-induced endotoxemia yielded comparable WT and *Pacc1^-/-^* survival, as well as similar inflammatory responses. Finally, we engineered *Pacc1*-floxed (^fl/fl^) mice crossed with a myeloid lineage Cre-deleter strain to interrogate myeloid cell-intrinsic PACC1 *in vivo*. Consistent with a predominate role for PACC1 during phagocytosis and bacterial clearance in these cells, *LysM-Cre/Pacc1^fl/fl^* mice exhibited impaired *E. coli* sepsis survival but indifferent endotoxemia phenotypes. In conclusion, PACC1 links sterilizing phagolysosomal activity with immune networks in sepsis pathobiology.

**Significance Statement:** Bacterial sepsis remains a major global health burden. Here, we report an essential role for the recently discovered acid-sensitive chloride channel, PACC1 (PAC/ASOR/TMEM206), in protective host defense during bacterial infection and sepsis. PACC1 is highly expressed in human and mouse phagocytic myeloid cells, particularly macrophages, where it regulates phagocytic bacterial clearance and inflammatory responses. Using *de novo* generated mice, we show that global or myeloid cell-targeted deletion of PACC1 impairs development of phagolysosomal acidification, confers susceptibility to bacterial infection and excessive inflammation, and undermines host defense. These findings warrant further investigation of PACC1 in sepsis pathobiology.

## Introduction

Life-threatening bacterial infections are a devastating clinical problem, and the molecular mechanisms of protective host defense require continued investigation. An estimated ∼50 million annual cases of sepsis worldwide have mortality rates of 20-50% (1). Strikingly, sepsis may account for up to 50% of US hospital deaths and is thought to be the most expensive condition affecting the US healthcare system (2, 3). Pneumonia is a major risk factor for sepsis, causing approximately 50% of all cases (4) and is itself a leading cause of death in children under five and in the elderly (5). Bacterial infections cause the majority of sepsis deaths (6), despite a recent surge in viral sepsis due to COVID-19 (7). Gram-negative *Escherichia coli* is a major pathogen of systemic bacteremia, especially in higher income countries, and the gram-positive bacterium, *Streptococcus pneumoniae*, is the most common etiology of bacterial pneumonia (8). Unfortunately, antibiotic resistance has been increasing amongst bacterial pathogens, and infection-associated mortality in the US has not significantly improved since the widespread use of antibiotics (6, 9). Treatment of bacterial infections is further complicated by limited diagnostic capabilities—the exact infectious microbe remains unidentified in a plurality of bacterial sepsis cases (6). Due to these ongoing public health threats, our laboratory (10–13) and many others (5, 14, 15) have been investigating broadly applicable host defense mechanisms important for innate immune responses to bacterial infections.

Tissue-resident macrophages mediate initial host defense against invading bacterial infections and interface at mucosal/serosal sites (16). Phagocytosis by these surveilling macrophages and other myeloid cells which are rapidly recruited to the site of infection, including neutrophils and monocytes, are essential for preventing and limiting infection severity (17, 18). During phagocytosis, bacteria at the plasma membrane are internalized in a receptor-mediated process into the membrane-bound phagosome, which is followed by maturation and acidification of this compartment, culminating in fusion with the acidic lysosome to form the phagolysosome. The strongly acidic pH of the phagolysosome and a cocktail of proteolytic enzymes kill bacteria (17, 18). Thus, the study of host defense mechanisms related to phagolysosomal development and function are of major relevance to bacterial infections.

Chloride is the most abundant free anion in humans and governs biological functions ranging from intercellular signaling to intracellular vesicular acidification (19, 20). Decades of research has uncovered many subtypes of cellular/intracellular membrane chloride channels and transporters (19). To cross cellular membranes, chloride requires channels for passive diffusion, or specific proteins to actively transport chloride in conjunction with other ions. Chloride channels/transporters have been linked to the etiology of many diseases and implicated as therapeutic targets in cystic fibrosis, neurodegeneration, epilepsy, osteopetrosis, hypertension, and infection (19, 20).

The molecular identity of a widely expressed acid-activated chloride channel, PACC1 (also known as PAC, ASOR, TMEM2026) has recently been identified in two independent genome-wide siRNA screens (21, 22), validating years of electrophysiological experimental data that had suggested the existence of such a channel (21–25). PACC1 channels are homotrimers of TMEM206 subunits that each contain two transmembrane domains (21, 22, 26–29). Mutagenesis and cryo-electron microscopy studies have shown that all six transmembrane domains of the trimer participate in forming a chloride-selective pore that is open upon acid-induced contraction of their extracellular/luminal domains (21, 22, 26–29). This results in a steep pH dependence with PACC1 being closed at neutral pH and maximally activated at pH 4.5, in addition to a steep voltage dependence (21–29).

Due to its unique acid sensitivity, studies initially focused on how PACC1 promotes acid-induced cell death (21, 22, 25, 30, 31), particularly in the brain where PACC1 is highly expressed and where extracellular pH can drop during ischemic stroke (30, 31). However, beyond acidosis, the role of PACC1 in health and disease remains largely unknown.

PACC1 is ubiquitously expressed and has been detected in all mammalian cells (21, 22). PACC1 is also highly evolutionarily conserved, with >90% conservation between mice and humans (21, 22), suggesting essential biological functions. Beyond the established role of PACC1 in acidotoxicity, stroke, and regulating responses to extracellular acidification (21, 22, 30, 32), its primary function may be the regulation of intracellular vesicles whose acidic pH allows PACC1 activity (20, 33). Indeed, after its identification, PACC1 has been characterized in these compartments (21, 22, 34–37) and recent studies have identified a role for PACC1 in endosomal acidification and macropinosome resolution (34–37).

Here, we hypothesize PACC1 is essential in the phagolysosome, which contains a highly acidic compartment optimal for PACC1 activation (28) with one of the highest concentrations of chloride (20). We found PACC1 in macrophages is required for effective bacterial eradication, inflammatory responses, and host resistance against sepsis.

## Results

### Human PACC1 is expressed abundantly in macrophages and other mononuclear phagocytes

PACC1 is ubiquitously expressed and has been detected in every mammalian cell type (21, 22). To explore where PACC1 may play a more predominant role, we first analyzed publicly available human single-cell RNA sequencing (scRNA-seq) data from the Human Protein Atlas (HPA) (38). The HPA scRNA-seq is based on 31 published datasets, and expression profiles were generated by converting read counts to normalized transcripts per million (nTPM). Amongst all cells, *PACC1* expression was highly enriched within immune cells (**Fig. 1A**). Moreover, within immune cells in peripheral blood, *PACC1* was preferentially expressed within phagocytic mononuclear myeloid cells, such as monocytes and dendritic cells, both in the HPA dataset (**Fig. 1B**), as well as by independent bulk RNA-seq datasets from the Immune Cell Atlas (**Fig. 1C**) (39). PACC1 was also highly elevated in glia, which include brain-resident microglial “professional” phagocytes, as well as astrocytes and oligodendrocytes with phagocytic capacity (**Fig. 1A**) (40). To determine whether PACC1 was likewise expressed in tissue-resident macrophages, the predominant mononuclear phagocytes that initially interface with bacteria during infection (12, 13, 15, 16, 41), we next examined HPA data from the 19 tissues with annotated tissue macrophage populations (**Fig. 1D**). *PACC1* was consistently enriched in immune cells compared to non-immune cells. Most strikingly, tissue macrophages were the highest *PACC1*-expressing cell type in almost every tissue. These included Kupffer cells and Hofbauer cells, which are specialized tissue macrophages in the liver and placenta, respectively; and Langerhans cells, which are specialized tissue macrophages/dendritic cells in the skin (16). In addition, HPA bulk RNA-seq data from 69 cell lines of various tissue origin revealed that myeloid lineage cell lines were enriched in the top 50% in terms of ranked *PACC1* expression (***SI Appendix,* Fig. S1A**). The highest *PACC1* expression was in THP-1 cells, and the third highest expression in U-937 cells, the most widely used cell lines for macrophage studies (42). In addition, we analyzed an expanded pool of cell lines that combined HPA bulk RNA-seq with data from >1000 cell lines in the Cancer Cell Line Encyclopedia (CCLE) and a Genentech study (43). The top 2% of *PACC1*-expressing cells contained a plurality of myeloid lineage cells, including THP-1 cells (***SI Appendix,* Fig. S1B**).

**Fig. 1.**
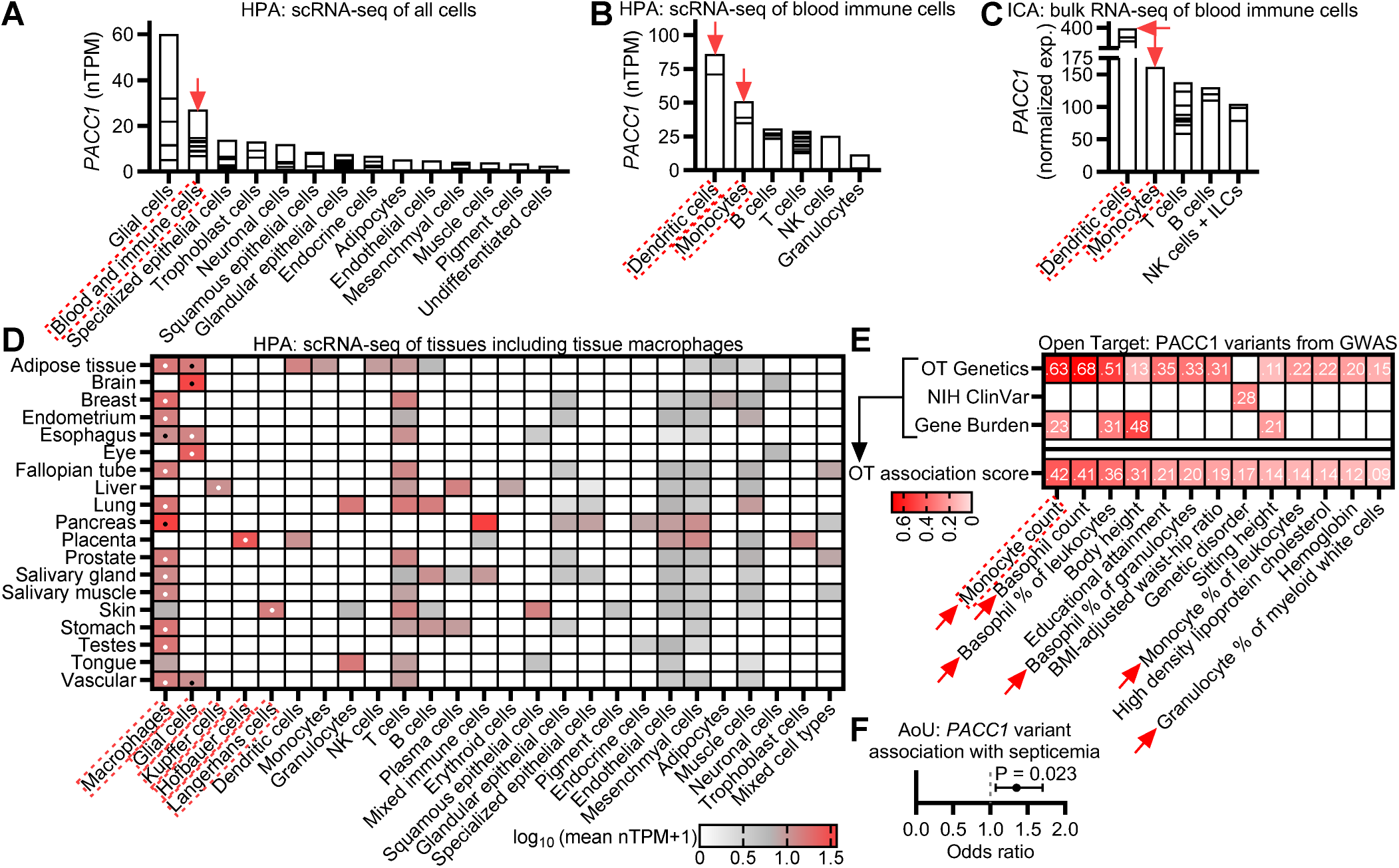
*PACC1* expression is enriched in human mononuclear phagocytes, particularly tissue macrophages, and *PACC1* variants are implicated in myeloid cell-mediated immunity. **A-D** Single-cell RNA sequencing (scRNA-seq) data from the Human Protein Atlas (HPA) or bulk RNA-seq data from the ImmGen Immune Cell Atlas (ICA), as indicated. Curated cell types along the x-axis contain *superimposed* (not stacked) data from cells that were clustered and annotated (HPA), or sorted (ICA), such that each horizontal line on the bar graphs represents the gene expression for an individual cell type/cluster. HPA data are normalized transcripts per million (nTPM), corresponding to means of different individual samples from each cell type. ICA data are mean normalized counts from different individual samples. Red arrows/boxes indicate cells of interest within each panel (e.g., immune cells, mononuclear phagocytes, or macrophages). (**A**) *PACC1* expression in somatic human cell types from the HPA. (**B**) *PACC1* expression in blood immune cells from the HPA. (**C**) *PACC1* expression in blood immune cells from the ICA. (**D**) *PACC1* expression in all 19 somatic tissues from the HPA wherein tissue macrophages were annotated. Cell clusters are broadly curated in combined columns (e.g., epithelial cells, endothelial cells), while immune cell clusters are resolved in separate columns (e.g., macrophages, granulocytes). Data represent log_10_(mean nTPM + 1) corresponding to means of different clusters from each cell type. Dots indicate tissue macrophage-containing cell types that rank first (white) and/or second (black) in terms of PACC1 expression amongst all cell types within each tissue. (**E**) Heatmap ranking showing the most well-evidenced *PACC1* genetic variant associations across all phenotypes, according to the Open Targets Platform for genome-wide association studies (GWAS), ranked by “association scores” which synthesize evidence across databases (e.g., OT Genetics, NIH ClinVar, Gene Burden). Red arrows/boxes indicate *PACC1* variant associations with phenotypes of interest (i.e., myeloid cell-related phenotypes). (**F**) Forest plot showing association between gene-level predicted loss-of-function variant burden (low, rare and ultra rare, minor allele frequency <1%) in *PACC1* with septicemia phecode in the AoU dataset. The plot shows burden test P-value, odds ratio, and 95% confidence interval.

To further assess whether PACC1 may be implicated in myeloid cell functions, we next analyzed *PACC1* variant data from publicly available genome-wide association studies (GWAS). Although no specific studies have functionally characterized any human *PACC1* genetic variants, we found multiple GWAS have identified *PACC1* variant associations with myeloid cell-related phenotypes. According to the Open Target GWAS aggregator (44), among all biological phenotypes associated with *PACC1* variants, myeloid cell-related phenotypes were highly represented. “Monocyte counts” ranked as the most supported phenotype (**Fig. 1E**). Closer examination revealed a modest but consistent pattern: most *PACC1* variants were predicted to be loss-of-function (pLoF), and these pLoF variants were associated with greater myeloid cell counts/frequencies, whereas the few predicted gain-of-function (pGoF) variants were associated with the converse (***SI Appendix,* Table S1**). Furthermore, the recent large *All of Us* (AoU) cohort revealed the association of septicemia with gene-level burden of low frequency *PACC1* pLoF variants (**Fig. 1F**) (45). A separate large population cohort from the UK Biobank (46), which used a broader assortment of clinical codes, showed an overall modest trend in association between *PACC1* gene-level variant burden and several codes pertaining to predominantly bacterial infections (***SI Appendix,* Fig. S2**). In summary, these data demonstrate that *PACC1* expression in humans is highly enriched in mononuclear phagocytes, particularly macrophages, and that *PACC1* variants may impact myeloid cell biology and innate immunity.

### Pacc1 expression in mice is enriched in macrophages and other mononuclear phagocytes

To further evaluate PACC1 in macrophages, we proceeded with mouse studies. We first sought to determine whether *Pacc1* was also highly expressed in murine macrophages. Reanalysis of bulk RNA-seq data from our previous sepsis study revealed *Pacc1* expression was ranked in the top 5% of all genes in murine bone marrow-derived macrophages (BMDMs) at baseline (**Fig. 2A**) (11). Since monoclonal antibodies for the PACC1 protein were not commercially available (47), we next employed mRNA PrimeFlow (48) to specifically detect *Pacc1* mRNA across macrophage and splenocyte populations by flow cytometry. As negative controls, we leveraged cells from our *de novo* generated *Pacc1* knockout mice (***SI Appendix,* Fig. S3**), or WT cells without *Pacc1* target probe. Using this PrimeFlow approach, robust *Pacc1* expression was first validated in murine BMDMs (**Fig. 2B**; gating scheme in ***SI Appendix,* Fig. S4A**), as also recently confirmed at the protein level (35). Consistent with human data (**Fig. 1B-C**), *Pacc1* expression in murine splenocytes was detectable in mononuclear phagocytes, including macrophages, dendritic cells, and a small population of neutrophils, as well as B cells, which also possess phagocytic capacity (18) (**Fig. 2C**; gating scheme in ***SI Appendix,* Fig. S4B**). Overall, *Pacc1* was enriched in mononuclear phagocytes compared to other splenocytes (***SI Appendix,* Fig. S4C**). These findings were further corroborated by reverse transcription quantitative-PCR (RT-qPCR) in fluorescence-activated cell (FAC)Sorted splenic myeloid cell populations, which showed higher expression in macrophages, modest expression in monocytes, and detectable expression in polymorphonuclear neutrophils (**Fig. 2D**). To ascertain PACC1 levels across a broader set of murine tissue macrophages, we analyzed published transcriptomic data (49). Relevant to pneumonia and peritoneal sepsis (16), *Pacc1* was highly expressed in peritoneal macrophages and alveolar macrophages (**Fig. 2E**). Next, we analyzed single-nucleus sequencing data to examine the peritoneal compartment. *Pacc1* was highly expressed within murine macrophages and other mononuclear phagocytes, (including monocytes and dendritic cells), of naïve mice (**Fig. 2F-G**). Finally, we tested if *Pacc1* in macrophages was differentially regulated in response to inflammatory stimuli associated with infection, such as Toll-like receptor (TLR) ligands. In BMDMs, *Pacc1* expression was differentially responsive to such stimuli (**Fig. 2H**). Of note, *Pacc1* was strongly downregulated by lipopolysaccharide (LPS), a component of gram-negative bacterial cell walls (12). Direct stimulation of macrophages with inactivated *E. coli* also inhibited *Pacc1* (**Fig. 2I**). Collectively, these data show that *Pacc1* expression is abundant in human and mouse macrophages, and differentially regulated by inflammatory stimuli, further supporting that PACC1 could be involved in innate immune responses to infection.

**Fig. 2.**
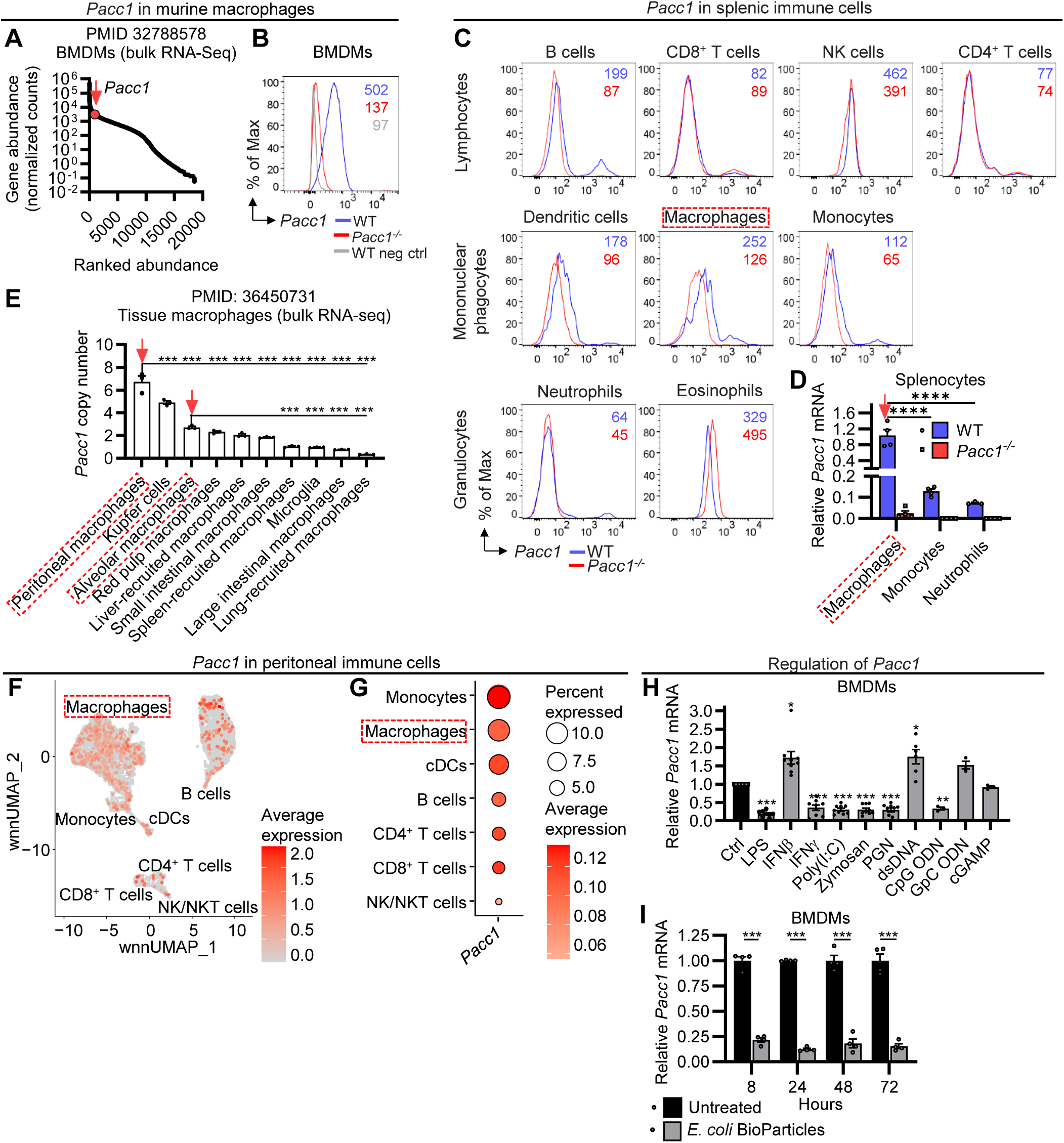
*Pacc1* expression is highly expressed in murine macrophages and other mononuclear phagocytes. (**A**) Ranked gene expression of bulk RNA sequencing (bulk RNA-seq) data of bone marrow-derived macrophages (BMDMs) isolated *ex vivo* from wildtype (WT) mice. Data reanalyzed from our previous study (*11*). *Pacc1* expression shown in red in the top 5%. (**B-C**) *Pacc1* mRNA fluorescence by RNA PrimeFlow staining in splenocytes from WT and *Pacc1* knockout (*Pacc1*^-/-^) mice. Geometric mean fluorescence intensities (MFIs) are indicated in the upper right. (**B**) Representative flow cytometric histograms of *Pacc1* expression in resting bone marrow-derived macrophages (BMDMs). Neg ctrl indicates no target probe. (**C**) Representative flow cytometric histograms of *Pacc1* expression in splenocytes. (**D**) *Pacc1* expression in selected FACSorted splenocyte myeloid cell populations (live CD11b^+^F4/80^+^ macrophages, CD11b^+^Ly6C^+^Ly6G^-^ monocytes, and CD11b^+^Ly6C^+^Ly6G^+^ neutrophils) by RT-qPCR normalized to WT macrophage expression. (**E**) *Pacc1* copy number in bulk RNA-seq data from tissue macrophages isolated from WT mice. Data reanalyzed from Qie *et al.* (49). (**F-G**) *Pacc1* expression by single-nucleus RNA sequencing of peritoneal cells from WT mice. (**F**) *Pacc1* expression in annotated immune cell clusters visualized on the weight-nearest neighbor uniform manifold approximation and projection (wnnUMAP) plot. (**G**) Dot plot of *Pacc1* expression from **F**. (**H**) *Pacc1* expression in stimulated WT BMDMs analyzed by RT-qPCR, normalized to untreated control. In brief, BMDMs were stimulated for 24 h with various stimuli corresponding to inflammatory/infectious insults, including lipopolysaccharide (LPS), interferon-beta (IFN-β), interferon-gamma (IFN-ɣ), polyinosinic:polycytidylic acid (poly I:C), zymosan, peptidoglycan (PGN), double-stranded DNA (dsDNA), CpG oligodeoxynucleotides (ODN), GpC ODN, or cyclic guanosine monophosphate–adenosine monophosphate (cGAMP). (**I**) *Pacc1* expression in BMDMs stimulated with inactivated *E. coli* Bioparticles analyzed by RT-qPCR, normalized to untreated control at each time point. For **B**-**D**, **H**, **I**, data represent ≥2 independent experiments. For **D**, **E**, **H**, **I**, data show one symbol per mouse, mean±SEM, *P<0.05, **P<0.01, ***P<0.001, ****P<0.0001,one-way ANOVA with Tukey (**D**), one-way ANOVA with Sidak (**E**, **I**), two-way ANOVA with Tukey (**D**), or one-way ANOVA with Dunett comparing each to unstimulated control (**H**). For **C**-**G**, red arrows/boxes indicate cells of interest (i.e., macrophage populations).

### De novo generation of Pacc1 knockout mice

To further study the role of PACC1 *in vivo*, we generated mice with a global deletion of *Pacc1* using a CRISPR/Cas-9 approach (***SI Appendix,* Fig. S3A**). Single-guide RNAs aligned to regions flanking *Pacc1* exon 2 were injected into pronuclei/zygotes (C57BL/6J), resulting in a 120 base pair deletion with a frame shift and a premature stop codon. This truncated exon was detectable by PCR (***SI Appendix,* Fig. S3B**). In addition, RT-qPCR for exons 2-8 verified absent *Pacc1* mRNA in macrophages (***SI Appendix,* Fig. S3C**), suggesting nonsense-mediated mRNA decay (50). Acid-induced cytotoxicity was lower in *Pacc1^-/-^* BMDMs (***SI Appendix,* Fig. S3D**), confirming loss of PACC1 protein function in line with previous electrophysiology studies (21–25). In agreement with other independently generated mice (21, 22), our *Pacc1^-/-^*strain was viable and fertile, displaying normal sex distributions, although the frequency of homozygous offspring was slightly reduced from expected Mendelian ratios (25%) in heterozygous breeding schemes (***SI Appendix,* Fig. S3E**). *Pacc1^-/-^* mice presented without gross abnormalities. Males and females had normal body weights and clinical chemistry parameters including blood proteins, metabolites, and electrolytes (***SI Appendix,* Fig. S3F-I**). Importantly, *Pacc1^-/-^* mice housed under specific pathogen-free conditions showed no apparent immunologic abnormalities, with normal numbers of peripheral myeloid and lymphoid cells in the blood and spleen (***SI Appendix,* Fig. S3J-L**, gating scheme in ***SI Appendix,* Fig. S5**). Together, these data suggest that *Pacc1^-/-^* mice lack developmental immunologic abnormalities, making them appropriate to study bacterial infections *in vivo*.

### Pacc1^-/-^ mice have defective development of phagolysosomal acidification in vitro and in vivo

Acid-triggered PACC1 may play an active role in the acidifying phagolysosome of myeloid cells (28). To study if bacteria induce PACC1-containing phagosomes in macrophages, we re-analyzed published proteomic data: Li *et al.* used an engineered ascorbate peroxidase (APEX)-based proximity labeling technique combined with mass spectrometry in murine macrophages to identify phagosomal host proteins following phagocytosis of APEX-tagged inactivated gram-negative *E. coli* and gram-positive *Staphylococcus aureus* (51). Indeed, PACC1 protein was enriched in bacteria-containing phagosomes in BMDMs (**Fig. 3A**). Next, we tested the development of phagolysosomal acidification in *Pacc1^-/-^* mice using inactivated *E. coli* BioParticles conjugated with an acid-sensitive fluorophore. These phagocytosed BioParticles become fluorescent at low pH as the phagosome fuses with the lysosome (52). We found BioParticles incubated with *Pacc1^-/-^*BMDMs displayed decreased fluorescence compared to WT counterparts (**Fig. 3B**). These data suggested a defect in the development of the acidifying phagolysosome, as normal phagocytic engulfment of synthetic beads by macrophages has been reported not to require PACC1 (35), in accordance with PACC1 inactivity at neutral plasma membrane pH (22, 26–29). We also confirmed normal phagocytic uptake of *E. coli* cargo in *Pacc1^-/-^* macrophages by exposure to acid-insensitive Alexa Fluor (AF)488-conjugated *E. coli* BioParticles. AF488-*E. coli* emit a stable signal from pH 3-10 independent of phagolysosomal acidification (53). AF488 fluorescence in *Pacc1^-/-^*macrophages was indistinguishable from WT, and signals in both strains were abrogated by cytochalasin D, an inhibitor of actin polymerization and phagocytic uptake (53, 54) (**Fig. 3C-D**). Accordingly, *Pacc1^-/-^*macrophages exhibited impaired intracellular killing of live pathogenic *E. coli*, while levels of extracellular bacteria were comparable to WT cultures (**Fig. 3E**), indicating intact phagocytic uptake but defective phagolysosomal killing. When acid-sensitive *E. coli* BioParticles were injected intraperitoneally (i.p.) in mice, a similar reduction in fluorescence in *Pacc1^-/-^* vs. WT myeloid cells was noted (**Fig. 3F-G**). Additionally, *Pacc1^-/-^* mice displayed heightened inflammatory responses to inactivated BioParticles, characterized by increased recruitment of myeloid cells to the peritoneum, including monocytes and monocyte-derived macrophages (**Fig. 3H**, gating scheme in ***SI Appendix,* Fig. S6**). IL-6, a major pro-inflammatory cytokine (12), was also elevated in peritoneal lavage fluid of *Pacc1^-/-^* mice injected with BioParticles (**Fig. 3I**). In contrast, no aberrant signs of pre-existing inflammation were present in the peritoneum of naïve *Pacc1^-/-^* mice (***SI Appendix,* Fig. S7**), consistent with immunoprofiling in the periphery (***SI Appendix,* Fig. S3J-L**). Both naïve WT and *Pacc1^-/-^* mice had low numbers of infiltrating peritoneal monocytes and neutrophils, with modestly fewer monocytes in *Pacc1^-/-^* mice (***SI Appendix,* Fig. S7A-B**); and IL-6 was undetectable in both strains (***SI Appendix,* Fig. S7C**). As a whole, these data suggest acidification of the phagolysosome is defective in *Pacc1^-/-^*myeloid cells, and that PACC1 deficiency may result in hyperinflammation following bacterial challenge.

**Fig. 3.**
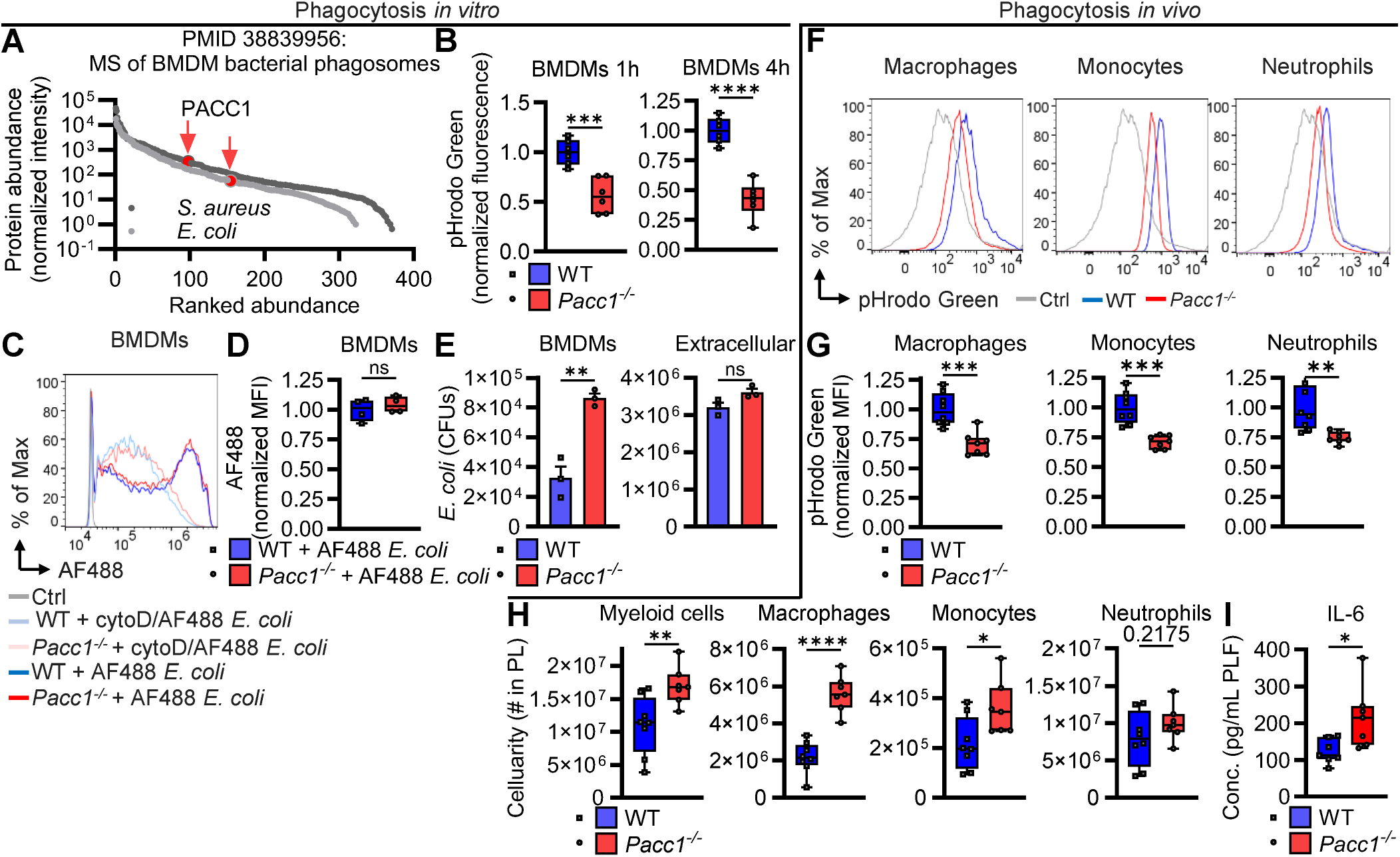
*Pacc1^-/-^* mice have defective development of the acidifying phagolysosome. (**A**) Ranked protein abundance from mass spectrometry (MS) data of the phagosomes of bone-marrow-derived macrophages (BMDMs) stimulated with bacteria. Data reanalyzed from Li *et al.* (51). In brief, BMDMs were stimulated with ascorbate peroxidase (APEX)-modified inactivated bacteria followed by APEX-mediated proximity ligation, and subsequently MS, to identify bacterial phagosomal proteins. PACC1 is highlighted in red. (**B**) Wildtype (WT) and *Pacc1^-/-^* BMDMs were treated with 330 µg/mL acid-sensitive pHrodo Green *E. coli* BioParticles for the indicated time points. Fluorescence was measured by plate-reader and normalized to WT for each time point to assay development of phagolysosomal acidification. (**C-D)** WT and *Pacc1^-/-^* BMDMs were treated with acid-insensitive Alexa Fluor (AF)488 *E. coli* BioParticles. After 1 h, AF488 geometric mean fluorescence intensity (MFI) was measured by flow cytometry to assay phagocytic uptake. (**C**) Representative flow cytometric histograms of BMDMs are shown. CytoD indicates negative control cells pre-treated with actin inhibitor, cytochalasin D, for 1 hour at 10μM to limit phagocytic uptake. Ctrl indicates negative control WT cells treated with inactivated *E. coli* without fluorescent conjugation. (**D**) MFI quantifications from panel **C**, normalized to WT. (**E**) Colony-forming unit (CFU) counts from WT and *Pacc1^-/-^*BMDMs infected with pathogenic *E. coli* serotype O6:K2:H1 at an MOI of 5 for 2 h. CFUs in macrophage lysates and extracellular supernatants via serially diluted plated cultures to assay bacterial killing capacity. (**F-I**) WT and *Pacc1^-/-^* mice were injected i.p. with 100 µg pHrodo Green *E. coli* BioParticles. After 24 h, cellularity and pHrodo Green MFI were measured by flow cytometry to assay development of phagolysosomal acidification from peritoneal lavage cells. (**F**) Representative flow cytometric histograms of live CD11b^+^Ly6C^-/lo^F4/80^+^ macrophages, CD11b^+^Ly6C^hi^Ly6G^-^ monocytes, and CD11b^+^Ly6C^+^Ly6G^+^ neutrophils are shown. Ctrl indicates fluorescence of total live cells from an unstimulated WT mouse. (**G**) MFI quantifications from panel **F**, normalized to WT. (**H**) Cell numbers from the peritoneal lavage (PL) of the same *E. coli* experiments above, including total live CD11b^+^ myeloid cells by flow cytometry. (**I**) IL-6 concentration in peritoneal lavage fluid (PLF) by ELISA from the same *E. coli* experiments above. For **B**-**I**, data represent ≥2 independent experiments. For **B**, **D**, **E**, **G**-**I**, data show one symbol per mouse. Bar graphs depict mean±SEM, and box plots show median and interquartile range with whiskers extended to the minimum and maximum. *P<0.05, **P<0.01, ***P<0.001, ****P<0.0001, ns=not significant, Student’s t-test.

### Pacc1^-/-^ macrophages have dysregulated immunologic and phagolysosomal transcriptomic responses

To screen for the transcriptome-wide effects of *Pacc1* loss on host defense and phagolysosomal pathways, we sequenced RNA from BMDMs stimulated with *E. coli* BioParticles. Principal component analysis revealed strong segregation based on *E. coli* stimulation, as well as segregation based on genotype (**Fig. 4A**). *Pacc1^-/-^* control compared to WT control samples were distinguishable at baseline and further segregated following phagocytic stimulation. *E. coli* triggered robust transcriptomic activity, with >10,000 significant differentially expressed genes (DEGs) in each group of WT and *Pacc1^-/-^*macrophages (***SI Appendix,* Fig. S8A-C; Dataset S1**). In response to *E. coli*, *Pacc1^-/-^*vs. WT macrophages displayed 2,162 upregulated and 1,795 downregulated DEGs (**Fig. 4B**). WT and *Pacc1^-/-^*macrophages also displayed a smaller proportion of DEGs at baseline (***SI Appendix,* Fig. S8D**); however, the majority were uniquely dysregulated by *E. coli* (**Fig. 4C**).

**Fig. 4.**
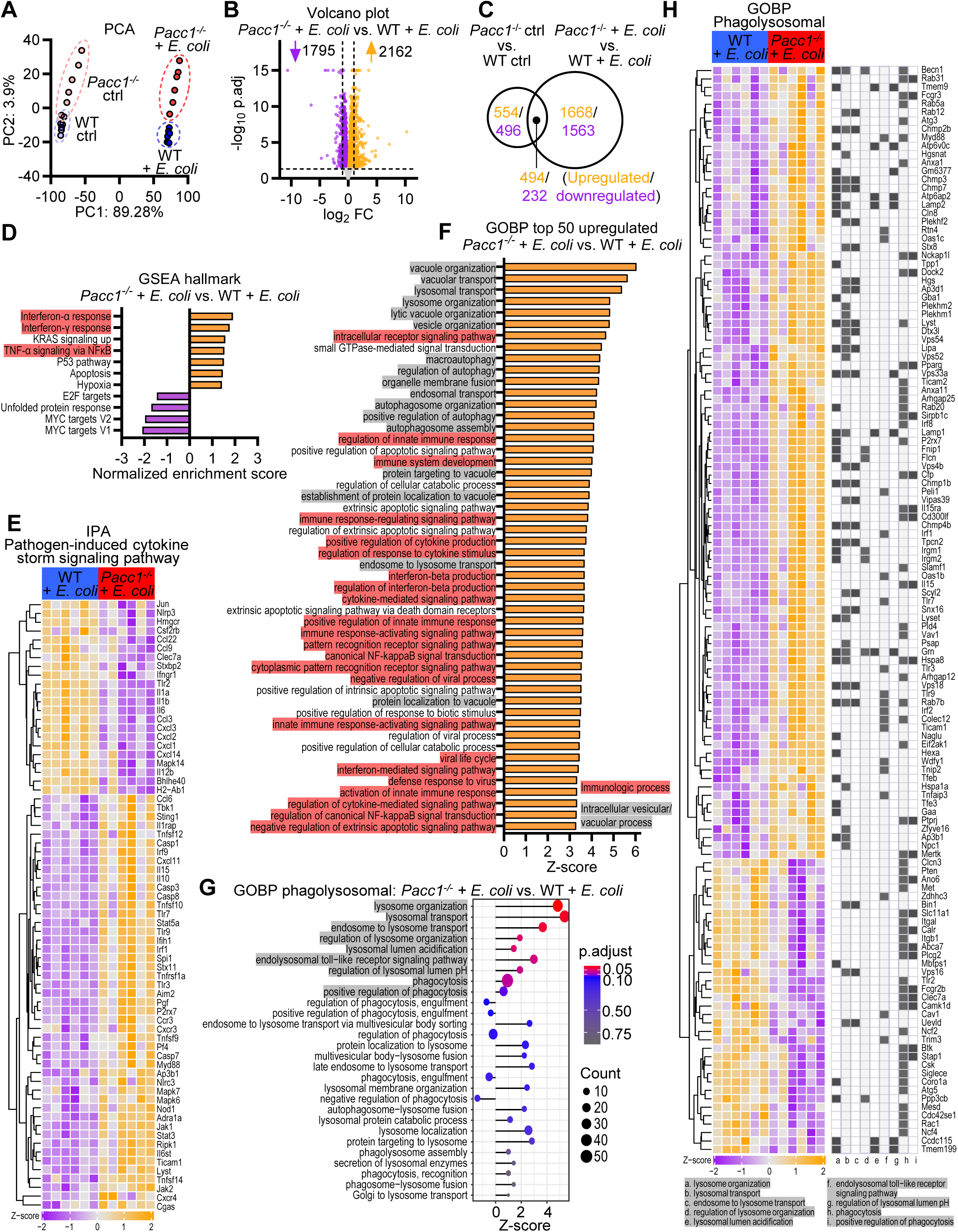
Transcriptomic profiling of dysregulated immunologic and phagolysosomal pathways in bone marrow-derived macrophages (BMDMs) from *Pacc1^-/-^* mice and wildtype mice stimulated with *E. coli* Bioparticles. Wildtype (WT) and *Pacc1^-/-^* BMDMs were stimulated with *E. coli* BioParticles at 100 µg/mL or left as unstimulated controls (ctrl) for 12 h and subjected to bulk RNA sequencing (n=6 mice/group). (**A**) Principal component analysis (PCA) plot of the top 5000 variable genes after variance-stabilizing transformation of sequencing counts, displaying *E. coli*-dependent and genotype-dependent clusters. (**B**) Volcano plot of differentially expressed genes (DEGs) in *Pacc1^-/-^* vs. WT BMDMs following *E. coli* stimulation. Dotted lines indicate adj. P=0.05 and absolute log_2_ fold change (FC)=1; values for -log_10_ adj. P>15 were set=15 for visualization. (**C**) Venn diagram showing common and specific significant DEGs in *Pacc1^-/-^* vs. WT BMDMs with or without *E. coli*. (**D**) Bar plot showing hallmark gene set enrichment analysis (GSEA) in *Pacc1^-/-^* vs. WT BMDMs with *E. coli*. Immunologic pathways in red. (**E**) Heatmap of significant DEGs enriched in “Pathogen Induced Cytokine Storm Signaling Pathway” from Ingenuity Pathway Analysis (IPA) in *Pacc1^-/-^*vs. WT BMDMs with *E. coli*. Normalized expression values. (**F**) Bar plot showing top 50 significant (adj. P<0.1) positively enriched gene ontology biological processes (GOBPs) in *Pacc1^-/-^* vs. WT BMDMs with *E. coli*. Immunologic pathways are in red, intracellular vesicular/vacuolar pathways in gray. (**G**) Lollipop plot of manually curated phagolysosomal GOBPs showing both enriched (adj. P<0.1) and non-enriched in *Pacc1^-/-^* vs. WT BMDMs with *E. coli*. (**H**) Heatmap of significant DEGs within enriched (adj. P<0.1) phagolysosomal GOBPs from panel **G** in *Pacc1^-/-^* vs. WT BMDMs with *E. coli*. Genes enriched within each GOBP are indicated by shaded squares. Heatmaps show normalized expression values with Z-score from -2/purple to +2/orange and significance at adj. P<0.05. See also *SI Appendix,* Datasets 1–4.

To assess global changes, we performed a gene set enrichment analysis (GSEA) for hallmark biological processes using both significant and non-significant DEGs. Immunologic gene sets were differentially enriched in *Pacc1^-/-^* vs. WT macrophages both at baseline (***SI Appendix,* Fig. S9A; Dataset S2**) and following *E. coli* phagocytosis (**Fig. 4D; *SI Appendix,* Dataset S2**), including TNF-α via NFκB signaling (***SI Appendix,* Fig. S9B-C**) and interferon responses (***SI Appendix,* Fig. S9D-G**), as well as JAK/STAT signaling (***SI Appendix,* Fig. S9A, H**). Further analysis via the “Interferome database”, a curated list of empirically determined interferon-responsive genes (55), revealed strong dysregulation of type I and II interferon signaling in *Pacc1^-/-^* macrophages following bacterial challenge (***SI Appendix,* Fig. S10**).

Ingenuity Pathway Analysis (IPA) identified dysregulated signaling in *Pacc1^-/-^* macrophages in response to *E. coli* (***SI Appendix,* Dataset S3**). Notably, the “pathogen-induced cytokine storm signaling pathway” was one of the most highly upregulated pathways in *Pacc1^-/-^* vs. WT macrophages (**Fig. 4E**). Uncontrolled cytokine release is a leading contributor to death in sepsis patients (15). DEGs enriched in this pathway resolved that this activation response was not uniform: expression of some inflammatory mediators were elevated in *Pacc1^-/-^*vs. WT macrophages following *E. coli* exposure (e.g., *Il1rap, Ccl6*, *Pf4, Ripk1, Jak1, Stat3, Tlr9, Tlr3, Cgas, Nod1,* and *Sting1*), while others were downregulated (e.g., *Il6*, *Il1b*, *Cxcl1*, and *Cxcl2*).

Next, to study dysregulated biological processes in *Pacc1^-/-^*macrophages at a more granular level, including phagolysosomal functions, we analyzed the enrichment of gene ontology biological processes (GOBPs) for significant DEGs (***SI Appendix,* Dataset S4**). Amongst the top upregulated GOBPs, a majority were associated with either innate immunity or intracellular vacuoles/vesicles (**Fig. 4F**). More than 100 DEGs were enriched in “regulation of innate immune response” processes in *Pacc1^-/-^*vs. WT macrophages, the majority of which were associated with positive regulation, both following *E. coli* incubation and at baseline (***SI Appendix,* Fig. S11**). In contrast, the top downregulated pathways in *Pacc1^-/-^* vs. WT macrophages during phagocytosis were related predominantly to RNA processing (***SI Appendix,* Fig. S12**).

To determine if *Pacc1* loss affected downstream phagolysosomal regulation, we manually curated a list of annotated processes. Consistent with an important role for PACC1 during bacterial challenge, multiple GOBPs associated with lysosomal activity were differentially regulated in *Pacc1^-/-^* vs. WT macrophages (e.g., lysosomal organization/transport, lysosomal lumen acidification; adj. *P*<0.1), while differences in GOBPs associated with early phagocytic processes (e.g., engulfment) were not significant (**Fig. 4G**). Interestingly, a majority of these enriched DEGs were upregulated (**Fig. 4H**), which may indicate a potential transcriptomic compensatory response for the defective phagolysosomal acidification in *Pacc1^-/-^* macrophages (**Fig. 3B-G**). A subset of phagolysosomal processes were also dysregulated at baseline in *Pacc1^-/-^* compared to WT macrophages (***SI Appendix,* Fig. S13**), suggesting unexpected PACC1 activity in resting cells. In summary, these findings indicate that loss of *Pacc1* in macrophages results in transcriptomic dysregulation in immunological and phagolysosomal pathways, particularly in response to phagocytotic bacterial stimulus.

### Pacc1 knockout mice and myeloid cell-targeted conditional Pacc1 knockout mice are susceptible to E. coli sepsis

Because defective phagolysosomal acidification inhibits pathogen clearance (17, 18), we next studied if *Pacc1^-/-^* mice were susceptible to bacterial infections. In an established *E. coli* sepsis model (11) (**Fig. 5A**), *Pacc1^-/-^*mice succumbed to severe mortality (**Fig. 5B**). In contrast to 100% of WT counterparts, only 50% of *Pacc1^-/-^* mice survived. This increased lethality was associated with significantly higher bacterial burden in peritoneal lavage (**Fig. 5C**) and systemic bacteremia (**Fig. 5D**, gating scheme in ***SI Appendix,* Fig. S14A**). Moreover, a greater inflammatory response in *Pacc1^-/-^* mice was indicated by increased myeloid cell infiltration (**Fig. 5E**, gating scheme in ***SI Appendix,* Fig. S14B**) and IL-6 release in peritoneal lavage fluid (**Fig. 5F**). However, this enhanced compensatory response was apparently unable to clear infection. In conclusion, these data suggest *Pacc1^-/-^* mice are susceptible to bacterial infection and dysregulated inflammation, which may be explained by defective myeloid cell responses.

**Fig. 5.**
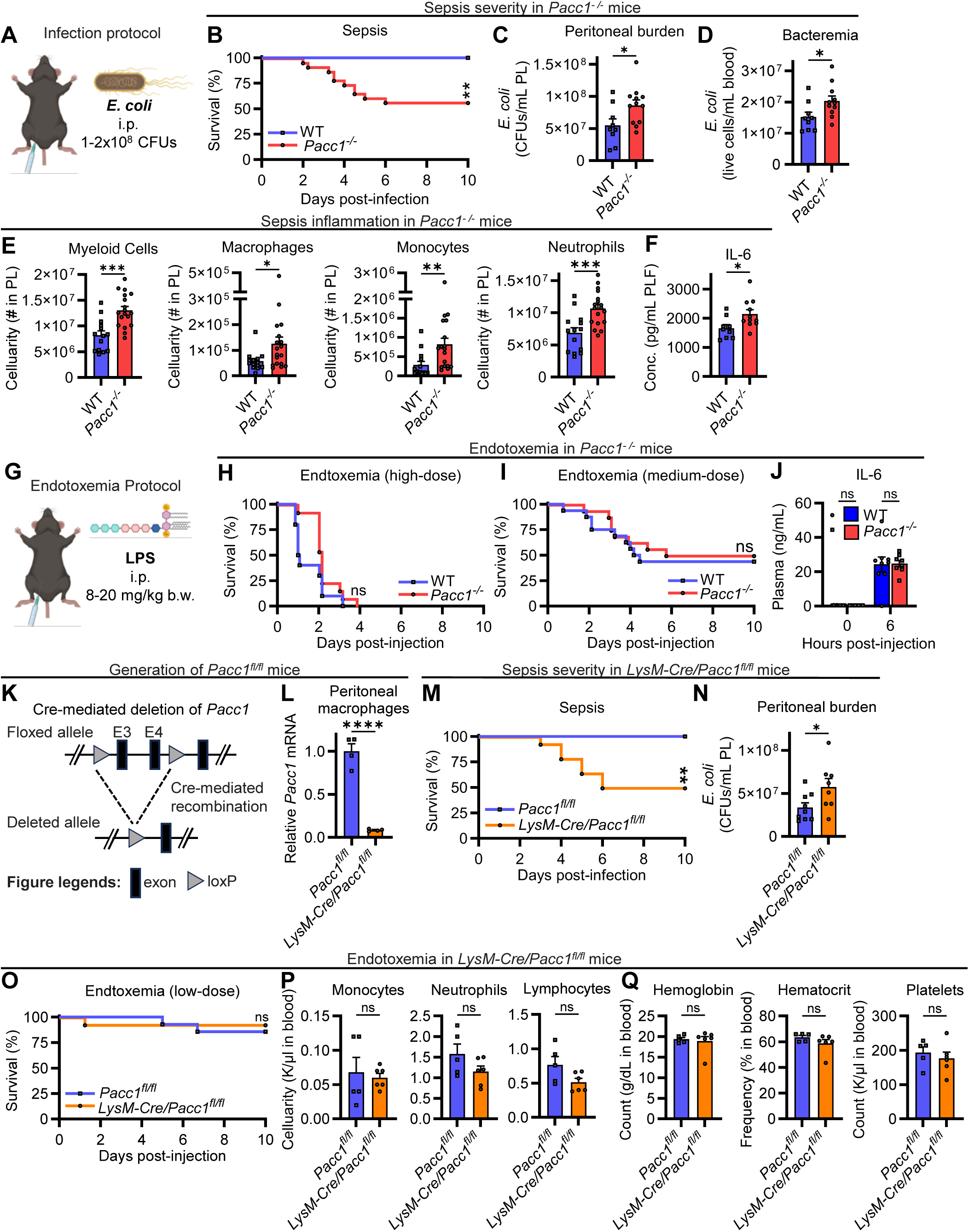
Global and myeloid cell conditional *Pacc1* deletion confers susceptibility to *E. coli* sepsis. (**A**) Schematic showing sepsis infection. Wildtype (WT) and *Pacc1^-/-^*mice were given 1-2×10^8^ colony-forming units (CFUs) of *Escherichia (E). coli* intraperitoneally (i.p.). Peritoneal lavage (PL) and blood were collected after 6 or 24 h, or survival was monitored, as indicated. (**B**) Sepsis survival (n=17-22 mice/group, pooled from 3 independent experiments). (**C**) CFUs in PL after 6 h assayed by serially diluted plated cultures. (**D**) Number of live bacteria in blood after 24 h by flow cytometry. (**E**) Myeloid cell numbers in PL after 24 h by flow cytometry. (**F**) IL-6 in PL fluid (PLF) after 6 h by ELISA. (**G**) Schematic showing endotoxemia induction. Mice were given *E. coli*-derived lipopolysaccharide (LPS) i.p. to induce phagocytosis-independent endotoxemia. (**H**) Endotoxemia survival with high-dose (20 mg/kg body weight [b.w.]) LPS (n=10-13 mice/group) and (**I**) medium-dose (10 mg/kg b.w.) LPS (n=16 mice/group, 2 pooled experiments). (**J**) IL-6 in plasma after low-dose endotoxemia as in panel H by ELISA. (**K**) Schematic showing generation of *Pacc1*-floxed mice and Cre-mediated deletion. In brief, LoxP sites flanking exons 3/4 of the *Pacc1* were inserted using CRISPR/Cas9 in WT mice, such that their Cre-mediated deletion eliminates productive *Pacc1* expression. (**L**) *Pacc1* expression loss in live CD11b^+^F4/80^+^ peritoneal macrophages from *LysM-Cre/Pacc1^fl/fl^* mice by qRT-PCR. *Pacc1*-floxed (*Pacc1^fl/fl^*) and *LysM-Cre* mice were crossed for targeted deletion of *Pacc1* in myeloid cells, and macrophages were FACSorted from the PL of naïve mice. (**M**) Sepsis survival following *E. coli* infection as in A (n=11-14 mice/group, 2 pooled experiments). (**N**) CFUs in PL after 6 h by serially diluted plated cultures. (**O**) Endotoxemia survival with low-dose (8 mg/kg b.w.) LPS (n=14 mice/group). Parameters determined by blood analyzer in mice with low-dose endotoxemia after 24 h, including (**P**) immune cell counts and (**Q**) blood parameters. Data represent ≥2 independent experiments and show one symbol per mouse, mean±SEM, *P<0.05, **P<0.01, ***P<0.001, ns=not significant, Student’s t-test (**C**-**F, J**, **L**, **N**, **P**, **Q**), or Mantel-Cox log-rank test (**B**, **H**, **I**, **M**, **O**).

Next, to test whether this susceptibility of *Pacc1^-/-^*mice to a gram-negative bacterial infection in the peritoneum (**Fig. 5A-F**) was broadly applicable, we examined susceptibility in a gram-positive pneumonia setting (56). Mice were intranasally (i.n.) inoculated with S. *pneumoniae* (***SI Appendix,* Fig. S15A**). As with *E. coli* sepsis, *Pacc1^-/-^* mice experienced significantly higher mortality (***SI Appendix,* Fig. S15B**), greater peak weight loss (***SI Appendix,* Fig. S15C**), and more lung injury as indicated by bronchoalveolar protein leakage (***SI Appendix,* Fig. S15D**). Accordingly, *Pacc1^-/-^* mice tended to have an elevated bacterial burden (***SI Appendix,* Fig. S15E**). Like *E. coli* sepsis, pneumococcal pneumonia induced more pronounced local myeloid cell infiltration in bronchoalveolar lavages from *Pacc1^-/-^* mice (***SI Appendix,* Fig. S15F**, gating scheme in **Fig. S16**). This was accompanied by a general elevation of inflammatory mediators (***SI Appendix,* Fig. S15G-H**), including IL-6 as well as IFN-γ, a cytokine important for macrophage activation and bacterial clearance (5, 15, 57). In contrast, we found no signs of pulmonal inflammation at baseline in naïve *Pacc1^-/-^* mice (***SI Appendix,* Fig. S17**), which was consistent with immunoprofiling in other sites (***SI Appendix,* Fig. S3J-L, S7**). We noted an elevation in homeostatic numbers of alveolar macrophages in *Pacc1^-/-^* mice (***SI Appendix,* Fig. S17A-B**), which could represent a potential compensatory response for their phagocytic defects. Like WT, naive *Pacc1^-/-^*mouse lungs displayed low numbers of infiltrating monocytes and neutrophils (***SI Appendix,* Fig. S17A-B**), and undetectable levels of IL-6 (***SI Appendix,* Fig. S17C**). Overall, these data suggest *Pacc1^-/-^* mice are susceptible to bacterial infection and trigger hyperinflammation associated with defective myeloid cells in this context.

To further explore if severe mortality and inflammation in septic *Pacc1^-/-^* mice was specifically dependent on phagocytosis, we induced sterile endotoxemia by i.p. injection of *E. coli*-derived LPS (**Fig. 5G**). In contrast to live *E. coli*, LPS activates TLR4 without phagocytic uptake (12). High-dose LPS resulted in 100% mortality (**Fig. 5H**), while medium-dose LPS led to approximately 50% mortality (**Fig. 5I**). Survival curves for *Pacc1^-/-^* and WT mice were identical at both titrations, indicating PACC1 is not essential for phagocytosis-independent inflammatory responses. Accordingly, IL-6 concentrations were comparable between strains (**Fig. 5J**). Transcriptional analysis of *Pacc1^-/-^* BMDMs following LPS stimulation *in vitro* also confirmed responses were indistinguishable from WT (***SI Appendix,* Fig. S18**).

Although *PACC1* is enriched in macrophages (**Fig. 1-2**), its expression remains ubiquitous (**Fig. 1**), challenging the assignment of a causal role for PACC1 in macrophages during bacterial sepsis. To evaluate PACC1 loss in specific cell types during *in vivo* infection, we initiated the *de novo* generation of *Pacc1-floxed ^(fl/fl)^* mice via CRISPR/Cas9 (**Fig. 5K**). The expected elongated bands in *Pacc1^fl/fl^* mice that received loxP insertions were detectable by PCR (***SI Appendix,* Fig. S17A**). As expected, *Pacc1^fl/fl^* mice showed unchanged *Pacc1* expression in peritoneal cells before Cre-mediated recombination (***SI Appendix,* Fig. S17B**). *Pacc1^fl/fl^* mice crossed with *LysM-Cre* mice (58) for myeloid cell-targeted deletion showed >90% deletion in peritoneal tissue macrophages (**Fig. 5L**, gating scheme in ***SI Appendix,* Fig. S17C**). Crucially, these conditional knockout mice recapitulated the high sepsis mortality in global *Pacc1^-/-^* (**Fig. 5M**), which was similarly associated with greater peritoneal bacterial burden (**Fig. 5N**). Furthermore, in low-dose endotoxemia to detect potentially subtle sensitivity differences, myeloid cell-targeted *Pacc1* deletion did not impact survival (**Fig. 5O**) or peripheral blood and immune cell parameters (**Fig. 5P-Q**), further implicating PACC1 in phagocytosis-mediated immunity. In conclusion, PACC1 promotes protective host responses, as PACC1 deficiency within myeloid cells undermines anti-bacterial defense mechanisms and weakens resistance to sepsis and pneumonia.

## Discussion

Bacterial sepsis remains a major health burden (1). Our study has identified the recently discovered acid-sensitive chloride channel, PACC1 (also known as PAC, ASOR, TMEM206) (21, 22), as a regulator of host defense against bacterial infections and sepsis. PACC1 was highly expressed within human and mouse phagocytic myeloid cells, particularly macrophages, where it controlled sepsis outcomes. Using *de novo*-generated mice, *Pacc1* knockout globally or in myeloid cells impaired development of phagolysosomal acidification and conferred susceptibility to bacterial infection and excessive inflammation.

While PACC1 is uniquely activated under acidic conditions, its ubiquitous expression has obscured its physiological roles (21, 22). The data we have presented showing elevated *PACC1* in human and murine macrophages, along with GWAS data linking *PACC1* variants with myeloid cell phenotypes, suggest PACC1 is relevant for macrophage responses. Our re-analysis of proteomic data from Li *et al.* (51) confirmed PACC1 localized to bacteria-containing phagosomes in primary macrophages, although the lack of reliable, commercial monoclonal PACC1 antibodies constrained further experimental protein detection (47). These data are consistent with other systems using in-house generated PACC1 antibodies or reporters that show PACC1 colocalizes strongly with early endosomal and partially with late endosomal/lysosomal markers in BMDMs and cell lines (32, 34, 35). Recently, PACC1 expression has been reported in osteoclasts, a specialized multi-nucleated macrophage (32). Our findings explicitly identified elevated *Pacc1* in murine tissue macrophages and other primary immune cells *in vivo.* Our data also showed *Pacc1* expression in macrophages was dynamically regulated by diverse inflammatory stimuli. This suggests context-specific roles for PACC1 in infection that adapt to local milieus. For example, PACC1 downregulation at the plasma membrane may promote cell survival against extracellular acidic environments during inflammation/infection (21, 22, 30, 31).

Our view that PACC1 is essential for microbial phagocytosis and related endo/exo/pinocytic processes is shared by others (32, 34–37, 59). However, the contextual outcomes of *Pacc1* loss may depend on specific cell types, cargos, and other factors. For example, while PACC1 may limit endosomal acidification during transferrin recycling in cell lines and receptor trafficking in neurons (34, 36), we found PACC1 facilitates acidification in macrophage phagolysosomes. This may occur via the influx of chloride which acts as a negative electrical shunt to permit proton pumping that drives acidification (20, 33). While PACC1 is typically outwardly rectifying, it is a passive channel, and thus chloride transit depends on local electrochemical gradients (21–29). Of note, the chloride anion shunt hypothesis remains debated (20). Studies have suggested chloride is dispensable for lysosomal acidification (60, 61) and serves functions beyond shunting (20). There remains ambiguity around even the most well-studied chloride channel, cystic fibrosis transmembrane conductance regulator (CFTR). Multiple groups have reported CFTR is required for intracellular acidification, while others have shown no effect on acidification and macrophage bacterial clearance (20, 33, 62, 63). Posited explanations include differences in CFTR knockout, inhibition, or mutations; differences in human and mouse primary cells and cell lines; and differences in protocols and models (20, 33, 62, 63). Most recently, Badr *et al*. suggested bacterial cargo is critical, as CF patient macrophages display defective lysosomal acidification following phagocytosis of *Burkholderia cenocepacia*, but not *E. coli* (64). Further work is needed to clarify the nuanced context-dependent roles of chloride in vesicular acidification, as well as the specific biophysical mechanisms undermining the acidifying phagolysosome in *Pacc1^-/-^*macrophages.

In the present study, transcriptomic profiling of *Pacc1^-/-^*macrophages exposed to inactivated *E. coli* revealed dysregulated immunologic functions. Consistent with our hypothesis that PACC1 is critical for phagolysosomal activity, lysosomal processes were highly dysregulated in *Pacc1^-/-^* macrophages; while processes associated with early phagocytosis (e.g., engulfment) were largely unaffected. The observed upregulation of lysosomal processes in *Pacc1^-/-^* macrophages was unexpected. In contrast, we observed defective phagolysosomal activity using acid-sensitive fluorescent *E. coli* BioParticles, and corresponding host defense impairments against live bacteria. It is possible transcriptomic upregulation of lysosomal processes represents a compensatory response. *Pacc1^-/-^* macrophages also displayed enhanced signaling from endolysosomal TLRs that respond to phagocytosed bacteria (17, 18). Impaired acidification may have caused inefficient cargo degradation which extended ligand–receptor contact. The “extended contact” hypothesis (65) acknowledges that while some acidification is required for endolysosomal TLR activation (66), partial defects can enhance signaling (65). Innate immune activation in *Pacc1^-/-^* macrophages was heterogenous, with both positive and negative regulation of various inflammatory markers, indicating PACC1 may differentially regulate inflammatory pathways via distinct mechanisms that deserve future study. Unexpectedly, we found *Pacc1^-/-^* macrophages displayed certain transcriptomic perturbations at baseline, despite the absence of overt abnormalities in naive mice (21, 22, 32). This may reflect homeostatic phago/efferocytosis (16–18), or possibly non-canonical roles for PACC1 (47, 67–74), as has been described for other chloride channels (75, 76).

Interestingly, we observed some differential inflammatory responses between *Pacc1^-/-^* vs. WT BMDMs challenged *in vitro* with inactivated *E. coli* compared to mice *in vivo* with live or inactivated *E. coli* (e.g. IL-6 production). This may be due to differences from tissue macrophage populations, differences from live bacterial burdens, as well as differences from the inflammatory milieu *in vivo* including infiltrating myeloid cells. *Pacc1* deficiency did not affect LPS-induced IL-6 production and low baseline IL-6 levels in mice. The present findings support that elevated IL-6 in *Pacc1⁻^/^⁻* mice challenged with bacteria reflects the particular *in vivo* inflammatory context rather than discrepant baseline macrophage-intrinsic responses.

The predominant impact of PACC1 during sepsis appeared dependent on phagocytosis. LPS, which activates TLR4 independent of phagocytosis (12), resulted in *Pacc1^-/-^* endotoxemia survival comparable to WT. Conversely, in line with impaired phagolysosomal acidification, *Pacc1^-/-^* mice exposed to live bacteria showed increased bacterial burdens and sepsis susceptibility. The augmented neutrophil recruitment in septic *Pacc1^-/-^* mice may have served as a compensatory response when tissue-resident macrophages failed to control bacterial pathogens during early infection (5, 15). Although heightened inflammation likely reflects increased bacterial burden, *Pacc1^-/-^* myeloid cells also displayed intrinsic hyperresponsiveness to bacterial challenge, as evidenced by exaggerated cytokine responses to inactivated *E. coli*. The translational relevance of these findings was supported by congruence with human genetic data. Predicted loss-of-function *PACC1* variants were associated with increased myeloid cell counts and risk for septicemia and various bacterial infections, reflecting the phenotype observed in *Pacc1^-/-^* mice during live infection. In our myeloid cell-targeted *LysM-Cre*/*Pacc1^fl/fl^*mice, sepsis susceptibility was recapitulated while endotoxemia outcomes remained unaltered. These findings further validated that PACC1 mediates phagolysosomal bacterial clearance via action within macrophages, monocytes, and neutrophils. Although PACC1 expression is greater in mononuclear phagocytes, including tissue macrophages that first encounter pathogens, recruited neutrophils become the predominant myeloid cell type (5, 16). *Pacc1^-/-^* neutrophils also exhibited defective development of the acidifying phagolysosome in response to *E. coli* BioParticles *in vivo*. Future studies should elaborate on the direct role of PACC1 in other PACC1-expressing immune cells. Of note, human pLoF *PACC1* variants were associated with increased myeloid cell counts and risk for septicemia and bacterial infections, supporting the translationally relevance of our findings. Unfortunately, only limited studies have been conducted in PACC1-overexpressing cell lines *in vitro* (21, 59, 77, 78), and there are no known specific PACC1 agonists to test if augmenting PACC1 is therapeutic during sepsis *in vivo* (77).

In parallel to our work, Cheng *et al.* also recently studied PACC1 in macrophages during infection (59). Both studies suggest PACC1 is crucial for phagolysosomal acidification, bacterial clearance, inflammation, and sepsis outcomes (59). Both indicate PACC1 is abundant in macrophages and downregulated by LPS, with *Pacc1^-/-^* macrophages showing increased interferon and other inflammatory signaling following bacterial challenge, though no change following LPS (59). Although Cheng *et al.* suggest enhanced acidification during phagocytosis releases more bacterial ligands to enhance immunologic signaling (59), partial defects in acidification may also enhance signaling by extending ligand contact (65).

While our study characterized defective phagolysosomal acidification, decreased survival, and hyperinflammation in *Pacc1^-/-^* mice following bacterial infection, Cheng *et al.* reported the converse (59). This divergence may arise from differences in mouse strains, pathogens, and methods. For example, we employed independently generated *Pacc1*-deficient strains with strict genetic/microbiome controls (59, 79, 80). We speculate evolutionary conserved PACC1 (21, 22) may be a target of microbial evasion strategies and pathogenicity factors that could modulate its activity, potentially explaining *E. coli* strain-dependent differences between studies (59). Our findings in both gram-negative peritoneal infection and gram-positive lung infection support relevance across bacterial contexts, as does congruence with human data linking pLoF *PACC1* variants with increased myeloid cell counts and sepsis risk. Discrepant results between studies may also stem from differences in cell types, time points, and cargo (64) (e.g., Cheng *et al*. used zymosan-conjugated particles (59), while we employed *E. coli*).

Most PACC1 functional studies have focused on direct electrophysiological conductance of chloride (21, 22, 26–28, 30, 34, 35, 81), though some suggest complex context-dependent and potentially non-canonical chloride-independent PACC1 functions (74). For example, recent work suggests PACC1 decreases acid-induced death in colorectal cancer cells (68, 69), diverging from findings by us and others (21, 22, 25, 30, 31). PACC1 may have direct protein binding/signaling partners (47, 67). PACC1 has been implicated in the opposing cancer processes of proliferation and migration (68–72, 82), and at non-acidic pH may mediate ophthalmic pathology (73). Interestingly, both we and Cheng *et al.* (59) identified in *Pacc1^-/-^* macrophages dysregulated RNA processing for which the role of chloride remains undefined. In total, findings from this and prior studies underscore the context-dependent and multifaceted functions of PACC1.

In conclusion, our results demonstrate a novel role for PACC1 in protective host defense and warrant further studies on PACC1 during sepsis and other bacterial infections.

## Materials and Methods

All the materials used in this study, including sources for reagents, and detailed methodologies for the generation of mouse strains, BMDM assays, analysis of splenocytes and blood, *E. coli* BioParticle assays, *E. coli* sepsis, pneumococcal pneumonia, endotoxemia, flow cytometry, protein detection, RT-qPCR, bulk and single-cell/nucleus transcriptomics, proteomics, and GWAS, and statistical analyses, are in the ***SI Appendix***, Materials and Methods.

## Acknowledgments

This work was supported by the Federal Ministry of Education and Research (01EO1503 to MB), the Deutsche Forschungsgemeinschaft (BO3482/3-3, BO3482/4-1 to MB), the National Institutes of Health (1R01HL141513, 1R01HL139641, 1R01HL166588 to MB; F31HL176388-01A1 to LG; 5T32HL007035-50 supported the training of LG). CR is a member at the German Center for Cardiovascular Research (DZHK), the DFG Research Unit 5644 INFINITE (RE 3450/15-1), the Center for Translational Vascular Biology, the Research Center for Immunotherapy (FZI), and Potentialbereich EXPOHEALTH, and received a Fellowship from the Gutenberg Research College. We cordially thank Dr. Thomas J. Jentsch for collaborative efforts and Dr. Kara Vasilew for assistance. We also thank the Flow Cytometry Core of Boston University School of Medicine (S10 OD038346) and Shari Brezinsky for assistance. The authors are responsible for the contents of this publication.

## Data, Materials, and Software Availability

Single-nucleus multiome sequencing data from our previous study is available in the NCBI Gene Expression Omnibus (GEO) repository (accession code: GSE269703; reviewer access token: wbcxyyaahfandwz). Only data pertaining to unstimulated cells were used here. Bulk RNA-seq data from this study is available in GEO (accession code: GSE292930; reviewer access token: yjinyuiihnyjpep). Phenotypes associated with human *PACC1* variants were from Open Targets (https://platform.opentargets.org/) (44). Genetic association data for *PACC1* variants in the All of Us Biobank were from Jurgens *et al.* browser (https://hugeamp.org:8000/research.html?pageid=600_traits_app_home) (45) and the UK Biobank from Genebass (https://app.genebass.org/) (46). *PACC1* expression from human single-cell and bulk RNA-seq data were from the HPA (https://www.proteinatlas.org/) (38) and Immunological Genome Project ICA (http://immunecellatlas.net/ICA_Skyline.php) (39). Data from published studies (11, 49, 51) were from respective source/supplementary data. This paper reports no original code. Additional information is available from the corresponding author upon request.

## References

1. K. E. Rudd et al., Global, regional, and national sepsis incidence and mortality, 1990-2017: analysis for the Global Burden of Disease Study. Lancet 395, 200–211 (2020).

2. C. J. Paoli, M. A. Reynolds, M. Sinha, M. Gitlin, E. Crouser, Epidemiology and Costs of Sepsis in the United States-An Analysis Based on Timing of Diagnosis and Severity Level. Crit Care Med 46, 1889–1897 (2018).

3. C. M. Torio, B. J. Moore, “National Inpatient Hospital Costs: The Most Expensive Conditions by Payer, 2013” in Healthcare Cost and Utilization Project (HCUP) Statistical Briefs. (Agency for Healthcare Research and Quality (US), Rockville (MD), 2006).

4. C. Rhee et al., Prevalence, Underlying Causes, and Preventability of Sepsis-Associated Mortality in US Acute Care Hospitals. JAMA Netw Open 2, e187571 (2019).

5. L. J. Quinton, A. J. Walkey, J. P. Mizgerd, Integrative Physiology of Pneumonia. Physiol Rev 98, 1417–1464 (2018).

6. Anonymous, Global burden of bacterial antimicrobial resistance 1990-2021: a systematic analysis with forecasts to 2050. Lancet 10.1016/s0140-6736(24)01867-1 (2024).

7. E. Karakike et al., Coronavirus Disease 2019 as Cause of Viral Sepsis: A Systematic Review and Meta-Analysis. Crit Care Med 49, 2042–2057 (2021).

8. M. Bonten et al., Epidemiology of Escherichia coli Bacteremia: A Systematic Literature Review. Clin Infect Dis 72, 1211–1219 (2021).

9. Anonymous, Global mortality associated with 33 bacterial pathogens in 2019: a systematic analysis for the Global Burden of Disease Study 2019. Lancet 400, 2221–2248 (2022).

10. M. Bosmann et al., Interruption of macrophage-derived IL-27(p28) production by IL-10 during sepsis requires STAT3 but not SOCS3. J Immunol 193, 5668–5677 (2014).

11. J. Roewe et al., Bacterial polyphosphates interfere with the innate host defense to infection. Nat Commun 11, 4035 (2020).

12. M. Bosmann, N. F. Russkamp, P. A. Ward, Fingerprinting of the TLR4-induced acute inflammatory response. Exp Mol Pathol 93, 319–323 (2012).

13. M. Bosmann et al., CD11c+ alveolar macrophages are a source of IL-23 during lipopolysaccharide-induced acute lung injury. Shock 39, 447–452 (2013).

14. S. H. E. Kaufmann, A. Dorhoi, R. S. Hotchkiss, R. Bartenschlager, Host-directed therapies for bacterial and viral infections. Nat Rev Drug Discov 17, 35–56 (2018).

15. M. Bosmann, P. A. Ward, The inflammatory response in sepsis. Trends Immunol 34, 129–136 (2013).

16. E. Mass, F. Nimmerjahn, K. Kierdorf, A. Schlitzer, Tissue-specific macrophages: how they develop and choreograph tissue biology. Nature Reviews Immunology 23, 563–579 (2023).

17. A. M. Pauwels, M. Trost, R. Beyaert, E. Hoffmann, Patterns, Receptors, and Signals: Regulation of Phagosome Maturation. Trends Immunol 38, 407–422 (2017).

18. E. Uribe-Querol, C. Rosales, Phagocytosis: Our Current Understanding of a Universal Biological Process. Front Immunol 11, 1066 (2020).

19. T. J. Jentsch, M. Pusch, CLC Chloride Channels and Transporters: Structure, Function, Physiology, and Disease. Physiol Rev 98, 1493–1590 (2018).

20. T. Stauber, T. J. Jentsch, Chloride in vesicular trafficking and function. Annu Rev Physiol 75, 453–477 (2013).

21. J. Yang et al., PAC, an evolutionarily conserved membrane protein, is a proton-activated chloride channel. Science 364, 395–399 (2019).

22. F. Ullrich et al., Identification of TMEM206 proteins as pore of PAORAC/ASOR acid-sensitive chloride channels. Elife 8 (2019).

23. S. Lambert, J. Oberwinkler, Characterization of a proton-activated, outwardly rectifying anion channel. J Physiol 567, 191–213 (2005).

24. C. Auzanneau, V. Thoreau, A. Kitzis, F. Becq, A Novel voltage-dependent chloride current activated by extracellular acidic pH in cultured rat Sertoli cells. J Biol Chem 278, 19230–19236 (2003).

25. H. Y. Wang, T. Shimizu, T. Numata, Y. Okada, Role of acid-sensitive outwardly rectifying anion channels in acidosis-induced cell death in human epithelial cells. Pflugers Arch 454, 223–233 (2007).

26. Z. Ruan, J. Osei-Owusu, J. Du, Z. Qiu, W. Lü, Structures and pH-sensing mechanism of the proton-activated chloride channel. Nature 588, 350–354 (2020).

27. J. Osei-Owusu et al., Molecular determinants of pH sensing in the proton-activated chloride channel. Proc Natl Acad Sci U S A 119, e2200727119 (2022).

28. C. Wang, M. M. Polovitskaya, B. D. Delgado, T. J. Jentsch, S. B. Long, Gating choreography and mechanism of the human proton-activated chloride channel ASOR. Sci Adv 8, eabm3942 (2022).

29. Z. Deng et al., Cryo-EM structure of a proton-activated chloride channel TMEM206. Sci Adv 7 (2021).

30. J. Osei-Owusu, J. Yang, M. Del Carmen Vitery, M. Tian, Z. Qiu, PAC proton-activated chloride channel contributes to acid-induced cell death in primary rat cortical neurons. Channels (Austin) 14, 53–58 (2020).

31. X. M. Zha, Z. G. Xiong, R. P. Simon, pH and proton-sensitive receptors in brain ischemia. J Cereb Blood Flow Metab 42, 1349–1363 (2022).

32. P. Xue et al., Proton-Activated Chloride Channel Increases Endplate Porosity and Pain in a Mouse Spinal Degeneration Model. J Clin Invest 10.1172/jci168155 (2024).

33. J. E. DiCiccio, B. E. Steinberg, Lysosomal pH and analysis of the counter ion pathways that support acidification. J Gen Physiol 137, 385–390 (2011).

34. J. Osei-Owusu et al., Proton-activated chloride channel PAC regulates endosomal acidification and transferrin receptor-mediated endocytosis. Cell Rep 34, 108683 (2021).

35. M. Zeziulia, S. Blin, F. W. Schmitt, M. Lehmann, T. J. Jentsch, Proton-gated anion transport governs macropinosome shrinkage. Nat Cell Biol 24, 885–895 (2022).

36. K. H. Chen et al., Loss of the proton-activated chloride channel in neurons impairs AMPA receptor endocytosis and LTD via endosomal hyper-acidification. Cell Rep 44, 115302 (2025).

37. N. Koylass et al., The Proton-Activated Chloride Channel Inhibits SARS-CoV-2 Spike Protein-Mediated Viral Entry Through the Endosomal Pathway. J Cell Physiol 240, e70063 (2025).

38. M. Karlsson et al., A single-cell type transcriptomics map of human tissues. Further details available at: https://www.proteinatlas.org/about/assays+annotation#normalization_rna. Sci Adv 7 (2021).

39. T. S. Heng, M. W. Painter, The Immunological Genome Project: networks of gene expression in immune cells. Nat Immunol 9, 1091–1094 (2008).

40. Y. J. Jung, W. S. Chung, Phagocytic Roles of Glial Cells in Healthy and Diseased Brains. Biomol Ther (Seoul) 26, 350–357 (2018).

41. A. J. Byrne, S. A. Mathie, L. G. Gregory, C. M. Lloyd, Pulmonary macrophages: key players in the innate defence of the airways. Thorax 70, 1189–1196 (2015).

42. A. Madhvi, H. Mishra, G. R. Leisching, P. Z. Mahlobo, B. Baker, Comparison of human monocyte derived macrophages and THP1-like macrophages as in vitro models for M. tuberculosis infection. Comp Immunol Microbiol Infect Dis 67, 101355 (2019).

43. C. Klijn et al., A comprehensive transcriptional portrait of human cancer cell lines. Nat Biotechnol 33, 306–312 (2015).

44. D. Ochoa et al., The next-generation Open Targets Platform: reimagined, redesigned, rebuilt. Nucleic Acids Res 51, D1353–d1359 (2023).

45. S. J. Jurgens et al., Rare coding variant analysis for human diseases across biobanks and ancestries. Nat Genet 56, 1811–1820 (2024).

46. C. Bycroft et al., The UK Biobank resource with deep phenotyping and genomic data. Nature 562, 203–209 (2018).

47. S. F. Pastore et al., Biallelic inheritance in a single Pakistani family with intellectual disability implicates new candidate gene RDH14. Sci Rep 11, 23113 (2021).

48. C. Lai, D. Stepniak, L. Sias, C. Funatake, A sensitive flow cytometric method for multi-parametric analysis of microRNA, messenger RNA and protein in single cells. Methods 134-135, 136–148 (2018).

49. J. Qie et al., Integrated proteomic and transcriptomic landscape of macrophages in mouse tissues. Nature Communications 13, 7389 (2022).

50. S. Lykke-Andersen, T. H. Jensen, Nonsense-mediated mRNA decay: an intricate machinery that shapes transcriptomes. Nature Reviews Molecular Cell Biology 16, 665–677 (2015).

51. K. Li et al., Profiling phagosome proteins identifies PD-L1 as a fungal-binding receptor. Nature 630, 736–743 (2024).

52. A. Neaga, J. Lefor, K. E. Lich, S. F. Liparoto, Y. Q. Xiao, Development and validation of a flow cytometric method to evaluate phagocytosis of pHrodo™ BioParticles® by granulocytes in multiple species. J Immunol Methods 390, 9–17 (2013).

53. B. Lindner, T. Burkard, M. Schuler, Phagocytosis assays with different pH-sensitive fluorescent particles and various readouts. Biotechniques 68, 245–250 (2020).

54. C. Lambert et al., Cytochalasans and Their Impact on Actin Filament Remodeling. Biomolecules 13 (2023).

55. I. Rusinova et al., Interferome v2.0: an updated database of annotated interferon-regulated genes. Nucleic Acids Res 41, D1040–1046 (2013).

56. M. Dudek et al., Lung epithelium and myeloid cells cooperate to clear acute pneumococcal infection. Mucosal Immunol 9, 1288–1302 (2016).

57. J. C. Gomez et al., Mechanisms of interferon-γ production by neutrophils and its function during Streptococcus pneumoniae pneumonia. Am J Respir Cell Mol Biol 52, 349–364 (2015).

58. C. L. Abram, G. L. Roberge, Y. Hu, C. A. Lowell, Comparative analysis of the efficiency and specificity of myeloid-Cre deleting strains using ROSA-EYFP reporter mice. J Immunol Methods 408, 89–100 (2014).

59. H. Y. Cheng et al., Proton-activated chloride channel governs phagosome-mediated antibacterial immunity in peritoneal macrophages. J Exp Med 222 (2025).

60. B. E. Steinberg et al., A cation counterflux supports lysosomal acidification. J Cell Biol 189, 1171–1186 (2010).

61. S. Weinert et al., Uncoupling endosomal CLC chloride/proton exchange causes severe neurodegeneration. Embo j 39, e103358 (2020).

62. A. Lukasiak, M. Zajac, The Distribution and Role of the CFTR Protein in the Intracellular Compartments. Membranes 11, 804 (2021).

63. A. Meoli et al., Impact of CFTR Modulators on the Impaired Function of Phagocytes in Cystic Fibrosis Lung Disease. Int J Mol Sci 23 (2022).

64. A. Badr et al., CFTR Modulators Restore Acidification of Autophago-Lysosomes and Bacterial Clearance in Cystic Fibrosis Macrophages. Front Cell Infect Microbiol 12, 819554 (2022).

65. W. McAlpine et al., Excessive endosomal TLR signaling causes inflammatory disease in mice with defective SMCR8-WDR41-C9ORF72 complex function. Proc Natl Acad Sci U S A 115, E11523–e11531 (2018).

66. O. de Bouteiller et al., Recognition of double-stranded RNA by human toll-like receptor 3 and downstream receptor signaling requires multimerization and an acidic pH. J Biol Chem 280, 38133–38145 (2005).

67. J. H. Zhang, Y. Qin, S. C. Sun, TMEM206 gene knockout improves balance performance in SCA1 transgenic mice. IBRO Neurosci Rep 19, 996–999 (2025).

68. S. Kappel et al., CBA (4-chloro-2-(2-chlorophenoxy)acetamido) benzoic acid) inhibits TMEM206 mediated currents and TMEM206 does not contribute to acid-induced cell death in colorectal cancer cells. Front Pharmacol 15, 1369513 (2024).

69. K. Melek, B. Hauert, S. Kappel, TMEM206 Contributes to Cancer Hallmark Functions in Colorectal Cancer Cells and Is Regulated by p53 in a p21-Dependent Manner. Cells 13 (2024).

70. F. Peng, H. Li, J. Li, Z. Wang, Downregulation of the Proton-Activated Cl- Channel TMEM206 Inhibits Malignant Properties of Human Osteosarcoma Cells. Oxid Med Cell Longev 2021, 3672112 (2021).

71. L. Song, D. Feng, J. Tan, H. Zhang, Effects of TMEM206 on the malignant behavior of HepG2 human hepatocellular carcinoma cells. European Journal of Inflammation 20, 1721727X221122724 (2022).

72. L. Zhang, S. Y. Liu, X. Yang, Y. Q. Wang, Y. X. Cheng, TMEM206 is a potential prognostic marker of hepatocellular carcinoma. Oncol Lett 20, 174 (2020).

73. Z. J. Yang, S. Y. Huang, Y. F. Zhou, S. C. Sun, Knockout of TMEM206 in mice associated with a loss of corneal transparency. Int J Ophthalmol 17, 1967–1972 (2024).

74. K. H. Chen, T. Hagino, Z. Qiu, The Proton-Activated Chloride Channel: Molecular Identification, Structure, and Role in Organelle Physiology. Annu Rev Physiol 10.1146/annurev-physiol-022724-105357 (2025).

75. S. Gururaja Rao, N. J. Patel, H. Singh, Intracellular Chloride Channels: Novel Biomarkers in Diseases. Front Physiol 11, 96 (2020).

76. S. Gururaja Rao, D. Ponnalagu, N. J. Patel, H. Singh, Three Decades of Chloride Intracellular Channel Proteins: From Organelle to Organ Physiology. Curr Protoc Pharmacol 80, 11.21.11–11.21.17 (2018).

77. L. Mihaljević, Z. Ruan, J. Osei-Owusu, W. Lü, Z. Qiu, Inhibition of the proton-activated chloride channel PAC by PIP(2). Elife 12 (2023).

78. H. Y. Cheng et al., The phagosome-mediated anti-bacterial immunity is governed by the proton-activated chloride channel in peritoneal macrophages. bioRxiv 10.1101/2025.02.27.640612, 2025.2002.2027.640612 (2025).

79. A. C. Ericsson, C. L. Franklin, The gut microbiome of laboratory mice: considerations and best practices for translational research. Mamm Genome 32, 239–250 (2021).

80. V. M. Ripoll et al., Nicotinamide nucleotide transhydrogenase (NNT) acts as a novel modulator of macrophage inflammatory responses. Faseb j 26, 3550–3562 (2012).

81. J. Osei-Owusu et al., Molecular mechanism underlying desensitization of the proton-activated chloride channel PAC. Elife 11 (2022).

82. B. Boilly, A. S. Vercoutter-Edouart, H. Hondermarck, V. Nurcombe, X. Le Bourhis, FGF signals for cell proliferation and migration through different pathways. Cytokine Growth Factor Rev 11, 295–302 (2000).

