## Supplementary material for "Proton-Activated Chloride Channel 1 (PACC1) is essential for innate host defense against bacterial sepsis": SI Appendix

**This PDF file includes:**

Materials and Methods
Figures S1 to S19
Tables S1 to S4
Legends for Datasets S1 to S4
SI References

**Other supporting materials for this manuscript include the following:**

Datasets S1 to S4

### Materials and Methods

*Mice*

All animal studies were approved by the State Investigation Office of Rhineland-Palatinate and the Institutional Animal Care and Use Committee (IACUC) of Boston University.

*Pacc1*$^{-/-}$ mice (C57BL/6J-*Pacc1*$^{em1Bosm}$/Bosm, MGI: 6283429) were generated *de novo* by CRISPR/Cas9-mediated gene editing at the Johannes Gutenberg University Mainz, Germany. The single guide (sg)RNAs targeting exon 2 were microinjected into wildtype (WT) C57BL/6J zygotes, which were subsequently cultured under sterile conditions for 24 hours before transfer to pseudopregnant WT C57BL/6J females. The resulting offspring were backcrossed for 5 generations to further mitigate potential off-target effects. Sanger sequencing and PCR confirmed a 120 base pair (bp) deletion spanning intron 1 and exon 2, and reverse transcription-quantitative PCR (RT-qPCR) indicated reduced *Pacc1* mRNA levels across multiple exons, likely due to nonsense-mediated decay (1). This deletion causes a frameshift resulting in a premature stop codon in exon 4.

*Pacc1*$^{fl/fl}$ mice (C57BL/6J-*Pacc1*$^{em2Bosm}$/Bosm, MGI: 7703212, JAX stock No.: 039760) were donated by us to the Jackson Laboratories (Bar Harbor, ME) after generation by Biocytogen (Beijing, China) using CRISPR/Cas9-based extreme genome editing (EGE™) technology. Based on a Universal CRISPR Activity Assay (UCA)™ activity detection system two sgRNAs targeting the intron 2-3 (5'-TCCTTGCTAAGCACCCGACTTGG-3') and intron 4-5 (5'-GGACCCTTATACACTGTGCACGG-3') were selected to mediate the insertion of two loxP sites flanking a ~2 kb chromosomal region (exons 3-4) at the *Pacc1* locus in the mouse genome. The targeting vector DNA, Cas9 mRNA, and two sgRNAs were microinjected into zygotes derived from WT C57BL/6J mice and then the embryos were transferred to surrogate mothers. F0 positive founders were bred with WT C57BL/6J mice to get F1 pups. F1 germline transmission heterozygous mice were verified by two rounds of PCR, DNA sequencing, and Southern blot with 5' and 3' end to exclude random insertions. Heterozygous mice were bred with each other to obtain homozygous mice. Homozygous *Pacc1*$^{fl/fl}$ mice were then crossed with *LysM-Cre*-expressing mice (B6.129P2-*Lyz2*$^{tm1(cre)Ifo}$/J). Following Cre-mediated recombination, the exons 3-4 were excised and the deletion led to a protein-reading frameshift, resulting in the generation of a tissue-specific knockout mouse model.

Genotyping primer sequences and guide RNAs are listed in ***SI Appendix,* Table S2**. The single guide RNAs used to generate *Pacc1*$^{-/-}$ and *Pacc1*$^{fl/fl}$ mice were selected based on *in silico* predictions using online tools such as Sanger WGE CRISPR Finder, CRISPOR, and CCTop. These tools were employed to identify sgRNAs with minimal predicted off-target activity (2-4). The few predicted off-target sites each contained at least three nucleotide mismatches, including within the seed region, and were located on other chromosomes, predominantly in non-coding regions. Based on prior validations and our experience, off-target effects are unlikely, as zygotic injection limits Cas9 activity to a narrow developmental window. Founder mice were also backcrossed for multiple generations to minimize any potential off-target variants.

All mice were housed under specific pathogen-free conditions with 45-60% humidity, an ambient temperature of 22±2°C, controlled 12-hour light/dark cycles, and *ad libitum* access to food and water. Female and male mice (8-16 weeks old) were age- and sex-matched for all experiments. WT C57BL/6J and *LysM-Cre* mice were originally obtained from Jackson Laboratories and subsequently bred at our institutions under co-housing conditions with *Pacc1* strains, allowing the generation and use of Cre-lox littermates for experimental comparisons.

*Isolation and culture of BMDMs*

Bone marrow-derived macrophages (BMDMs) were generated as previously described (5, 6). In brief, bone marrow was extracted and differentiated for seven days in L-929 (ATCC, Manassas, VA, USA) cell-conditioned medium prepared in-house in a 5% $CO_2$ humidified incubator at 37°C. Mature BMDMs were assessed by morphology using light microscopy and flow cytometry confirming >95% F4/80$^+$ surface expression. BMDMs were separated from nonadherent cells by washing once with sterile phosphate buffered saline (PBS) (Gibco, Life Technologies, Grand Island, NE, USA), and maintained in macrophage media: Roswell Park Memorial Institute medium (RPMI)-1640 (Gibco, Life Technologies) supplemented with 0.1% bovine serum albumin (BSA) (Sigma-Aldrich, St. Louis, MO, USA) and 100 U/mL penicillin-streptomycin (Thermo Fisher Scientific, Waltham, MA, USA). BMDMs were counted with a cell counter (CytoSMART, Skillman, NJ, USA), and 5x10$^5$ BMDMs per well were seeded and stimulated for the indicated time points with various stimuli corresponding to inflammatory or infectious insults, including lipopolysaccharide (LPS) (100 ng/mL, *E. coli* O111:B4, Sigma-Aldrich); interferon-β (IFN-β) (500 U/mL, Pbl Assay Science, Piscataway, NJ, USA); interferon-γ (IFN-γ) (100 ng/mL, Thermo Fisher Scientific/PeproTech); polyinosinic:polycytidylic acid (poly I:C) (1 μg/mL, InvivoGen, San Diego, CA, USA); zymosan (10 μg/mL, *Saccharomyces cerevisiae*, InvivoGen); peptidoglycan (PGN) (10 μg/mL, *E. coli* O111:B4, InvivoGen), double-stranded (ds) DNA (1 μg/mL, *E. coli* K12, InvivoGen), CpG oligodeoxynucleotides (ODN) (1 μM, InvivoGen), GpC ODN (1 μM, InvivoGen), or cyclic guanosine monophosphate–adenosine monophosphate (cGAMP) (10 μg/mL, InvivoGen). The dsDNA was transfected using oligofectamine (Invitrogen, Waltham, MA, USA). Alternatively, BMDMs were stimulated with inactivated *E. coli* BioParticles (Thermo Fisher Scientific) at a 1:10 dilution for the indicated time points.

*Phagocytic function and transcriptomic profiling of macrophages*

Phagosomal uptake by macrophages was determined using (acid-insensitive) Alexa Fluor (AF)488 *E. coli* BioParticles (Thermo Fisher Scientific). AF488-*E. coli* emit a stable signal from pH 3-10 independent of changes in phagolysosomal acidification (7). To assess uptake *in vitro*, macrophages (BMDMs, 5x$10^5$/well, >98% viability by Trypan Blue staining) were seeded in 24-well low-attachment plates and incubated with AF488 *E. coli* BioParticles ($10^6$/well) for 1 hour on a rocker to facilitate contact and uptake. Negative control BMDMs were either left untreated; stimulated with non-fluorescently conjugated fixed *E. coli* (MG1655); or pre-treated for 1 hour with 10 µM cytochalasin D, an inhibitor of actin polymerization and commonly used inhibitor of phagocytic uptake (7), before AF488 *E. coli* stimulation. Thereafter, macrophages were gently collected from the low-attachment surface and immediately placed on ice. Cells were fixed with 2% paraformaldehyde for 20 minutes on ice, stained for macrophage markers, and fluorescence was analyzed by flow cytometry.

Phagolysosomal acidification in activated phagocytes was determined using (acid-sensitive) pHrodo Green *E. coli* BioParticles (Thermo Fisher Scientific). These particles are phagocytosed and become fluorescent at low pH as the phagosome matures and fuses with the lysosome (8). To assess acidification *in vitro*, macrophages (BMDMs, 1x$10^5$/well, >98% viability by Trypan Blue staining) were seeded in 100 µL in a black-walled clear optical bottom 96-well cell culture plate and incubated overnight to permit cell adherence. 50 µL of the pHrodo *E. coli* suspension was added to a final concentration of 330 µg/mL. The fluorescence of the acid-sensitive pHrodo green dye was monitored for up to 4 hours at 37°C. Measurements were performed with excitation at 509 nm and emission at 533 nm, with a read height of 1mm, on a SpectraMax platereader (Molecular Devices, San Jose, CA, USA).

In addition, to assess acidification *in vivo*, 100 µg of these pHrodo Green *E. coli* BioParticles were injected intraperitoneally into mice in a volume of 100 µL. After 24 hours, the peritoneum was lavaged with 10 mL cold PBS containing 2% (v/v) heat-inactivated fetal bovine serum (FBS) with 0.5 mM ethylenediaminetetraacetic acid (EDTA) (Invitrogen, San Diego, USA), and the abdomen was gently agitated for 1 minute. Cells from collected peritoneal lavage (PL) were pelleted (650 g, 4°C,10 min), and cell numbers and pHrodo Green fluorescence were analyzed by flow cytometry. Cell-free PL fluid (PLF) was obtained by centrifuging the PL at 650 g, 4°C for 10 min to pellet mammalian cells. The resulting supernatant was centrifuged again at 2400 g, 4°C for 10 min to pellet bacteria, and the final supernatant was collected for protein quantification and stored at -80ºC. PL was also analyzed from naïve control mice without any treatment.

For transcriptomic experiments, WT and *Pacc1*$^{-/-}$ BMDMs from n=6 mice per genotype were generated and isolated as described in the above section. BMDMs (2x10$^{6}$/well, >98% viability by Trypan Blue staining) were seeded in 6-well plates in macrophage media and allowed to adhere for 2 hours. BMDMs were stimulated with *E. coli* BioParticles at 100 µg/mL for 12 hours, or left as unstimulated controls. RNA was isolated as described below and subjected to bulk RNA sequencing.

*Bacterial killing assay*

The pathogenic *E. coli* serotype O6:K2:H1 (ATCC 19138; a gift from Dr. Katrina Traber) was grown in LB medium (Thermo Fisher Scientific) to mid-log phase ($OD_{600}$ ~0.3). Macrophages (BMDMs, 5x10$^{5}$/well, >98% viability by Trypan Blue staining) were seeded in 12-well plates in macrophage media without antibiotic and allowed to adhere for 2 hours. BMDMs were infected at a multiplicity of infection (MOI) of 5 and incubated for 2 hours on a rocker to facilitate contact and uptake. Following infection, supernatants were collected and immediately placed on ice. Remaining adherent macrophages were washed twice with ice cold PBS and lysed in 0.2% Triton X-100 (Thermo Fisher Scientific) for 5 min on a rocker to release intracellular bacteria. Lysates and supernatants were serially diluted in PBS and plated on LB agar. Colony-forming units (CFUs) were enumerated after 24 hours at 37°C as an indicator of bacterial-killing capacity.

*Acid-induced cell death assay*

Macrophages (BMDMs, 5x10$^{5}$/well) were seeded in 24-well low-attachment plates and incubated for the indicated time points with either normal pH medium (pH 7.2) or acidic pH medium (pH 5.5). The pH 7.2 medium contained 25 mM 4-(2-hydroxyethyl)-1-piperazineethanesulfonic acid (HEPES) (Gibco, Thermo Fisher Scientific) (adjusted to pH 7.2), 0.1 g glucose, 320 mg NaCl, 20 mg KCl, 7 mg $CaCl_2$, 5 mg $MgSO_4$ (all from Sigma-Aldrich), and 10% FBS per 50 ml. The acidic medium (pH 5.5) was prepared using the same composition but adjusted to pH 5.5 with HEPES. After incubation, macrophages were gently collected from the low-attachment surfaces and stained with fixable viability dye along with macrophage markers. Samples were analyzed by flow cytometry to assess cell viability.

*Bulk RNA sequencing data analyses from de novo generated dataset*

WT and *Pacc1*$^{-/-}$ BMDMs stimulated with *E. coli* BioParticles, or without *E. coli* as unstimulated controls, were analyzed by bulk RNA-seq. All analyses were performed using Boston University's Shared Computing Cluster and were based on the raw fastq sequencing files from Novogene. After an initial quality check using FastQC (v0.11.9), adapter trimming was performed using trimgalore (v0.6.7; parameters: --2colour 20; --paired) with cutadapt (v4.1) through GNU parallel (2019-02-22) (9-12). A Snakemake (v7.17.1) workflow pipeline was used for alignment to reference mouse genome (GRCm39, Gencode vM36) using STAR aligner (v2.7.10b), sorting using samtools (v1.12), and quantification using the featureCounts module in subread (v2.0.3) (13-16). Multiqc (v1.12) was used to visualize the quality check metrics for the results from these analyses (17).

Exploratory and differential expression analyses were performed using the counts data in RStudioServer/R (2024.04.2+764/4.4.0) using DESeq2 (v1.46.0) (18). Gene annotations were retrieved using the biomaRt (v2.58.2/R4.3.2) package based on Ensembl (v113) (19). A combined term of strain (*Pacc1*$^{-/-}$ vs. WT) and stimulation status (with *E. coli* BioParticles vs. without *E. coli* BioParticles) was used in the DESeq2 design formula. After variance-stabilizing transformation of the counts, principal component analysis (PCA) plots were visualized using pcaExplorer (v3.0.0) and GraphPad Prism (20). Differential expression (DE) analysis was based on Wald's test with Benjamini-Hochberg multiple testing method with the threshold for statistical significance set at adjusted (adj.) $P<0.05$; shrinkage of large $\log_2$ fold change (FC) values was done the using apeglm (v1.28.0) package (20).

Gene set enrichment analysis (GSEA) was based on hallmark gene sets retrieved from the Molecular Signatures Database using the msigdbr package (v7.5.1) with the source species set to mouse (21-23). This enrichment analysis was performed using the clusterProfiler (v4.14.4) package with Benjamini-Hochberg multiple testing (24). Genes were ranked for GSEA based on the product of -$\log_{10}$(unadjusted p value) and sign($\log_2$ fold change); P values for genes that were very low (close to zero due to detection limits) were set to $10^{-1}$ x minimum of unadjusted *P* value to circumvent an undefined logarithmic conversion with zero values. The GSEA results were visualized using GraphPad Prism. The clusterProfiler package was also used for gene ontology biological process (GOBP) enrichment analyses based on significant DE genes with the background set to the list of all DE genes (24). The statistical significance threshold for the GOBP analysis was set at adj. $P<0.1$. Z-scores for the GOBPs were calculated using the GeneTonic (v3.0.0) package's wrapper for clusterProfiler results (25). The VennDetail (v1.22.0) package was used to assess common and unique DE genes between different groups (26). The heatmaps were visualized using ComplexHeatmap (v2.22.0) with code to visualize GOBPs alongside heatmaps from an online tutorial (27), and all other plots using ggplot2 (v3.5.1) packages and GraphPad Prism (v10.4.1) (28, 29). All the heatmap visualizations were based on significant DE genes with a baseMean >100. A list of lysosomal genes/enzymes were retrieved from the SinoBiological "Other Enzymes Product Center", and statistically significant DEGs encoding these genes/enzymes were visualized using heatmaps (30). Significant DE genes with baseMean >100 were also used for Ingenuity Pathway Analysis (IPA, QIAGEN Inc., https://digitalinsights.qiagen.com/IPA) with pathway significance threshold set at P<0.05 based on right-tailed Fisher's exact test (31). All other IPA parameters were set to the default, with species set to any, confidence as experimentally observed, and tissue/cell lines and data sources set to the default of all available sources. Interferon type I- and II-regulated genes were analyzed by querying the Interferome database (v2.01) (32) with the significant differentially expressed genes (adj. P<0.05). To select only highly regulated genes, a >10-fold change cutoff in the Interferome datasets was applied. The parameters for the search were set to the default criteria such as both *in vitro* or *in vivo* assays, any species, system, organ, and sample type. The results were visualized using an MA (log mean expression vs. $\log_2$ fold change) plot using ggplot2.

*Isolation and analysis of splenocytes and blood*

Spleens were isolated from mice and digested as previously described (33). In brief, spleens were injected with 1mL of pre-digestion mix (1 mg/mL collagenase D [Sigma-Aldrich], 20 µg/ml DNase I [Sigma-Aldrich], 2% FCS in HEPES-buffered RPMI-1640), minced with a razor blade, incubated at 37°C for 1 hour, pressed through a 100 µm cell strainer, and washed with PBS lacking $Ca^{2+}$ or $Mg^{2+}$ before proceeding to flow cytometry. Additionally, serum and plasma were collected from mice as previously described (6). At the Institute of Clinical Chemistry and Laboratory Medicine at the University of Mainz, clinical chemistry analysis (Alinity, Abbott, Germany) was performed from 150-200 µL lithium heparin plasma. Hematology analysis was performed with 200 µL EDTA blood prepared 1:3 with NaCl on an automated blood cell counter, and cholesterol was examined from 80 µL of serum. Complete blood counts following endotoxemia were acquired on a Drew Scientific HemaVet 950FS Auto Blood Analyzer at the Boston University Analytical Instrumentation Core.

*Bacterial sepsis with live Escherichia coli*

*Escherichia coli* (*E. coli*, MG1655) culture and peritoneal infection were performed as previously described (34). Briefly, an overnight culture of bacteria was inoculated into LB medium and cultured at

37°C with gentle shaking to an optical density of 0.5−0.6 at wavelength 600 nm by Nanodrop 2000c (Thermo Fisher Scientific) to achieve a logarithmic growth phase. After centrifugation (2400 g, 4°C for 10 min), the bacterial pellet was resuspended in sterile PBS without $Ca^{2+}$ or $Mg^{2+}$ and kept on ice during infection. The bacterial count of the inoculum was confirmed by plating serial dilutions of the inoculum on LB agar plates and extrapolation of CFU count after overnight incubation. Bacteria were stored at -80ºC in PBS aliquots.

On the day of infection, mice received a 100 μL dose of $2x10^8$ CFUs intraperitoneally. After 6 hours, peritoneal lavage was collected as described above, and bacterial load was assessed by streaking the PL on agar plates as described above. After 24 hours, systemic bacteremia was assessed by retro-orbital blood sampling and flow cytometry using the LIVE/DEAD BacLight Bacterial Viability Kit (Invitrogen). After 24 hours, PL was also collected to analyze immune cells by flow cytometry, and cell-free PLF was collected for protein quantification. Survival was monitored for 10 days.

*Bacterial pneumonia with Streptococcus pneumoniae*

*Streptococcus pneumoniae* (TIGR4 [JNR.7/87 or SPN4]) was cultured as previously described and inoculated into mice intranasally (35). Briefly, pneumococci were initially grown on 5% Columbia blood agar plates (BBL, BD Biosciences, Franklin Lakes, NJ, USA) overnight in a humidified atmosphere at 37°C in 5% $CO_2$. On the day of infection, cultured colonies were transferred in THY medium containing 3% Todd-Hewitt broth (BD Biosciences) and 0.5% yeast extract (Sigma-Aldrich) and grown to an optical density of 0.3 at 600 nm wavelength (~$1x10^8$ CFU/mL). After centrifugation (2400 g, 4°C for 10 min), the bacterial pellet was resuspended in sterile PBS and kept on ice during infection. The bacterial count of the inoculum was calculated by plating serial dilutions of the inoculum on blood agar plates and extrapolation of CFU count after overnight incubation. Bacteria were stored at -80ºC using Microbank™ beads (Pro-Lab Diagnostics, Richmond Hill, Canada) as single use aliquots.

On the day of infection, mice were anesthetized by intraperitoneal injection of ketamine (100 mg/kg body weight) and xylazine (8 mg/kg body weight), and infected intranasally with 20 μL of $2x10^6$ CFUs. For survival studies, mice were alternatively infected with $1x10^6$ CFUs. CFU load was confirmed after infection by plating inoculum blood agar plates and counting overnight CFUs. After 18 hours, the bronchoalveolar lavage (BAL) was collected using 6 x 1 mL aliquots of cold sterile PBS supplemented with 0.1% BSA (Sigma- Aldrich) and 2 mM EDTA by inserting an 18G angiocatheter (Exel International, Redondo Beach, CA, USA) into the trachea. Cell-free BAL fluid (BALF) was obtained by centrifuging the first 1mL of BAL at 650 g, 4°C for 10 min to pellet mammalian cells. The resulting supernatant was centrifuged again at 2400 g, 4°C for 10 min to pellet bacteria, and the final supernatant was collected for protein quantification and stored at -80ºC. Next, cells from all lavages were combined and centrifuged at 650 g, 4°C for 10 min and analyzed by flow cytometry. Bacterial load was assessed by plating BAL on blood agar plates and counting overnight CFUs. Cells were prepared for flow cytometric analysis. BAL was also analyzed from naïve control mice without any treatment.

#### *Endotoxemia*

Mice were injected i.p. with lipopolysaccharide (LPS) from *E. coli* (O111:B4, Sigma-Aldrich) to model endotoxemia as described previously (36). Mice were given either a low dose (8 mg/kg body weight) to model mild endotoxemia (resulted in approximately 10% lethality in WT mice), a medium dose (10 mg/kg body weight) to model moderate endotoxemia (resulting in approximately 50% lethality in WT mice), or a high dose (20 mg/kg body weight) to study severe endotoxemia (resulting in approximately 100% lethality in WT mice).

#### *Flow Cytometry*

Flow cytometry protocols for analysis of immune cells were used as previously reported (6, 37). BAL and PL samples were collected from mice as described above. Cell viability was confirmed via 4',6-diamidino-2-phenylindole (DAPI) or fixable viability dye (FVD-eFluor 780), as indicated in figure panels. Cell pellets were subjected to red blood cell lysis by resuspension in 200 µL ammonium-chloride-potassium (ACK) lysis buffer (Gibco) for 3 min on ice. Next, 800 µL ice-cold PBS was added to stop lysis followed by centrifugation (650 g, 4ºC for 5 min). For samples that received FVD eF780 (eBioscience), cells were resuspended in PBS and incubated with FVD (1:1000, 4°C, dark for 20 min). Cells were centrifuged (650 g, 4ºC for 5 min), washed twice with PBS, and incubated with TruStain fcX anti-CD16/CD32 Fc receptor block (BioLegend, San Diego, CA, USA) (1:100, 4ºC, dark for 20 min) followed by fluorophore-labelled mouse antibodies, as listed in ***SI Appendix,*** **Table S3**. All antibodies were diluted in fluorescence-activated cell sorting (FACS) buffer containing sterile PBS supplemented with 1% (w/v) BSA, 0.01% (w/v) sodium azide, and 2.5 mM EDTA. If cells were fixed prior to analysis, the stained cells were washed twice in FACS buffer and fixed in 2% paraformaldehyde (Santa Cruz Biotechnology, Dallas, TX, USA) (4ºC, dark for 20 min). Prepared cells were washed twice, centrifuged, and resuspended in 300 µL FACS buffer for acquisition. When needed, 12 µL DAPI (Life Technologies, Waltham, MA, USA) per sample was added for viability staining directly before measurements. For cell quantification on flow cytometers without precise volumetric control, samples received 20 µL of CountBright Absolute Counting Beads (Thermo Fisher Scientific). On flow cytometers with precise volumetric control, a fixed proportion of the sample was analyzed to count cells. Fluorescence minus one (FMO) controls were used to assist with gating strategies. For analysis of pHrodo Green *E. coli* BioParticle geometric mean fluorescence intensity (MFI) within myeloid cells following i.p. injection, baseline fluorescence of total live cells from an unstimulated WT mouse was used as a control. We have observed a similar MFI from unstimulated controls and controls treated with non-fluorescent fixed *E. coli* in this project (**Fig 3C**) and our previous work (6).

Flow cytometric acquisition of samples was performed on a BD Biosciences (Franklin Lakes, NJ, USA) LSR II, a BD FACS Canto flow cytometer with BD FACSDiva software, or a Cytek (Fremont, CA, USA) Aurora spectral flow cytometer using Cytek SpectroFlo software. Flow cytometric sorting of samples was performed on a BD FACSARIA II SORP cell sorter with BD FACSDiva software or BD FACSADiscover S8 spectral cell sorter. Preparation and analyses of final flow cytometric plots were performed in FlowJo (Ashland, OR, USA) version 10, or with Dotmatics cloud-based OMIQ software for spectral flow cytometric data.

Intracellular *Pacc1* mRNA expression was analyzed using the PrimeFlow RNA assay kit (Thermo Scientific) according to the manufacturer's instructions. Either WT samples which received "no *Pacc1* mRNA target probe condition," or "*Pacc1*$^{-/-}$ cells which still received *Pacc1* mRNA target probe" served as a negative control. The MFI of *Pacc1* was determined for each cell type relative to negative control.

*Protein Detection and ELISAs*

Cell-free BALF and PLF samples were diluted in PBS with 0.1% BSA to fit in standard range of commercially available ELISA kits and measured according to the manufacturer's protocol. IL-6 and IFN-γ were detected by ELISA kits from R&D Systems (Minneapolis, MN, USA) and BioLegend, respectively. The absorbance at 540 nm was measured with a Tecan Infinite M Nano plate reader (Tecan, Mannedorf, Switzerland), and concentration was determined from the standard curve generated by Magellan software v7.2 (Tecan). Other mediators were quantified using the Cytokines & Chemokines 26-Plex Mouse ProcartaPlex Panel 1 (Invitrogen) multiplex immunoassay at the Analytical Instrumentation Core at Boston University. BALF total protein leakage as a proxy for lung injury was measured with the Pierce bicinchoninic acid (BCA) Protein Assay Kit (Thermo Fisher Scientific).

*Reverse transcription quantitative PCR*

RNA isolation, reverse transcription and quantitative PCR were performed. In brief, total RNA was isolated from samples using the Qiagen (Hilden, Germany) RNeasy Plus Mini kit according to the manufacturer's instructions. RNA was quantified using a NanoDrop 2000c spectrophotometer (Thermo Fisher Scientific). The cDNA synthesis was performed using Maxima H Minus Reverse Transcriptase Kit (Thermo Fisher Scientific). 3-5 ng cDNA complemented with PowerUp™ SYBR™ Green Master Mix (Applied Biosystems, Thermo Fisher Scientific) and sequence-specific primers at a concentration of 500 nM each were used for qRT-PCR in a QuantStudio 3 Real-Time PCR instrument (Applied Biosystems, Thermo Fisher Scientific). To assay gene expression after LPS stimulation of macrophages *in vitro*, cDNA was amplified using sequence-specific primers and Fast Advanced Real-Time PCR Mix (Applied Biosystems, Thermo Fisher Scientific) on the QuantStudio Flex 7 Real-Time PCR system (Thermo Fisher Scientific). Gene expression was compared between samples by normalizing to *Gapdh* expression and applying the $2^{-\Delta\Delta Ct}$ formula. *Pacc1* primer sequences are provided in the ***SI Appendix,*** **Table S4**. Primers used to assay LPS-mediated transcription in macrophages *in vitro* were as follows: *Gapdh*, Mm99999915_g1; *Ccl2*, Mm00441242_m1, *Il1b*, Mm01336189_m1; *Il6*, Mm99999064_m1; *Il23*, Mm01160011_m1; *Tnfa* Mm00443258_m1; *Il12a*, Mm00434169_m1; Ifng, Mm01168134_m1; *Il10*, Mm01288386_m1; *Tgfb1*, Mm01178820_m1; *Il27*, Mm00461162_m1 (Applied Biosystems, Thermo Fisher Scientific).

*Single nucleus RNA sequencing and data analyses from de novo generated data*

*Pacc1* RNA expression in WT (C57BL/6) mice were based on single nucleus RNA/ATAC sequencing data in unstimulated peritoneal cells from another research study in our lab (manuscript under preparation; GSE269703). Peritoneal cells were FACSorted for 7-AAD$^-$CD45$^+$ cells and nuclei were isolated using the Chromium Single Cell Multiome ATAC and Gene Expression kit (10X Genomics) according to the manufacturer's instructions. Briefly, the sequencing data was aligned and quantified to the mouse reference genome (GRCm39, Gencode vM33) using 10X Genomics Cell Ranger ARC (v2.0.1). The filtered counts matrix generated was imported into RStudio Server running the R framework (v4.1.2) (38, 39) and analyzed using Seurat and Signac, with normalization using sctransform method and the standard integration workflow for multiome data (40). Cell types were identified based on markers from literature. *Pacc1* gene expression was plotted using ggplot2 (v3.3.5) in RStudio/R (v4.1.2) (41).

*Genetic association analyses from open-source databases*

The phenotypes associated with human *PACC1* variants were retrieved from the Open Targets Platform (42, 43). The platform integrates evidence from multiple sources such as OT Genetics (44), the ClinVar (45-48) human variation database, and genetic association data (gene burden) from sources such as Genebass, FinnGen, and the AstraZeneca PheWAS portal (49-58), and assigns a phenotype-gene association score for each source, and a cumulative association score. The OpenTargets association score is a quantitative value that represents the likelihood that a gene is linked to a particular phenotype, with higher scores indicating stronger evidence of association.

Genetic association data for *PACC1* variants within the All of Us Biobank was retrieved from the Broad CVDI human disease portal repository (59). We filtered the data in the portal to retrieve the summary statistics for *PACC1* gene-level associations for septicemia (phecode 38.0) in individuals with mixed (either non-European or European) ancestry. The beta statistic, standard error, and P-value for the low-frequency (minor allele frequency, MAF<1%: includes rare and ultra-rare) predicted loss-of-function (pLoF) variants were used for estimation of the odds ratio and 95% confidence interval, and were visualized through a forest plot using Graph Pad Prism v10.

The Genebass web portal (58, 60), developed using the UK Biobank (61) exome sequence data, was queried to retrieve aggregated gene-level association results (based on burden test) for predicted loss-of-function variants in *PACC1*. We then navigated to the “infectious and parasitic diseases” category by first selecting “health-related outcomes” and then selecting “certain infectious and parasitic diseases”. Among the 54 diseases listed, we shortlisted ICD-10 codes related to any predominantly bacterial infections that retained 18 traits. To account for multiple tests, we applied the Bonferroni correction method. The odds ratio and 95% confidence intervals were estimated from the burden test P-values and the beta statistic values, and the forest plot was generated using GraphPad Prism v10.

*RNA expression data analyses from the Human Protein Atlas (HPA), Cancer Cell Line Encyclopedia (CCLE), and a Genentech study*

The HPA repository was queried to retrieve *PACC1* expression from single-cell RNA sequencing (scRNA-seq) data across different human tissues and bulk RNA sequencing data from the different cell lines that were sequenced by the HPA team (https://www.proteinatlas.org/ENSG00000065600-PACC1) (62). The HPA scRNA-seq data was based on 31 different datasets from published studies, which were processed by the HPA team using a custom pipeline, and expression profiles were generated by normalizing read counts to normalized transcripts per million (nTPM) (63). *PACC1* expression data was available for 69 cell lines, of which expression from 33 cell lines included integrated data from the HPA and CCLE (https://www.proteinatlas.org/humanproteome/cell+line/method). We first focused on these data from the HPA for a more granular view of *PACC1* expression among cell lines that are commonly used in experimental studies. We also retrieved the data from all 1206 cell lines that were available in the HPA, which included the 69 cell lines mentioned previously, 1055 cell lines specific to CCLE, and an additional 151 cell lines incorporated into HPA from a Genentech published study (64) and in-house HPA sequencing. The expression data from these 1206 cell lines were ranked using the rank function from the base package in R (v 4.4.0) (65) and plotted for a broader distribution of *PACC1* expression. All graphs were plotted using Prism v10 (GraphPad, San Diego, CA, USA).

*Bulk RNA sequencing data analyses from other published datasets*

Normalized counts from bulk RNA sequencing of unstimulated murine macrophages (BMDMs) were retrieved from our previous study, Roewe *et al.* (6). The counts were averaged across the five replicates and the mean counts were ranked using the rank function from the base package in R (v 4.4.0) (65). The copy number values from the bulk RNA sequencing of different murine tissue macrophage populations were retrieved from supplementary data in the study by Qie *et al.* (66). Human *PACC1* normalized expression counts were also retrieved from the Immunological Genome Project Immune Cell Atlas (ICA) (67), which includes data from bulk RNA sequencing of sorted immunocytes from healthy human blood (http://immunecellatlas.net). All graphs were plotted using GraphPad Prism v10.

*Proteomic data analyses from a published dataset*

The normalized mass spectrometry intensities of the phagosomal proteins that were labelled by soybean ascorbate peroxidase (APEX)-expressing *Staphylococcus aureus* and *E. coli* were retrieved from the supplementary data of the study by Li *et al.* (68). Data from the three replicates for each bacterial species was averaged, then ranked using the rank function from the base package in R (v 4.4.0) (65). The rank-protein abundance graph was plotted using GraphPad Prism v10.

*Statistical analysis*

Statistical analyses were performed with GraphPad Prism v10 software. Data from *in vitro* and *in vivo* experiments are representative of at least two independent experiments. Exact sample sizes and number of technical and biological replicates are included in the figure images (e.g., each biological replicate or mouse per group is indicated by a circle) and/or captions. Data in bar graphs represent mean ± standard error of the mean (SEM). Data in box pots represent median and interquartile range with whiskers extended to the minimum and maximum values. Tests for normality (D'Agostino & Pearson test, Anderson-Darling test, Shapiro-Wilk test, Kolmogrov-Smirnov test) were conducted prior to further analysis. Comparisons of two groups were performed based on the two-sided Student's t test while multiple group testing was done through one-way analysis of variance (ANOVA) with Sidak, Tukey, or Dunnet correction. Survival data was analyzed by Mantel-Cox log-rank test. Values of $P<0.05$ were considered significant. *P<0.05, **P<0.01, ***P<0.001 and ****P<0.0001. Statistical analyses for sequencing data were performed as described in the above sections.

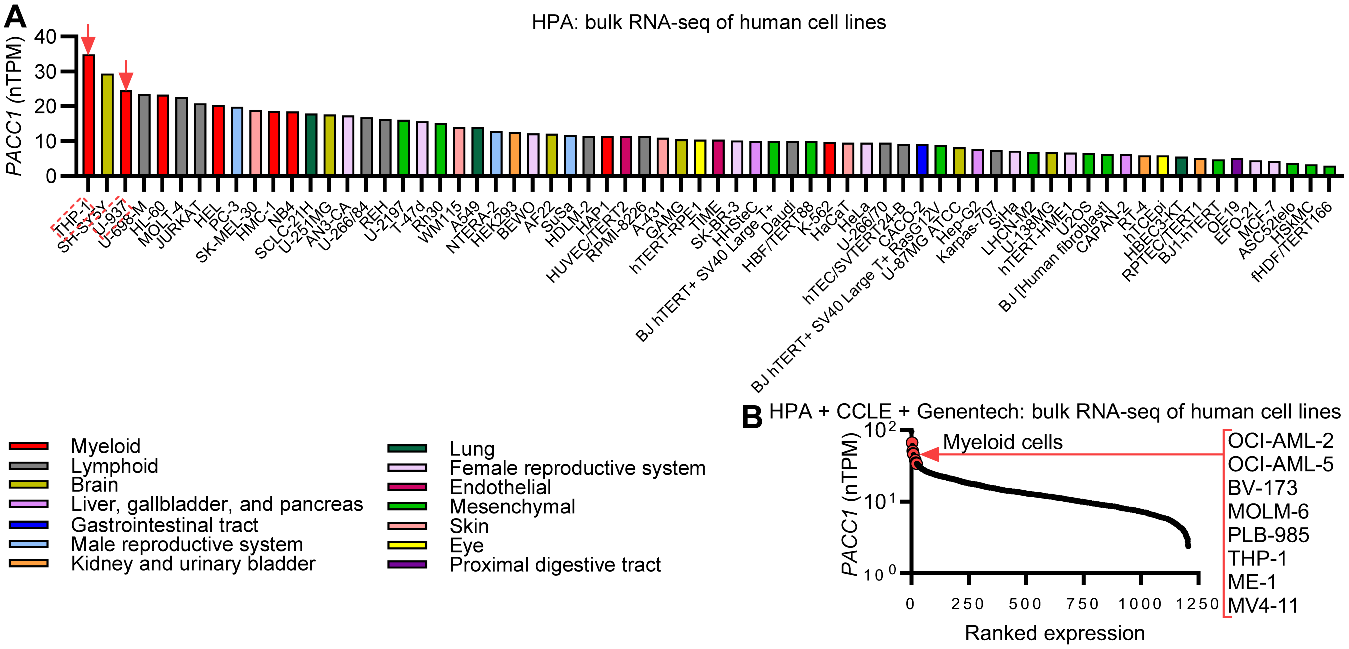


**Fig. S1. *PACC1* is enriched in human myeloid cell-derived cell lines, especially THP-1 cells, a monocyte/macrophage cell line.**

(**A**) Ranked expression values from bulk RNA sequencing data in the Human Protein Atlas (HPA) of 69 human cell lines. Data as normalized transcripts per million, corresponding to mean values of the different individual samples from each cell type. Expression from 33 overlapping cell lines includes integrated data from the HPA and Cancer Cell Line Encyclopedia (CCLE). Red arrows/boxes indicate cells of interest (i.e., monocyte/macrophage cell lines). (**B**) Ranked expression values from bulk RNA sequencing in the HPA in combination with an expanded dataset of over 1000 human cell lines from the CCLE and a Genentech study. Data show normalized transcripts per million, corresponding to mean values of the different individual samples from each cell type. Red circles and arrows indicate cell lines of myeloid lineage in the top 2% of the rankings.

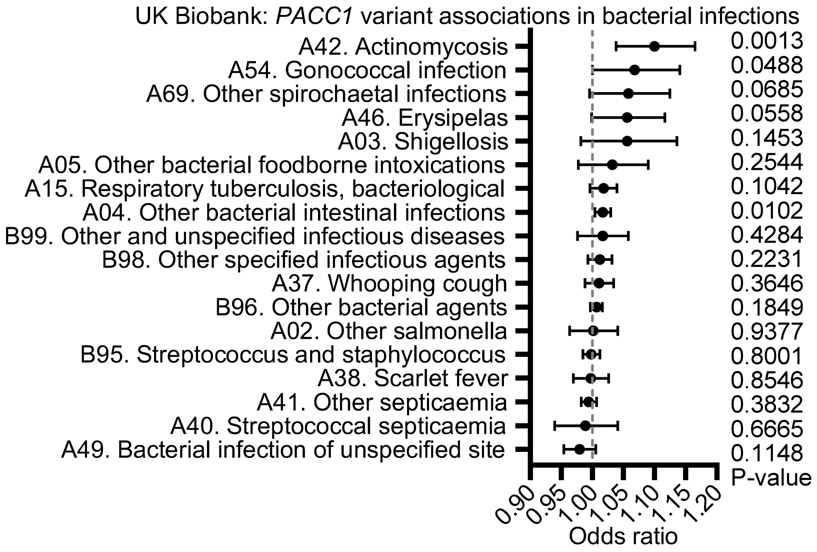


**Fig. S2. Analysis of associations between human *PACC1* variants and the risk of bacterial infections.**

Forest plot showing associations between gene-level predicted loss-of-function variant burden in *PACC1* with a curated list of any clinical codes characterizing predominantly bacterial infections, from the UK Biobank accessed through the Genebass platform. Infection associations with burden test p-values, odds ratios, and 95% confidence interval from this large patient cohort (n=~400,000 controls) are shown. Infections include 10th revision of the International Classification of Diseases (ICD-10) A04, “Other bacterial intestinal infections,” (n=5769 cases) which consists predominantly of *E. coli* infections.

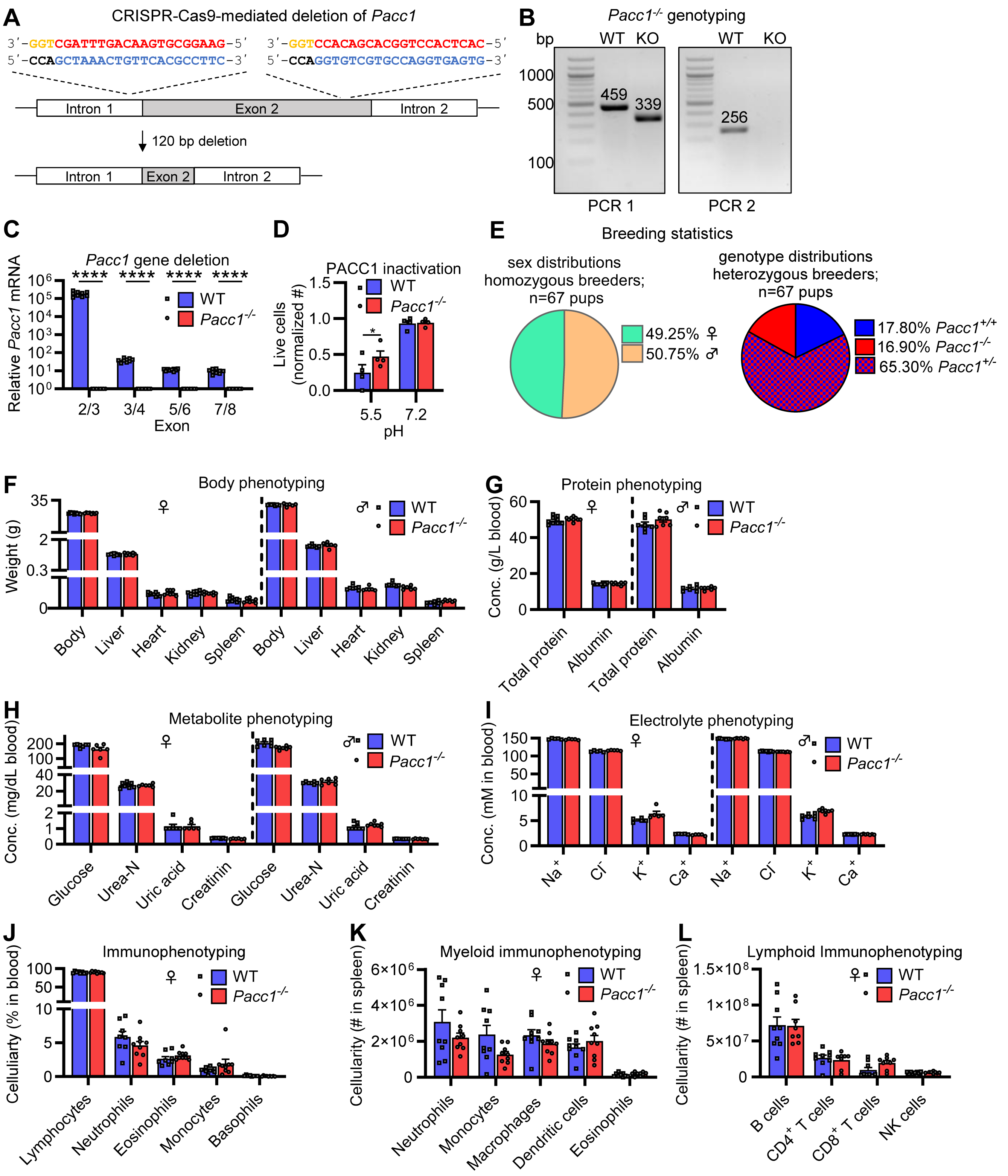

A
CRISPR-Cas9-mediated deletion of Pacc1
3′-GGTCGATTTGACAAGTGCGGAAG-5′
5′-CCAGCTAAACTGTTCACGCCTTC-3′
3′-GGTCCACAGCACGGTCCACTCAC-5′
5′-CCAGGTGTCGTGCCAGGTGAGTG-3′
Intron 1
Exon 2
Intron 2
120 bp deletion
B
Pacc1-/- genotyping
bp
WT
KO
1000
500
100
459
339
256
PCR 1
PCR 2
C
Pacc1 gene deletion
Relative Pacc1 mRNA
WT
Pacc1-/-
Exon
D
PACC1 inactivation
Live cells (normalized #)
pH
E
Breeding statistics
sex distributions homozygous breeders; n=67 pups
49.25% ♀
50.75% ♂
genotype distributions heterozygous breeders; n=67 pups
17.80% Pacc1+/+
16.90% Pacc1-/-
65.30% Pacc1+/-
F
Body phenotyping
Weight (g)
Body
Liver
Heart
Kidney
Spleen
G
Protein phenotyping
Conc. (g/L blood)
Total protein
Albumin
H
Metabolite phenotyping
Conc. (mg/dL blood)
Glucose
Urea-N
Uric acid
Creatinin
I
Electrolyte phenotyping
Conc. (mM in blood)
Na+
Cl-
K+
Ca+
J
Immunophenotyping
Celluliarty (% in blood)
Lymphocytes
Neutrophils
Eosinophils
Monocytes
Basophils
K
Myeloid immunophenotyping
Cellularity (# in spleen)
Neutrophils
Monocytes
Macrophages
Dendritic cells
Eosinophils
L
Lymphoid Immunophenotyping
Cellularity (# in spleen)
B cells
CD4+ T cells
CD8+ T cells
NK cells

**Fig S3. *De novo* generation of *Pacc1* knockout mice.**

(**A**) Schematic showing generation of *Pacc1* knockout ($^{-/-}$, KO) mice. A CRISPR/Cas9 approach used guide RNA sequences (red) aligned to intron 1 and 2 sequences (blue) flanking exon 2 such that protospacer adjacent motif (PAM) sequence (yellow)-mediated cleavage and recombination resulted in a 120 base pair (bp) deletion, causing a frame shift and premature stop codon. (**B**) Representative gels showing PCR-based genotyping of homozygous wildtype (WT) and *Pacc1*$^{-/-}$ mice. PCR 1 primers aligned to intron 1 and 2, resulting in a truncated KO band at the indicated bp length, as compared to the WT band. PCR 2 primers aligned to exon 2, resulting in no amplified KO band and a WT band at the indicated base pair (bp) length. (**C**) *Pacc1* expression by qRT-PCR using primers aligned to the indicated exons in bone-marrow derived macrophages (BMDMs) from WT and *Pacc1*$^{-/-}$ mice. Expression normalized to Pacc1$^{-/-}$ for each exon. (**D**) Differences in cell survival between WT and *Pacc1*$^{-/-}$ BMDMs were determined after 24 h of incubation at pH 5.5 and pH 7.2 using flow cytometry. (**E**) Breeding statistics of heterozygous and homozygous *Pacc1*$^{-/-}$ mice showing normal viable offspring and normal sex ratios following homozygous breeding, and slightly below 25% Mendelian ratios of *Pacc1* deficiency following heterozygous breeding. (**F-L**) 16-week-old male and female WT and *Pacc1*$^{-/-}$ mice were phenotyped as indicated. (**F**) Body and organ weights in WT and *Pacc1*$^{-/-}$ mice. (**G**) Blood total proteins in WT and *Pacc1*$^{-/-}$ mice by Alinity Analyzer. (**H**) Blood metabolites in WT and *Pacc1*$^{-/-}$ mice by Alinity Analyzer. (**I**) Blood electrolytes in WT and *Pacc1*$^{-/-}$ mice by Alinity Analyzer. (**J**) Blood immune cellularity (frequencies) in WT and *Pacc1*$^{-/-}$ mice in these mice by flow cytometry. (**K**) Splenic myeloid cellularity (counts) in WT and *Pacc1*$^{-/-}$ mice by flow cytometry. (**L**) Splenic lymphoid cellularity (counts) in WT and *Pacc1*$^{-/-}$ mice by flow cytometry. For **C**, **D**, **F**-**L**, data represent ≥2 independent experiments. Data show one symbol per mouse, mean±SEM, *P<0.5, ****P<0.0001, Student's t-test.

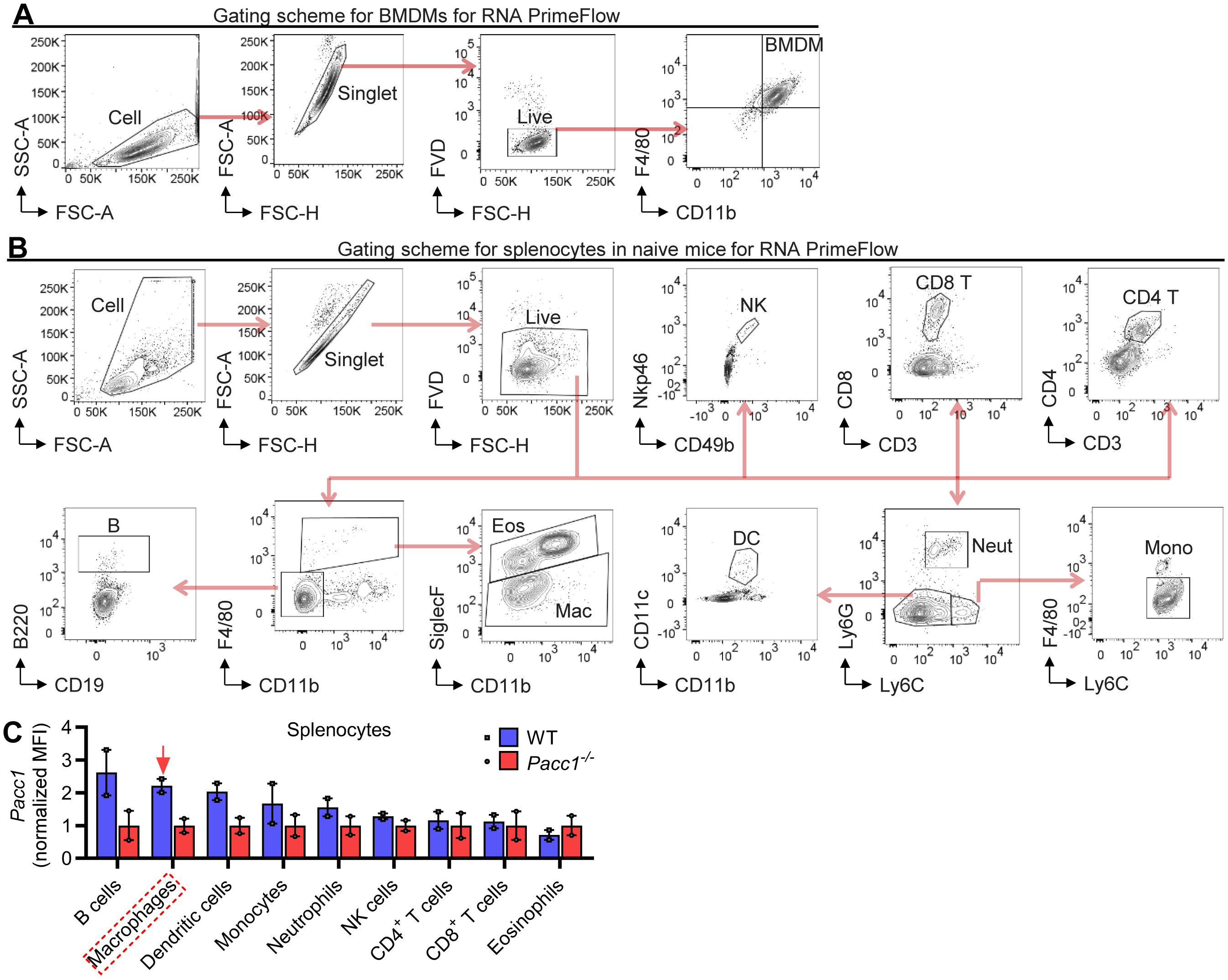


**Fig. S4. RNA PrimeFlow analysis of mouse bone-marrow derived macrophages (BMDMs) and splenocytes.**

(**A**) Gating scheme for flow cytometric analysis of macrophages (BMDMs) as in **Fig. 2B**. Singlet cells were discriminated using forward scatter (FSC) and side scatter (SSC), and live cells were defined as negative for fixable viable dye eFluro780 (FVD). BMDMs were defined as CD11b$^+$F4/80$^+$. (**B**) Gating scheme for flow cytometric analysis of splenocytes in naïve adult mice in **Fig. 2C** and for FACSorting of splenocyte populations in **Fig. 2D**. Singlet cells were discriminated using FSC and SSC, and live cells were defined as negative for FVD. NK cells (NK) were defined as CD49b$^+$NKp46$^+$, CD8 T cells (CD8 T) as CD3$^+$CD8$^+$, CD4 T cells (CD4 T) as CD3$^+$CD4$^+$, B cells (B) as F4/80$^-$CD11b$^{lo/-}$CD19$^+$B220$^+$, eosinophils (Eos) as CD11b$^+$SiglecF$^+$ confirmed via back-gating to be SSC-A$^{hi}$, macrophages (Mac) as CD11b$^+$F4/80$^+$SiglecF$^-$, dendritic cells (DCs) as Ly6C$^-$Ly6G$^-$CD11c$^+$CD11b$^+$, neutrophils (Neut) as Ly6C$^+$Ly6G$^+$ confirmed via back gating to be CD11b$^+$, and monocytes (Mono) as Ly6C$^{hi}$Ly6G$^-$F4/80$^-$ confirmed via back gating to be CD11b$^+$. (**C**) Geometric mean fluorescence intensities (MFIs) of *Pacc1* expression in splenocytes from two independent experiments represented in **Fig. 2C**, normalized to *Pacc1*$^{-/-}$ values for each cell type and ranked by wildtype (WT) values. Data show one symbol per mouse, mean±SEM.

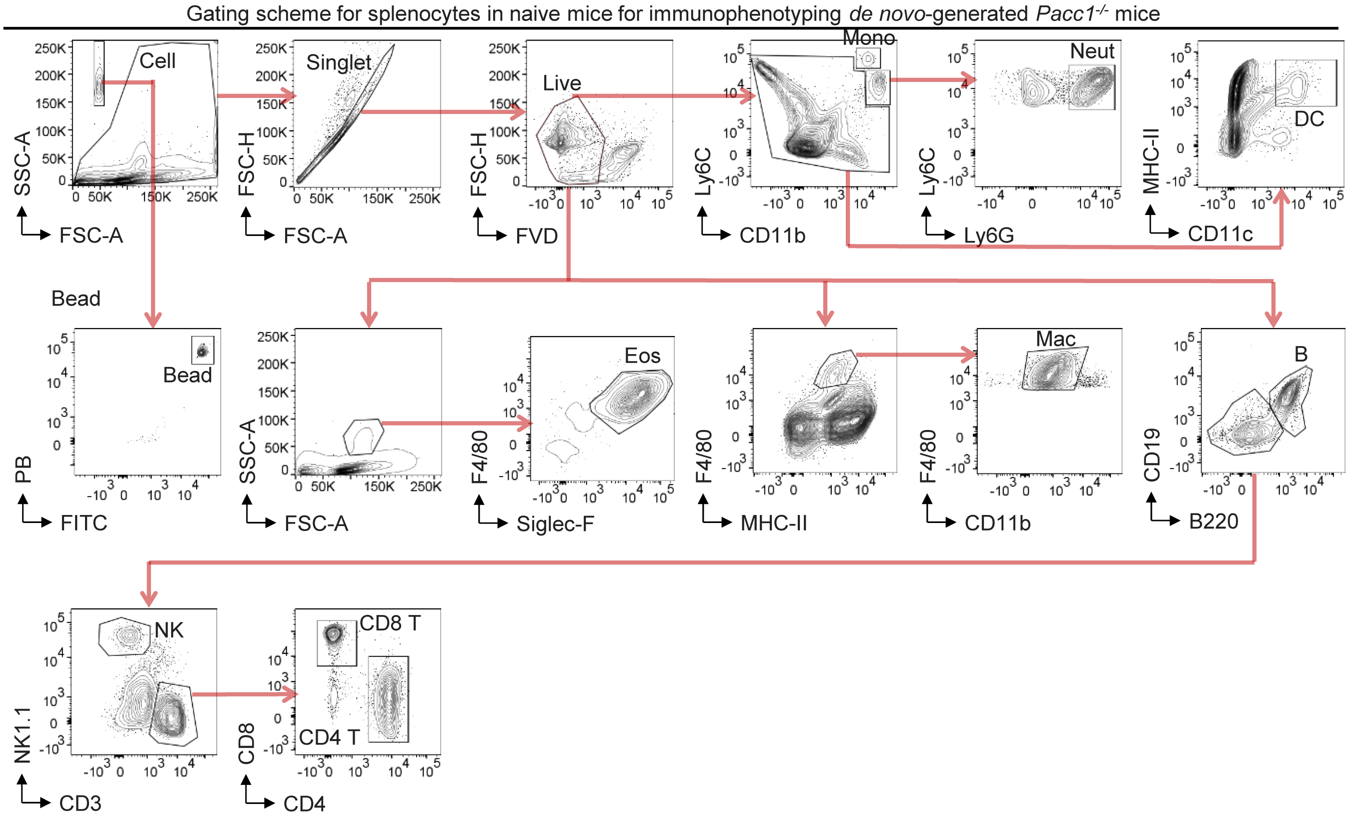


**Fig. S5. Gating scheme for splenocytes in naïve mice for immunophenotyping of *de novo*-generated *Pacc1*$^{-/-}$ mice.**

Gating scheme for flow cytometric analysis of splenocytes in naïve adult mice in **Fig. S3K-L**. Singlet cells were discriminated using forward scatter (FSC) and side scatter (SSC), and live cells were defined as negative for fixable viable dye eFluro780 (FVD). Monocytes (Mono) were defined as Ly6C$^{hi}$CD11b$^{hi}$, neutrophils (Neut) as Ly6C$^{+}$CD11b$^{hi}$Ly6G$^{+}$, Dendritic cells (DC) as CD11c$^{+}$MHC-II$^{+}$, eosinophils (Eos) as SSC-A$^{hi}$Siglec-F$^{+}$, macrophages (Mac) as F4/80$^{+}$MHC-II$^{+}$CD11b$^{+}$, B cells (B) as CD19$^{+}$B220$^{+}$, Natural Killer cells (NK) as CD19$^{-}$B220$^{-}$CD3$^{-}$NK1.1$^{+}$, CD8$^{+}$ T cells (CD8 T) as CD19$^{-}$B220$^{-}$NK1.1$^{-}$CD3$^{+}$CD8$^{+}$, and CD4$^{+}$ T cells (CD4 T) as CD19$^{-}$B220$^{-}$NK1.1$^{-}$CD3$^{+}$CD8$^{+}$. Counting beads were discriminated based on low FSC-A and high autofluorescence in the indicated channels.

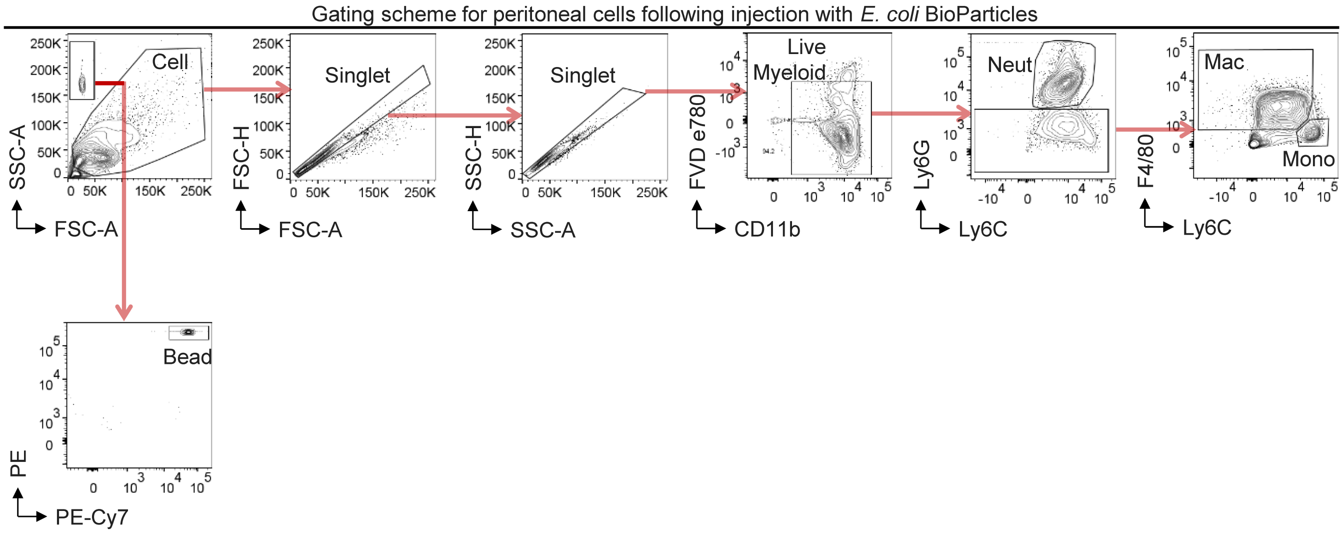


**Fig. S6. Gating scheme for peritoneal cells following injection with *E. coli* BioParticles.**
Gating scheme for flow cytometric analysis of peritoneal cells following injection with inactivated *E. coli* BioParticles as in **Fig. 3F-I**. Singlet cells were discriminated using forward scatter (FSC) and side scatter (SSC), and live cells were defined as negative for fixable viability dye eFluro780 (FVD). Myeloid cells (Live myeloid) were defined as CD11b$^{+}$, neutrophils (Neut) as CD11b$^{+}$Ly6C$^{+}$Ly6G$^{+}$, macrophages (Mac) as CD11b$^{+}$Ly6G$^{-}$Ly6C$^{-/lo}$F4/80$^{+}$, and monocytes (Mono) as CD11b$^{+}$Ly6G$^{-}$Ly6C$^{hi}$. Counting beads were discriminated based on low FSC-A and high autofluorescence in the indicated channels.

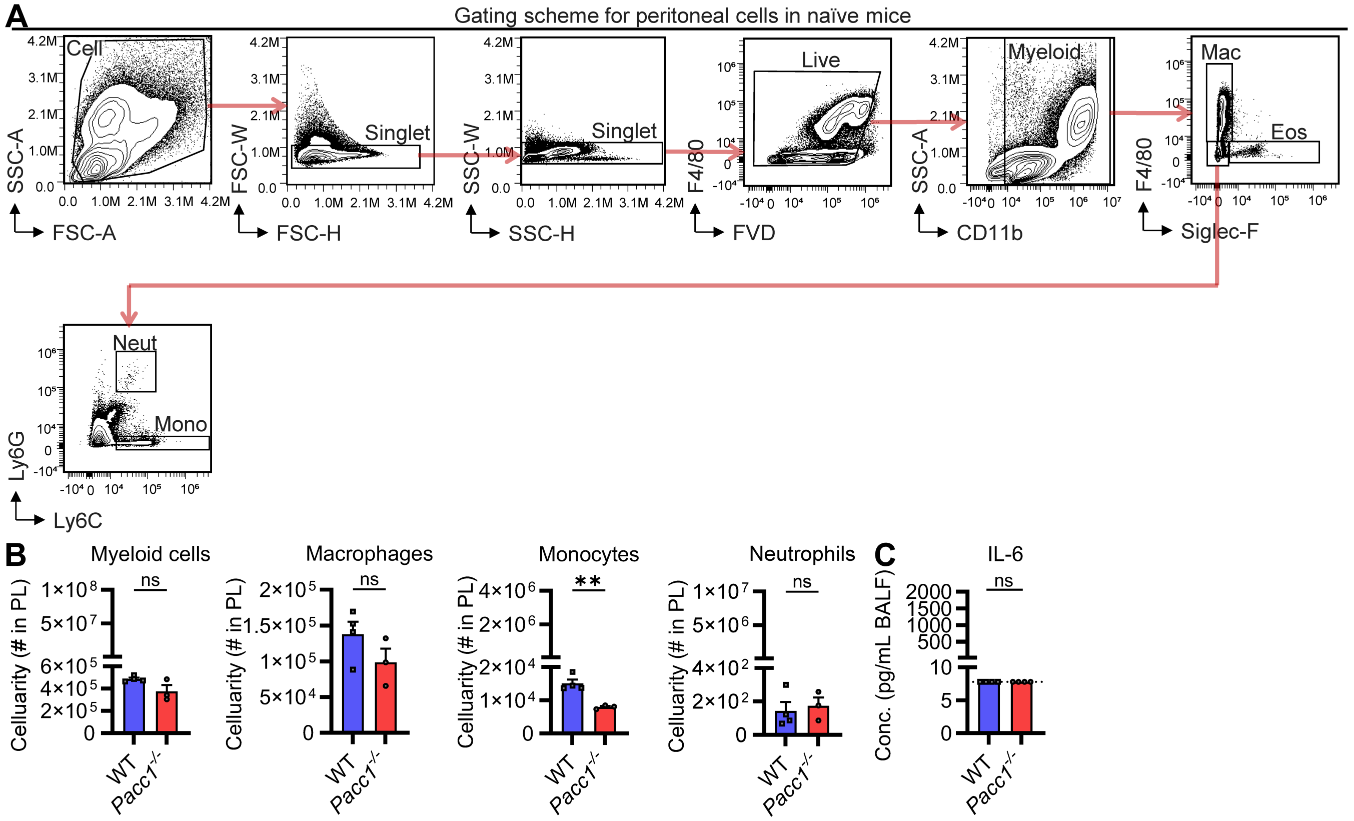


**Fig. S7. *Pacc1*$^{-/-}$ mice do not show aberrant inflammation in the peritoneum at baseline.**
(**A**) Gating scheme for flow cytometric analysis of cells from peritoneal lavages in naïve mice. Singlet cells were discriminated using forward scatter (FSC) and side scatter (SSC), and live cells were defined as negative for fixable viability dye eFluro780 (FVD). Myeloid cells (Live myeloid) were defined as CD11b$^{+}$, macrophages (Mac) as CD11b$^{+}$F4/80$^{+}$Siglec-F$^{-}$, eosinophils (Eos) as CD11b$^{+}$SiglecF$^{+}$ confirmed via back-gating to be SSC-A$^{hi}$, neutrophils (Neut) as CD11b$^{+}$F4/80$^{-}$SiglecF$^{-}$Ly6C$^{+}$Ly6G$^{+}$, and monocytes (Mono) as CD11b$^{+}$F4/80$^{-}$Siglec-F$^{-}$Ly6G$^{-}$Ly6C$^{+}$. (**B**) Cell numbers from the peritoneal lavage (PL) of naïve mice by flow cytometry. (**C**) IL-6 concentration in peritoneal lavage fluid (PLF) of naïve mice by ELISA. The dotted line indicates the limit of detection. Data represent ≥2 independent experiments. Data show one symbol per mouse, mean±SEM, ns=not significant, Student's t-test.

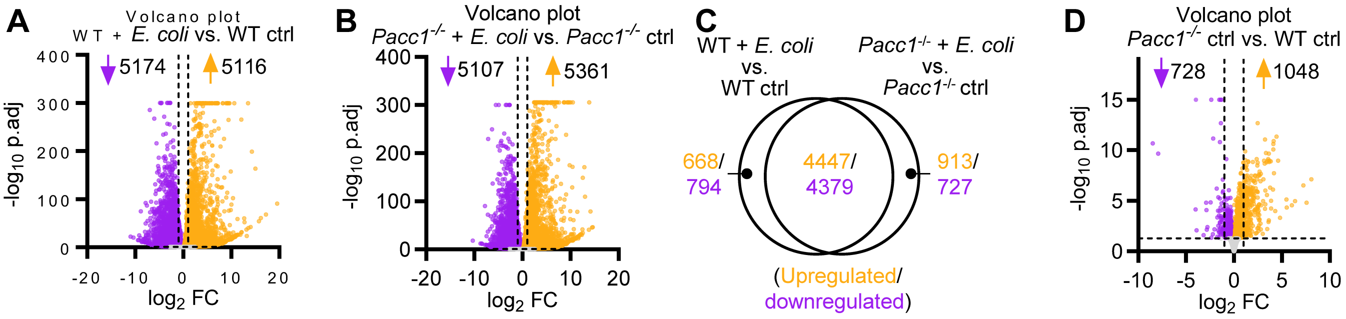


**Fig. S8. Global transcriptomic changes in bone marrow-derived macrophages (BMDMs) are primarily driven by *E. coli* BioParticle stimulus, followed by genotype (*Pacc1*$^{-/-}$).**
Wildtype (WT) and *Pacc1*$^{-/-}$ BMDMs were stimulated with *E. coli* BioParticles at 100 µg/mL or left as unstimulated controls (Ctrl) for 12 h and subjected to bulk RNA sequencing. (**A**) Volcano plot of differentially expressed genes (DEGs) in WT BMDMs with *E. coli* vs. unstimulated controls. (**B**) Volcano plot of DEGs in *Pacc1*$^{-/-}$ BMDMs with *E. coli* vs. unstimulated controls. (**C**) Venn diagram of significant DEGs with adj. P<0.05 in WT and *Pacc1*$^{-/-}$ BMDMs with *E. coli* vs. unstimulated controls. (**D**) Volcano plot of DEGs in *Pacc1*$^{-/-}$ vs. WT BMDM unstimulated controls. For volcano plots, dotted lines indicate adj. P=0.05 and absolute $\log_2$ fold change (FC)=1; and values for $-\log_{10}$ adj. P>300 were set to 300 (**A**, **B**), or values >15 were set to 15 (**D**), for visualization. See also ***SI Appendix*, Dataset S1**.

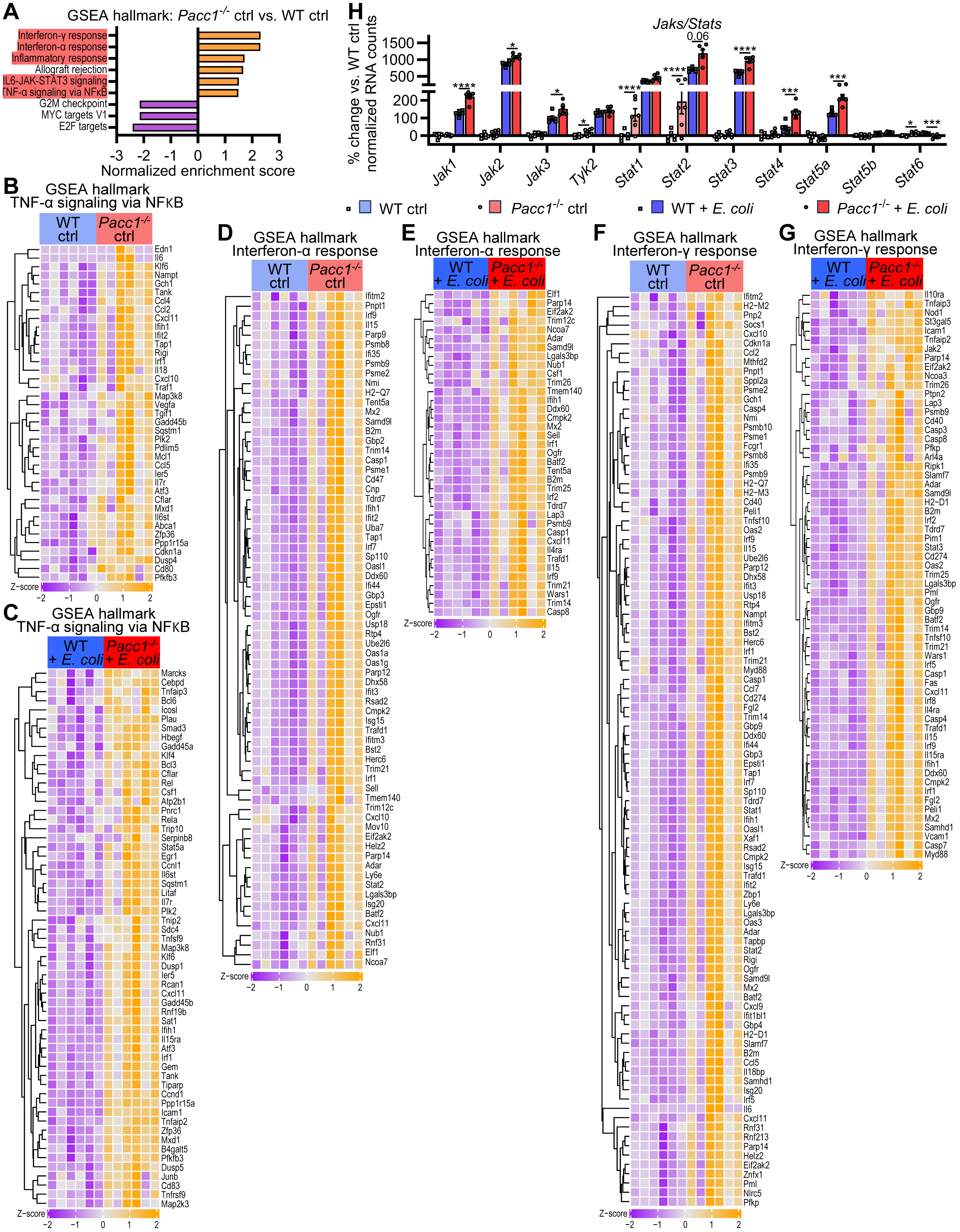
A
GSEA hallmark: Pacc1-/- ctrl vs. WT ctrl
Interferon-γ response
Interferon-α response
Inflammatory response
Allograft rejection
IL6-JAK-STAT3 signaling
TNF-α signaling via NFκB
G2M checkpoint
MYC targets V1
E2F targets
-3 -2 -1 0 1 2 3
Normalized enrichment score
H
Jaks/Stats
% change vs. WT ctrl normalized RNA counts
1500 1000 500 200 100 0
0.06
Jak1 Jak2 Jak3 Tyk2 Stat1 Stat2 Stat3 Stat4 Stat5a Stat5b Stat6
WT ctrl
Pacc1-/- ctrl
WT + E. coli
Pacc1-/- + E. coli
B
GSEA hallmark
TNF-α signaling via NFκB
WT ctrl
Pacc1-/- ctrl
Edn1 Il6 Klf6 Nampt Gch1 Tank Ccl4 Ccl2 Cxcl11 Ifih1 Ifit2 Tap1 Rigi Irf1 Il18 Cxcl10 Traf1 Map3k8 Vegfa Tgif1 Gadd45b Sqstm1 Plk2 Pdlim5 Mcl1 Ccl5 Ier5 Il7r Atf3 Cflar Mxd1 Il6st Abca1 Zfp36 Ppp1r15a Cdkn1a Dusp4 Cd80 Pfkfb3
Z-score
−2 −1 0 1 2
C
GSEA hallmark
TNF-α signaling via NFκB
WT + E. coli
Pacc1-/- + E. coli
Marcks Cebpd Tnfaip3 Bcl6 Icosl Plau Smad3 Hbegf Gadd45a Klf4 Bcl3 Cflar Rel Csf1 Atp2b1 Pnrc1 Rela Trip10 Serpinb8 Stat5a Egr1 Ccnl1 Il6st Sqstm1 Litaf Il7r Plk2 Tnip2 Sdc4 Tnfsf9 Map3k8 Klf6 Dusp1 Ier5 Rcan1 Cxcl11 Gadd45b Rnf19b Sat1 Ifih1 Il15ra Atf3 Irf1 Gem Tank Tiparp Ccnd1 Ppp1r15a Icam1 Tnfaip2 Zfp36 Mxd1 B4galt5 Pfkfb3 Dusp5 Junb Cd83 Tnfrsf9 Map2k3
Z-score
−2 −1 0 1 2
D
GSEA hallmark
Interferon-α response
WT ctrl
Pacc1-/- ctrl
Ifitm2 Pnpt1 Irf9 Il15 Parp9 Psmb8 Ifi35 Psmb9 Psme2 Nmi H2-Q7 Tent5a Mx2 Samd9l B2m Gbp2 Trim14 Casp1 Psme1 Cd47 Cnp Tdrd7 Ifih1 Ifit2 Uba7 Tap1 Irf7 Sp110 Oasl1 Ddx60 Ifi44 Gbp3 Epsti1 Ogfr Usp18 Rtp4 Ube2l6 Oas1a Oas1g Parp12 Dhx58 Ifit3 Rsad2 Cmpk2 Isg15 Trafd1 Ifitm3 Bst2 Herc6 Trim21 Irf1 Sell Tmem140 Trim12c Cxcl10 Mov10 Eif2ak2 Helz2 Parp14 Adar Ly6e Stat2 Lgals3bp Isg20 Batf2 Cxcl11 Nub1 Rnf31 Elf1 Ncoa7
Z-score
−2 −1 0 1 2
E
GSEA hallmark
Interferon-α response
WT + E. coli
Pacc1-/- + E. coli
Elf1 Parp14 Eif2ak2 Trim12c Ncoa7 Adar Samd9l Lgals3bp Nub1 Csf1 Trim26 Tmem140 Ifih1 Ddx60 Cmpk2 Mx2 Sell Irf1 Ogfr Batf2 Tent5a B2m Trim25 Irf2 Tdrd7 Lap3 Psmb9 Casp1 Cxcl11 Il4ra Trafd1 Il15 Irf9 Trim21 Wars1 Trim14 Casp8
Z-score
−2 −1 0 1 2
F
GSEA hallmark
Interferon-γ response
WT ctrl
Pacc1-/- ctrl
Ifitm2 H2-M2 Pnp2 Socs1 Cxcl10 Cdkn1a Ccl2 Mthfd2 Pnpt1 Sppl2a Psme2 Gch1 Casp4 Nmi Psmb10 Psme1 Fcgr1 Psmb8 Ifi35 Psmb9 H2-Q7 H2-M3 Cd40 Peli1 Tnfsf10 Oas2 Irf9 Il15 Ube2l6 Parp12 Dhx58 Ifit3 Usp18 Rtp4 Nampt Ifitm3 Bst2 Herc6 Irf1 Trim21 Myd88 Casp1 Ccl7 Cd274 Fgl2 Trim14 Gbp9 Ddx60 Ifi44 Gbp3 Epsti1 Tap1 Irf7 Sp110 Tdrd7 Stat1 Ifih1 Oasl1 Xaf1 Rsad2 Cmpk2 Isg15 Trafd1 Ifit2 Zbp1 Ly6e Lgals3bp Oas3 Adar Tapbp Stat2 Rigi Ogfr Samd9l Mx2 Batf2 Cxcl9 Ifit1bl1 Gbp4 H2-D1 Slamf7 B2m Ccl5 Il18bp Samhd1 Isg20 Irf5 Il6 Cxcl11 Rnf31 Rnf213 Parp14 Helz2 Eif2ak2 Znfx1 Pml Nlrc5 Pfkp
Z-score
−2 −1 0 1 2
G
GSEA hallmark
Interferon-γ response
WT + E. coli
Pacc1-/- + E. coli
Il10ra Tnfaip3 Nod1 St3gal5 Icam1 Tnfaip2 Jak2 Parp14 Eif2ak2 Ncoa3 Trim26 Ptpn2 Lap3 Psmb9 Cd40 Casp3 Casp8 Pfkp Arl4a Ripk1 Slamf7 Adar Samd9l H2-D1 B2m Irf2 Tdrd7 Pim1 Stat3 Cd274 Oas2 Trim25 Lgals3bp Pml Ogfr Gbp9 Batf2 Trim14 Tnfsf10 Trim21 Wars1 Irf5 Casp1 Fas Cxcl11 Irf8 Il4ra Casp4 Trafd1 Il15 Irf9 Il15ra Ifih1 Ddx60 Cmpk2 Irf1 Fgl2 Peli1 Mx2 Samhd1 Vcam1 Casp7 Myd88
Z-score
−2 −1 0 1 2

**Fig. S9. Hallmark interferon, TNF-α, and JAK/STAT signaling are dysregulated in *Pacc1*$^{-/-}$ bone marrow-derived macrophages (BMDMs).**

Wildtype (WT) and *Pacc1*$^{-/-}$ BMDMs were stimulated with *E. coli* BioParticles at 100 µg/mL or left as unstimulated controls (ctrl) for 12 h and subjected to bulk RNA sequencing. (**A**) Bar plot showing hallmark gene set enrichment analysis (GSEA) results in *Pacc1*$^{-/-}$ vs. WT BMDM unstimulated controls. (**B**-**G**) Heatmaps of differentially expressed genes (DEGs) from *Pacc1*$^{-/-}$ vs. WT BMDMs with *E. coli* or as unstimulated controls enriched in hallmark gene sets: (**B-C**) TNF-α signaling via NFkB; (**D**, **E**) interferon-α response; and (**F**, **G**) interferon-γ response. (**H**) Percent change compared to WT ctrl in normalized RNA counts of selected *Jak*/*Stat* genes. Data show one symbol per mouse, mean±SEM, *P<0.5, ****P<0.0001, Wald's test with Benjamini-Hochberg. Heatmaps show normalized expression values with Z-scores from -2/purple to +2/orange and significant differentially expressed leading edge genes driving these processes at adj. P<0.05. See also ***SI Appendix*****, Datasets S1 and S2**.

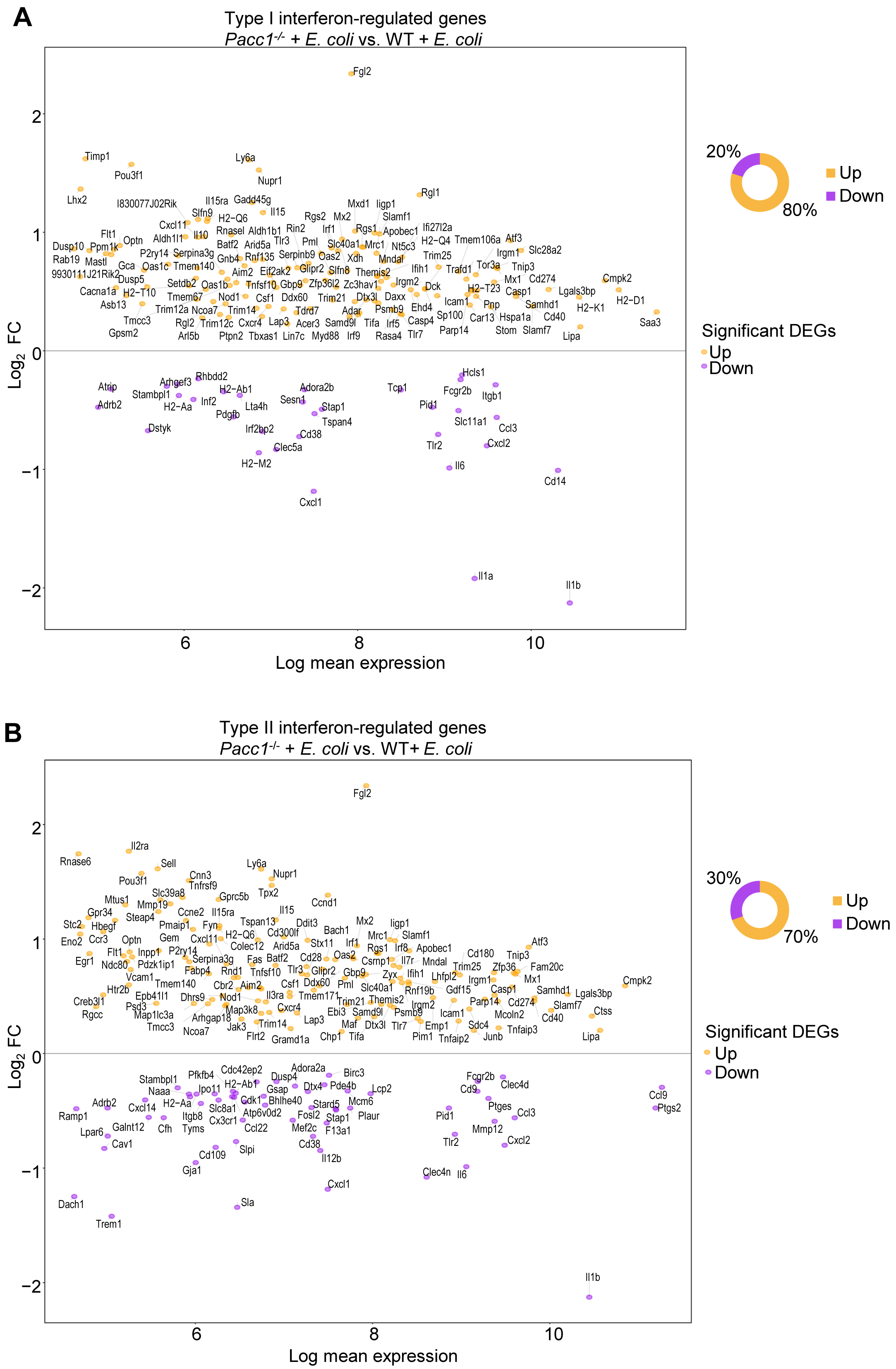
A
Type I interferon-regulated genes
Pacc1-/- + E. coli vs. WT + E. coli
Log2 FC
Log mean expression
2
1
0
−1
−2
6
8
10
Fgl2
Timp1
Pou3f1
Ly6a
Nupr1
Lhx2
I830077J02Rik
Il15ra
Gadd45g
Il15
Rgl1
Mxd1
Iigp1
Slamf1
Cxcl11
Slfn9
H2−Q6
Rgs2
Mx2
Rnasel
Aldh1b1
Rin2
Irf1
Rgs1
Apobec1
Ifi27l2a
Flt1
Aldh1l1
Il10
Tlr3
Pml
Slc40a1
Mrc1
Nt5c3
H2−Q4
Tmem106a
Atf3
Dusp10
Ppm1k
Optn
Batf2
Arid5a
Irgm1
Slc28a2
Rab19
Mastl
P2ry14
Serpina3g
Rnf135
Serpinb9
Oas2
Xdh
Mndal
Trim25
Gca
Oas1c
Tmem140
Gnb4
Tor3a
Tnip3
9930111J21Rik2
Aim2
Eif2ak2
Glipr2
Slfn8
Themis2
Ifih1
Trafd1
Dusp5
Setdb2
Oas1b
Tnfsf10
Gbp9
Zfp36l2
Zc3hav1
Irgm2
Dck
Mx1
Cd274
Cmpk2
Cacna1a
H2−T10
Tmem67
Nod1
Csf1
Ddx60
Trim21
Dtx3l
Daxx
H2−T23
Casp1
Lgals3bp
H2−D1
Asb13
Trim12a
Ncoa7
Trim14
Tdrd7
Adar
Psmb9
Ehd4
Icam1
Pnp
Samhd1
H2−K1
Tmcc3
Rgl2
Trim12c
Cxcr4
Lap3
Acer3
Samd9l
Tifa
Irf5
Casp4
Sp100
Car13
Hspa1a
Cd40
Saa3
Gpsm2
Arl5b
Ptpn2
Tbxas1
Lin7c
Myd88
Irf9
Rasa4
Tlr7
Parp14
Stom
Slamf7
Lipa
Atrip
Arhgef3
Rhbdd2
H2−Ab1
Adora2b
Tcp1
Hcls1
Fcgr2b
Stambpl1
Inf2
Sesn1
Itgb1
Adrb2
H2−Aa
Lta4h
Stap1
Pid1
Pdgfb
Tspan4
Slc11a1
Dstyk
Irf2bp2
Cd38
Ccl3
Clec5a
Tlr2
Cxcl2
H2−M2
Il6
Cd14
Cxcl1
Il1a
Il1b
20%
80%
Up
Down
Significant DEGs
Up
Down
B
Type II interferon-regulated genes
Pacc1-/- + E. coli vs. WT+ E. coli
Log2 FC
Log mean expression
2
1
0
−1
−2
6
8
10
Fgl2
Il2ra
Rnase6
Sell
Ly6a
Pou3f1
Cnn3
Nupr1
Tnfrsf9
Slc39a8
Tpx2
Gprc5b
Mtus1
Mmp19
Ccnd1
Gpr34
Ccne2
Il15ra
Il15
Steap4
Tspan13
Mx2
Stc2
Hbegf
Pmaip1
Fyn
Ddit3
Iigp1
Bach1
Cd300lf
Eno2
Ccr3
Optn
Gem
Cxcl11
H2−Q6
Mrc1
Slamf1
Colec12
Stx11
Irf1
Atf3
Inppl
P2ry14
Arid5a
Rgs1
Irf8
Apobec1
Flt1
Cd180
Tnip3
Egr1
Ndc80
Pdzk1ip1
Serpina3g
Fas
Batf2
Cd28
Oas2
Csmp1
Il7r
Mndal
Trim25
Zfp36
Fam20c
Fabp4
Rnd1
Tnfsf10
Tlr3
Glipr2
Gbp9
Zyx
Ifih1
Lhfpl2
Cmpk2
Vcam1
Tmem140
Cbr2
Aim2
Csf1
Ddx60
Pml
Slc40a1
Rnf19b
Gdf15
Irgm1
Mx1
Htr2b
Epb41l1
Il3ra
Tmem171
Casp1
Samhd1
Lgals3bp
Creb3l1
Dhrs9
Nod1
Trim21
Themis2
Parp14
Cd274
Slamf7
Psd3
Map3k8
Cxcr4
Irgm2
Rgcc
Map1lc3a
Arhgap18
Ebi3
Samd9l
Psmb9
Icam1
Mcoln2
Ctss
Tmcc3
Jak3
Trim14
Lap3
Maf
Dtx3l
Tlr7
Emp1
Sdc4
Tnfaip3
Cd40
Ncoa7
Flrt2
Gramd1a
Chp1
Tifa
Pim1
Tnfaip2
Junb
Lipa
Stambpl1
Pfkfb4
Cdc42ep2
Adora2a
Birc3
Fcgr2b
Clec4d
Naaa
Ipo11
H2−Ab1
Dusp4
Gsap
Dtx4
Pde4b
Lcp2
Cd9
Ccl9
Adrb2
Cxcl14
H2−Aa
Cdk1
Bhlhe40
Stard5
Mcm6
Ptges
Ptgs2
Ramp1
Slc8a1
Itgb8
Atp6v0d2
Fosl2
Stap1
Plaur
Pid1
Ccl3
Galnt12
Cfh
Tyms
Cx3cr1
Ccl22
Mef2c
F13a1
Mmp12
Lpar6
Cav1
Tlr2
Cxcl2
Slpi
Cd38
Cd109
Il12b
Gja1
Clec4n
Il6
Cxcl1
Dach1
Sla
Trem1
Il1b
30%
70%
Up
Down
Significant DEGs
Up
Down

**Fig. S10. Type I and type II interferon responses are altered in *Pacc1*$^{-/-}$ bone marrow-derived macrophages (BMDMs).**

Wildtype (WT) and *Pacc1*$^{-/-}$ BMDMs were stimulated with *E. coli* BioParticles at 100 µg/mL or left as unstimulated controls (ctrl) for 12 h and subjected to bulk RNA sequencing. MA plot of type I and type II interferon-regulated genes among the statistically significant differentially expressed genes (DEGs) in *Pacc1*$^{-/-}$ vs. WT BMDMs stimulated with *E. coli*. Only differentially expressed genes that show an absolute fold change ≥10 in the "Interferome" reference database are shortlisted. Statistical significance for DEGs is adj. P<0.05. See also ***SI Appendix*, Dataset S1**.

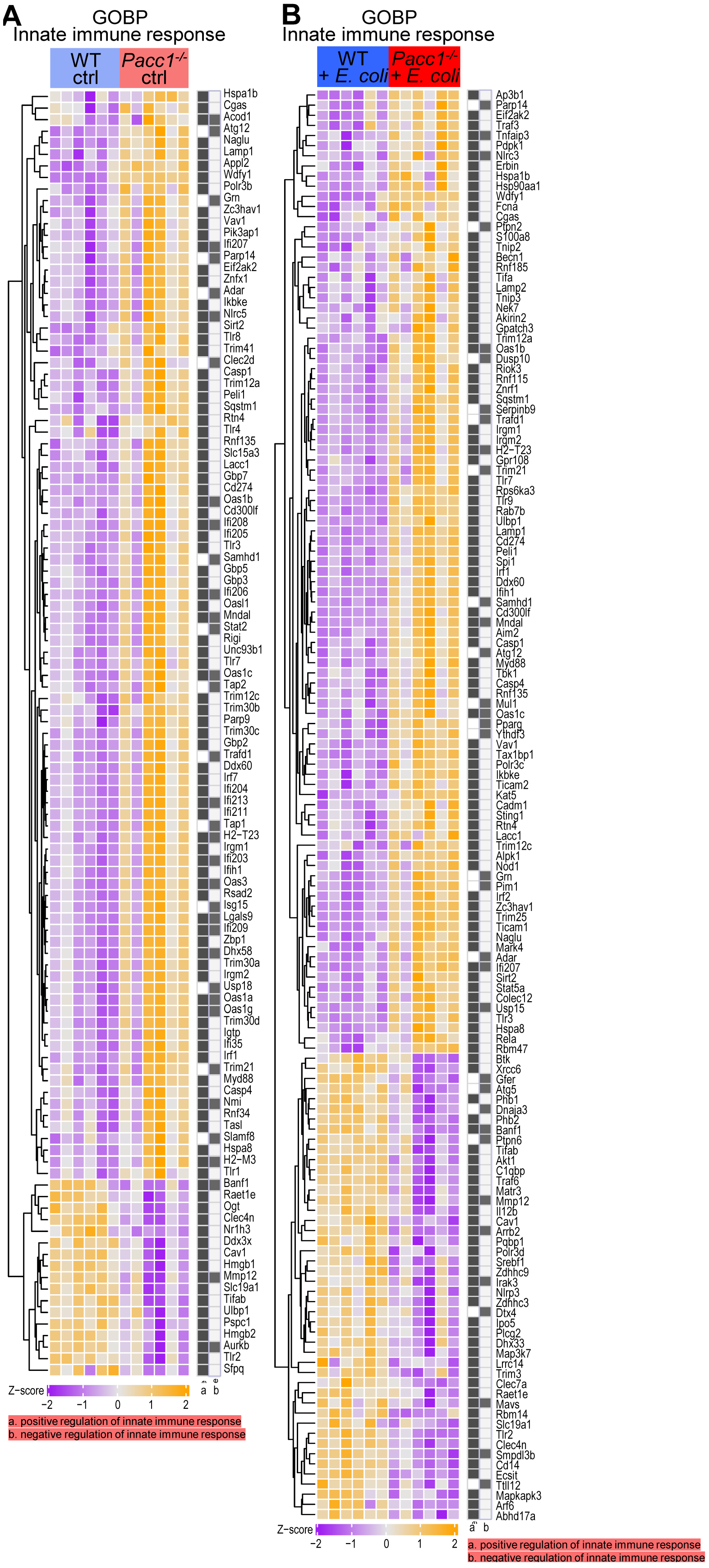
A
GOBP
Innate immune response
WT ctrl
Pacc1-/- ctrl
Hspa1b
Cgas
Acod1
Atg12
Naglu
Lamp1
Appl2
Wdfy1
Polr3b
Grn
Zc3hav1
Vav1
Pik3ap1
Ifi207
Parp14
Eif2ak2
Znfx1
Adar
Ikbke
Nlrc5
Sirt2
Tlr8
Trim41
Clec2d
Casp1
Trim12a
Peli1
Sqstm1
Rtn4
Tlr4
Rnf135
Slc15a3
Lacc1
Gbp7
Cd274
Oas1b
Cd300lf
Ifi208
Ifi205
Tlr3
Samhd1
Gbp5
Gbp3
Ifi206
Oasl1
Mndal
Stat2
Rigi
Unc93b1
Tlr7
Oas1c
Tap2
Trim12c
Trim30b
Parp9
Trim30c
Gbp2
Trafd1
Ddx60
Irf7
Ifi204
Ifi213
Ifi211
Tap1
H2-T23
Irgm1
Ifi203
Ifih1
Oas3
Rsad2
Isg15
Lgals9
Ifi209
Zbp1
Dhx58
Trim30a
Irgm2
Usp18
Oas1a
Oas1g
Trim30d
Igtp
Ifi35
Irf1
Trim21
Myd88
Casp4
Nmi
Rnf34
Tasl
Slamf8
Hspa8
H2-M3
Tlr1
Banf1
Raet1e
Ogt
Clec4n
Nr1h3
Ddx3x
Cav1
Hmgb1
Mmp12
Slc19a1
Tifab
Ulbp1
Pspc1
Hmgb2
Aurkb
Tlr2
Sfpq
a b
Z-score
-2 -1 0 1 2
a. positive regulation of innate immune response
b. negative regulation of innate immune response
B
GOBP
Innate immune response
WT + E. coli
Pacc1-/- + E. coli
Ap3b1
Parp14
Eif2ak2
Traf3
Tnfaip3
Pdpk1
Nlrc3
Erbin
Hspa1b
Hsp90aa1
Wdfy1
Fcna
Cgas
Ptpn2
S100a8
Tnip2
Becn1
Rnf185
Tifa
Lamp2
Tnip3
Nek7
Akirin2
Gpatch3
Trim12a
Oas1b
Dusp10
Riok3
Rnf115
Znrf1
Sqstm1
Serpinb9
Trafd1
Irgm1
Irgm2
H2-T23
Gpr108
Trim21
Tlr7
Rps6ka3
Tlr9
Rab7b
Ulbp1
Lamp1
Cd274
Peli1
Spi1
Irf1
Ddx60
Ifih1
Samhd1
Cd300lf
Mndal
Aim2
Casp1
Atg12
Myd88
Tbk1
Casp4
Rnf135
Mul1
Oas1c
Pparg
Ythdf3
Vav1
Tax1bp1
Polr3c
Ikbke
Ticam2
Kat5
Cadm1
Sting1
Rtn4
Lacc1
Trim12c
Alpk1
Nod1
Grn
Pim1
Irf2
Zc3hav1
Trim25
Ticam1
Naglu
Mark4
Adar
Ifi207
Sirt2
Stat5a
Colec12
Usp15
Tlr3
Hspa8
Rela
Rbm47
Btk
Xrcc6
Gfer
Atg5
Phb1
Dnaja3
Phb2
Banf1
Ptpn6
Tifab
Akt1
C1qbp
Traf6
Matr3
Mmp12
Il12b
Cav1
Arrb2
Pqbp1
Polr3d
Srebf1
Zdhhc9
Irak3
Nlrp3
Zdhhc3
Dtx4
Ipo5
Plcg2
Dhx33
Map3k7
Lrrc14
Trim3
Clec7a
Raet1e
Mavs
Rbm14
Slc19a1
Tlr2
Clec4n
Smpdl3b
Cd14
Ecsit
Tll12
Mapkapk3
Arf6
Abhd17a
a b
Z-score
-2 -1 0 1 2
a. positive regulation of innate immune response
b. negative regulation of innate immune response

**Fig. S11. Innate immune responses are dysregulated in *Pacc1*$^{-/-}$ bone marrow-derived macrophages (BMDMs).**

Wildtype (WT) and *Pacc1*$^{-/-}$ BMDMs were stimulated with *E. coli* BioParticles at 100 µg/mL or left as unstimulated controls (ctrl) for 12 h and subjected to bulk RNA sequencing. (**A-B**) Heatmap of significant differentially expressed genes (DEGs) *Pacc1*$^{-/-}$ vs. WT BMDMs (**A**) left as unstimulated controls and (**B**) with *E. coli* that are enriched in the gene ontology biological processes (GOBPs) for positive (GO:0045089) and negative (GO:0045824) regulation of the innate immune response. Heatmaps show normalized expression values with Z-score from -2/purple to +2/orange and statistical significance for DEGs at adj. $P<0.05$. Genes enriched in each process have been indicated within the shaded squares. See also ***SI Appendix*, Dataset S4**.

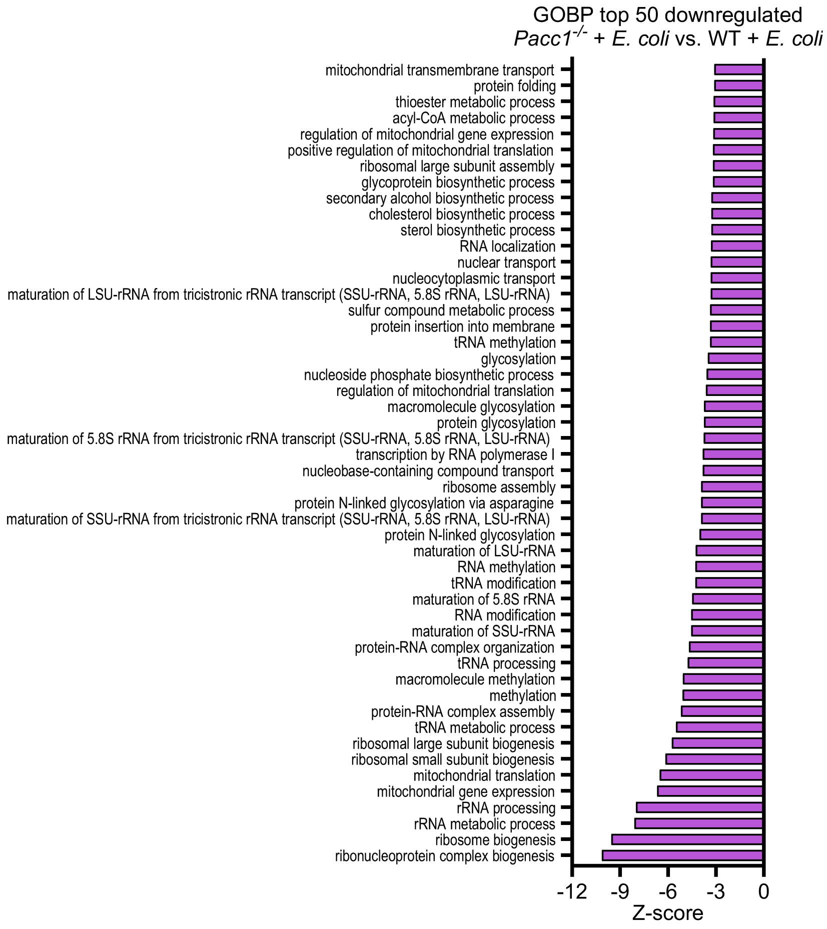


**Fig. S12. RNA processing is dysregulated in *Pacc1$^{-/-}$* bone marrow-derived macrophages (BMDMs).**

Wildtype (WT) and *Pacc1$^{-/-}$* BMDMs were stimulated with *E. coli* BioParticles at 100 µg/mL or left as unstimulated controls (ctrl) for 12 h and subjected to bulk RNA sequencing. Bar plot showing top 50 significant (adj. P<0.1) negatively enriched gene ontology biological processes (GOBPs) in *Pacc1$^{-/-}$* vs. WT BMDMs with *E. coli*. See also ***SI Appendix*, Dataset S4**.

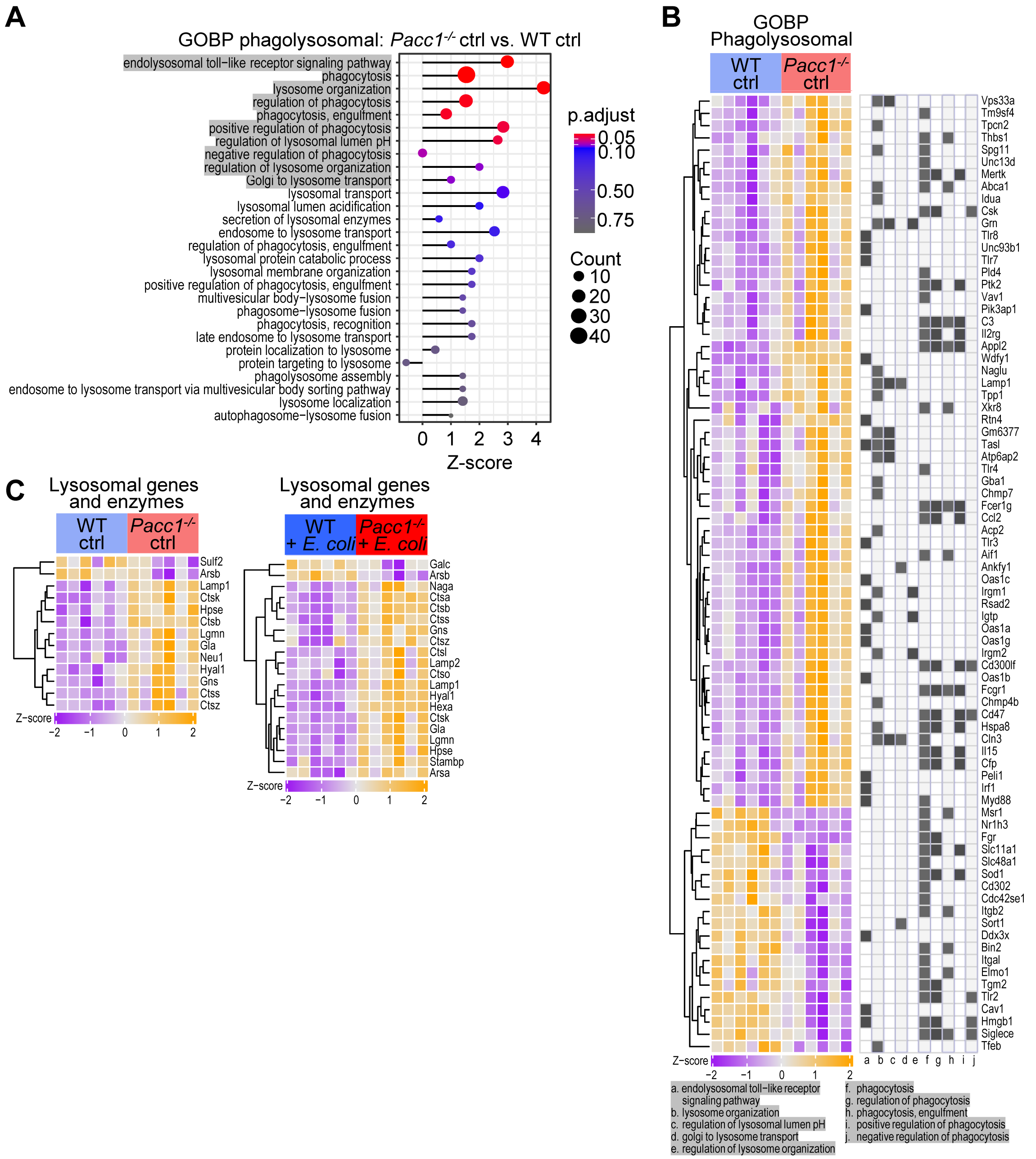

**Fig. S13. Phagolysosomal processes are dysregulated in *Pacc1*$^{-/-}$ bone marrow-derived macrophages (BMDMs).**

Wildtype (WT) and *Pacc1*$^{-/-}$ BMDMs were stimulated with *E. coli* BioParticles at 100 µg/mL or left as unstimulated controls (ctrl) for 12 h and subjected to bulk RNA sequencing. (**A**) Lollipop plot of manually curated phagolysosomal gene ontology biological processes (GOBPs) showing both enriched (adj. P<0.1) and non-enriched in *Pacc1*$^{-/-}$ vs. WT BMDM unstimulated controls. (**B**) Heatmap of significant differentially expressed genes (DEGs) within enriched (adj. P<0.1) phagolysosomal GOBPs from panel **A** in *Pacc1*$^{-/-}$ vs. WT BMDM unstimulated controls. Genes enriched in each GOBP have been indicated within the shaded squares. See also ***SI Appendix*, Dataset S4**. (**C**) Heatmap of manually curated lysosomal genes/enzymes showing significant DEGs in *Pacc1*$^{-/-}$ vs. WT BMDMs left as unstimulated controls and with *E. coli*. Heatmaps show normalized expression values with Z-score from -2/purple to +2/orange and statistical significance for DEGs at adj. P<0.05.

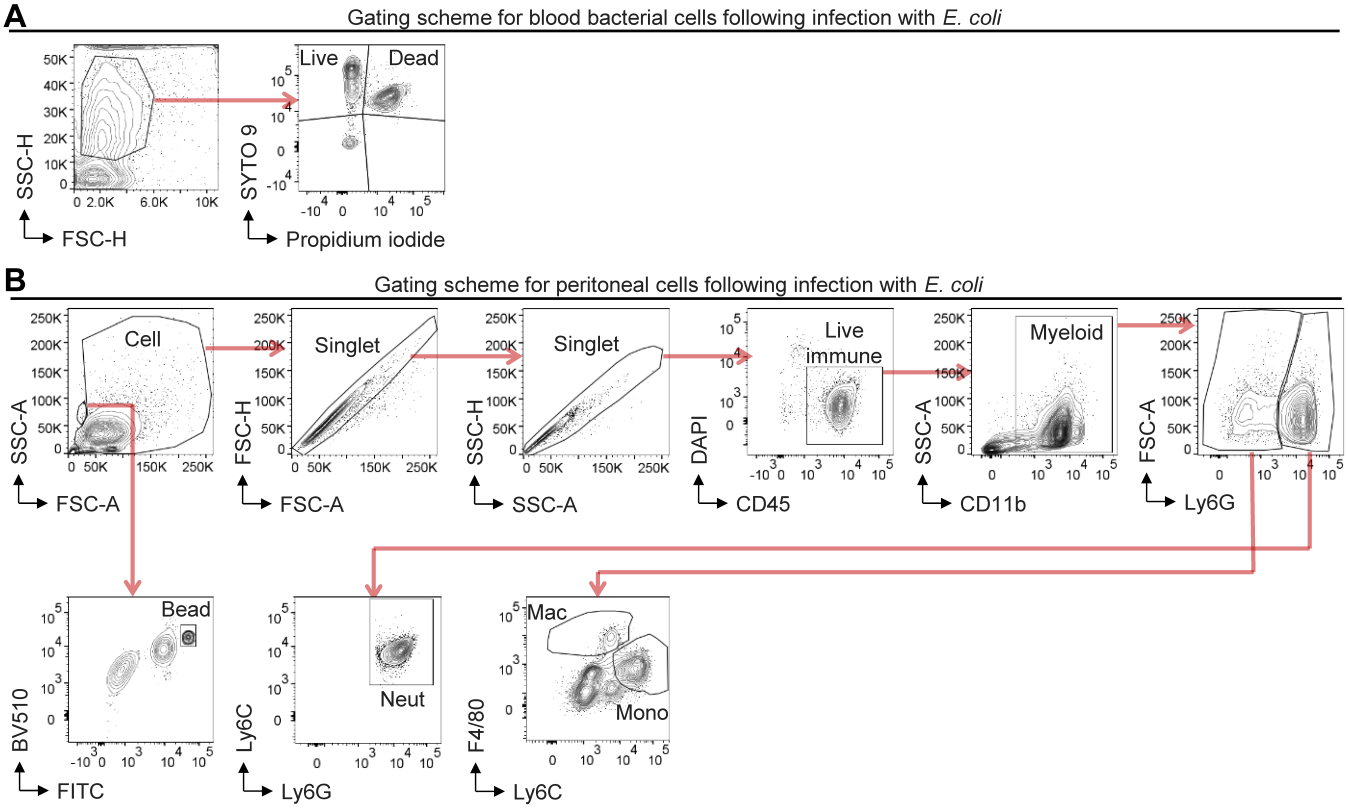


**Fig. S14. Gating schemes for peritoneal cells and bacteria in blood following peritoneal infection.** (**A**) Gating scheme for flow cytometric analysis of bacteria in blood following infection with live *E. coli* as in **Fig. 5D**. Blood was isolated and stained with cell membrane-permeable SYTO 9 nuclear stain to mark all cells, and non-cell membrane-permeable propidium iodide nuclear stain to mark dead cells with disrupted membranes. Bacterial cells were distinguished based on very low forward scatter (FSC). Live *E. coli* were defined as positive for SYTO 9 and negative for propidium iodide. (**B**) Gating scheme for flow cytometric analysis of peritoneal cells following injection with live *E. coli* as in **Fig. 5E**. Singlet cells were discriminated using FSC and side scatter (SSC), and live cells were defined as negative for DAPI. Immune cells (Live immune) were defined as CD45$^{+}$. Myeloid cells (Myeloid) were defined as CD45$^{+}$CD11b$^{+}$, neutrophils as CD45$^{+}$CD11b$^{+}$Ly6C$^{+}$Ly6G$^{+}$, macrophages as CD45$^{+}$CD11b$^{+}$Ly6G$^{-}$Ly6C$^{-/lo}$F4/80$^{+}$, and monocytes as CD45$^{+}$CD11b$^{+}$Ly6G$^{-}$Ly6C$^{hi}$. Counting beads were discriminated based on low FSC-A and high autofluorescence in the indicated channels.

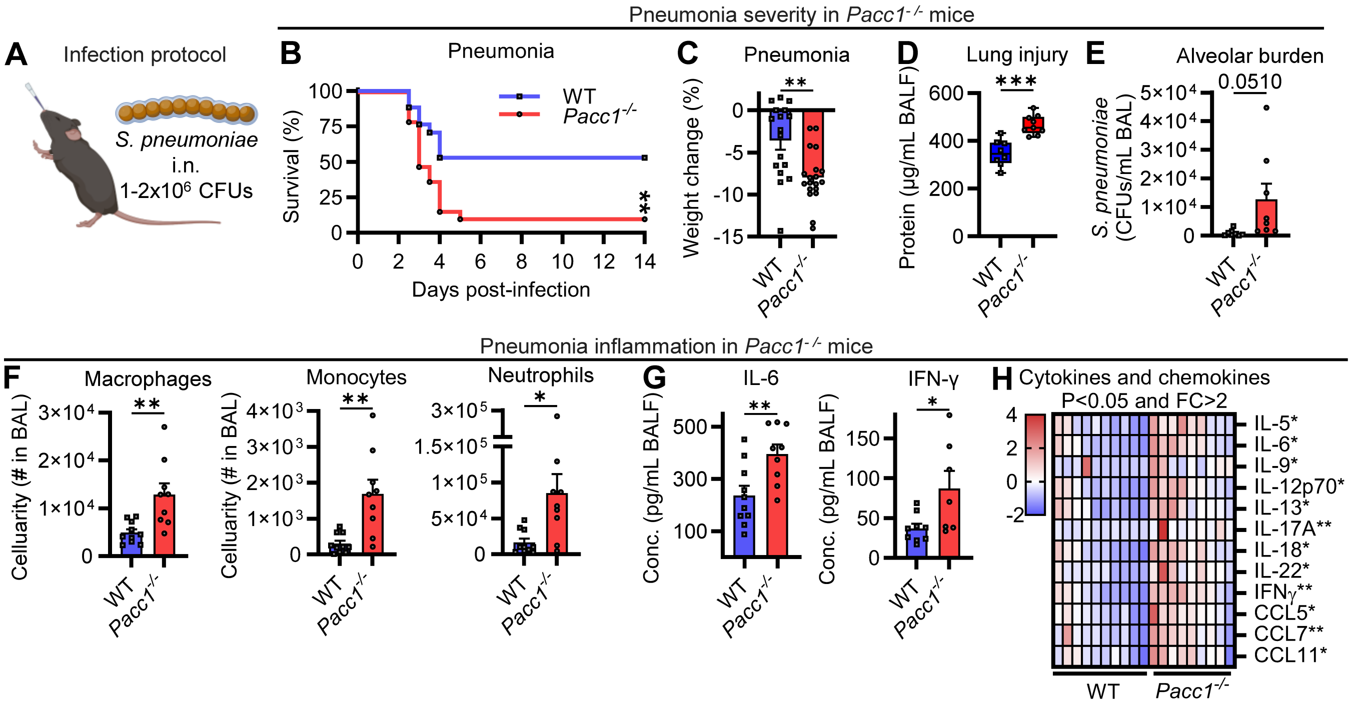


**Fig. S15. *Pacc1*$^{-/-}$ mice are susceptible to pneumococcal *pneumonia*.**

(**A**) Schematic showing pneumonia infection protocol. Mice were anesthetized (xylazine/ketamine) and given 1-2x10$^{6}$ colony-forming units (CFUs) of *S. pneumoniae*, serotype 4 (SPN4), intranasally (i.n.). Bronchoalveolar lavage (BAL) was collected 18 h later. (**B**) Pneumonia survival (n=9-11 mice/group, pooled from 2 independent experiments). (**C**) Pneumonia weight loss at day 2 post-infection. (**D**) Protein leakage in BAL fluid (BALF) after 18 h of infection as lung injury proxy by BCA assay. (**E**) CFUs in BAL after 18 h of infection assayed by serially diluted plated cultures. (**F**) Myeloid cellularity in BAL after 18 h of infection assessed by flow cytometry. (**G**) IL-6 and IFN-γ in BALF after 18 h of infection by ELISA. (**H**) Heatmap of significantly altered bronchoalveolar lavage fluid cytokines and chemokines showing >2-fold change (FC) after 18 h of infection; and analyzed by multiplex immunoassay. Data represent >2 independent experiments and show one symbol per mouse. Bar graphs depict mean±SEM, and box plots show median and interquartile range with whiskers extended to the minimum and maximum. *P<0.05, **P<0.01, ***P<0.001, Student's t-test (**C**-**H**), or Mantel-Cox log-rank test (**B**).

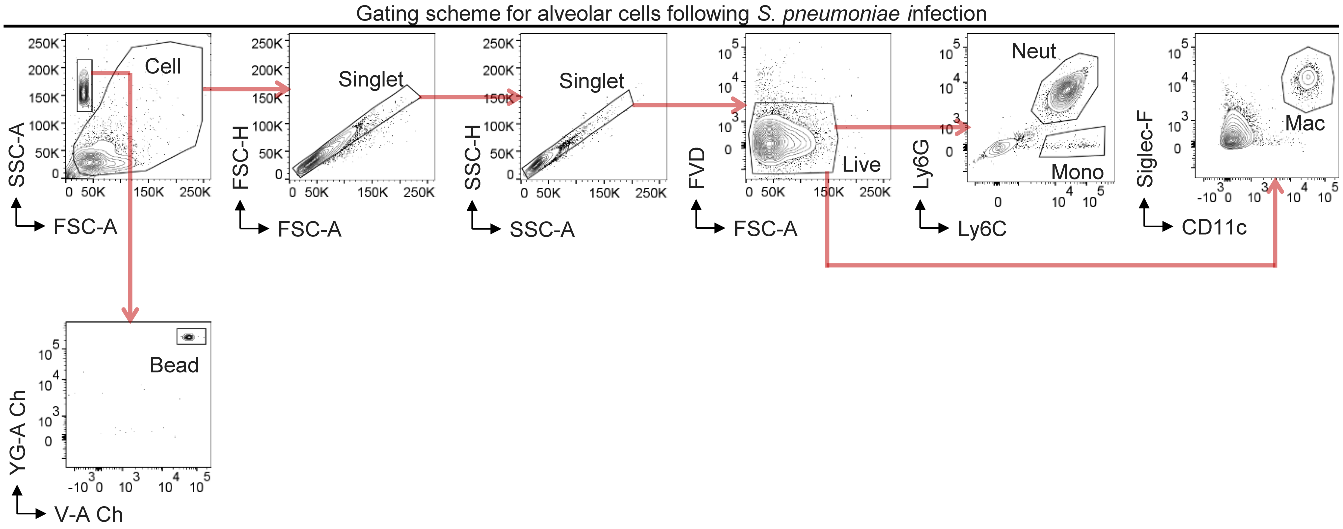


**Fig. S16. Gating scheme for alveolar cells following *S. pneumoniae* infection.**
Gating scheme for flow cytometric analysis of alveolar cells in bronchoalveolar lavage following infection with live *S. pneumoniae* as in **Fig. S15F**. Singlet cells were discriminated using forward scatter (FSC) and side scatter (SSC), and live cells (Live) were defined as negative for fixable viable dye eFluro780 (FVD). Neutrophils (Neut) were defined as Ly6C$^+$Ly6G$^+$, monocytes (Mono) as Ly6C$^+$Ly6G$^-$, and alveolar macrophages (Mac) as CD11c$^+$Siglec-F$^+$. Counting beads were discriminated based on low FSC-A and high autofluorescence in the indicated channels.

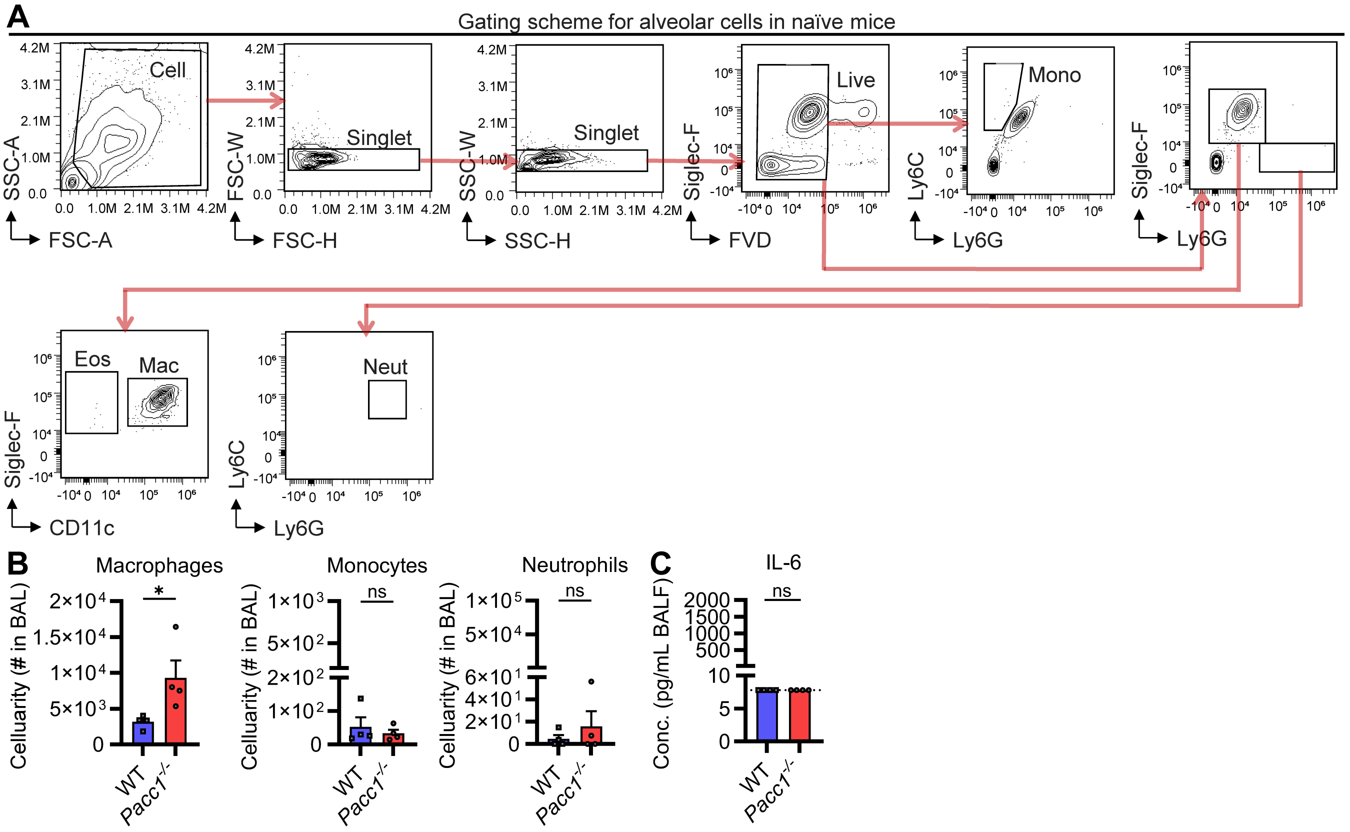


**Fig. S17. *Pacc1*$^{-/-}$ mice do not show significant aberrant inflammation in the lung at baseline.** (**A**) Gating scheme for flow cytometric analysis alveolar cells in bronchoalveolar lavage in naïve mice. Singlet cells were discriminated using forward scatter (FSC) and side scatter (SSC), and live cells were defined as negative for fixable viability dye eFluro780 (FVD). Neutrophils (Neut) were defined as Siglec-F$^{-}$Ly6C$^{+}$Ly6G$^{+}$, monocytes (Mono) as Ly6G$^{-}$Ly6C$^{+}$, alveolar macrophages (Mac) as CD11c$^{+}$Siglec-F$^{+}$, and eosinophils (Eos) as CD11c$^{-}$Siglec-F$^{+}$. (**B**) Cell numbers from the bronchoalveolar lavage (BAL) of naïve mice by flow cytometry. (**C**) IL-6 concentration in BAL fluid (BALF) of naïve mice by ELISA. The dotted line indicates the limit of detection. Data represent ≥2 independent experiments and show one symbol per mouse, mean±SEM, ns=not significant, Student's t-test.

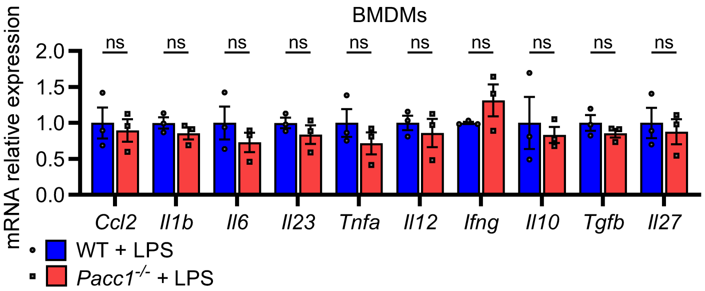


**Fig. S18.** ***Pacc1***$^{-/-}$ **bone marrow-derived macrophages (BMDMs) respond normally to lipopolysaccharide (LPS).**

LPS-induced gene expression in wildtype (WT) and *Pacc1*$^{-/-}$ BMDMs stimulated with LPS (100ng/mL) for 6 h. Gene expression measured by RT-qPCR and normalized to WT for each gene. Data represent ≥2 independent experiments and show one symbol per mouse, mean±SEM, ns=not significant, Student's t-test.

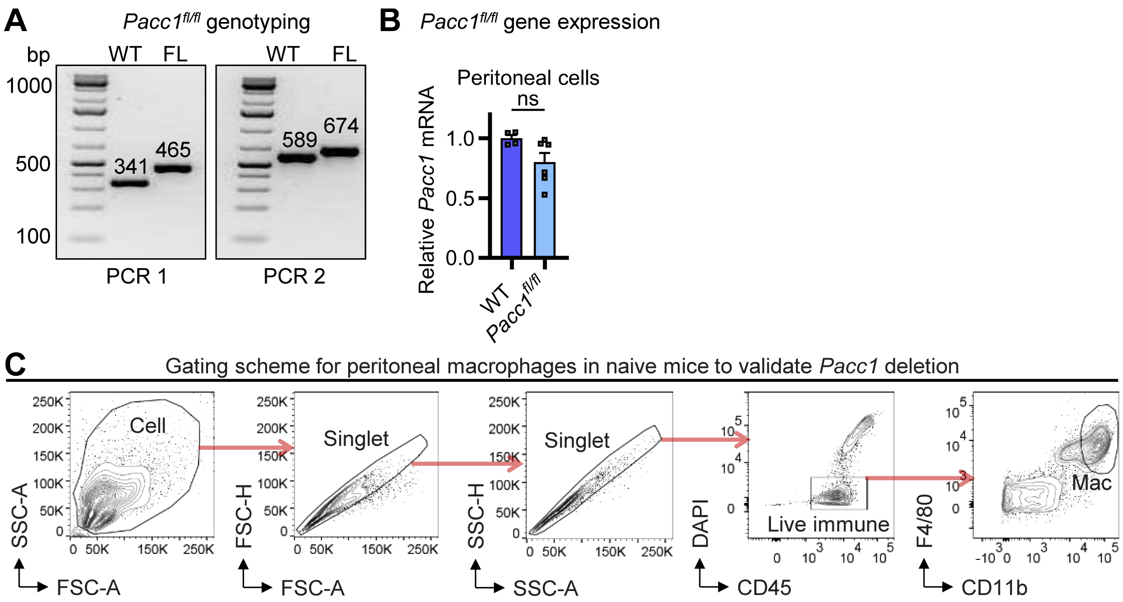


**Fig. S19. Genotyping and validating *Pacc1*-floxed mice.**

(**A**) Representative gels showing PCR-based genotyping of homozygous *Pacc1*-floxed (*Pacc1*$^{fl/fl}$) and wildtype (WT) mice. PCR 1 primers aligned to intron 3 resulted in an elongated band containing the loxP insertion at the indicated base pair (bp) length for the floxed (FL) band, as compared to the WT band. PCR 2 primers aligned to intron 4 also resulted in an elongated band containing the loxP sequence at the indicated bp length for the FL band, as compared to the WT band. (**B**) *Pacc1* expression was unchanged in total peritoneal lavage cells from naïve WT and *Pacc1*$^{fl/fl}$ mice by qRT-PCR. Data show one symbol per mouse, mean±SEM, ns=not significant, Student's t-test. (**C**) Gating scheme for flow cytometric analysis of peritoneal macrophages to validate *Pacc1* deletion in naïve *LysM-Cre/Pacc1*$^{fl/fl}$ vs. *Pacc1*$^{fl/fl}$ mice as in **Fig. 5L**. Singlet cells were discriminated using forward scatter (FSC) and side scatter (SSC), and live cells were defined as negative for DAPI. Immune cells (Live immune) were defined as CD45$^{+}$. Peritoneal macrophages were defined as CD45$^{+}$CD11b$^{hi}$F4/80$^{hi}$.

**Table S1. Effect size and p values associated with myeloid cells phenotypes in Fig. 1E; data retrieved from the OpenTargets platform.**

| Phenotype | Database | Reference | Variant (predicted function) | Beta (95% CI) | P value[#] |
|---|---|---|---|---|---|
| Monocyte count | OT Genetics | Chen MH, *et al.* (69) | rs701905 (pLoF) | 0.022 (0.018, 0.026) | $1.63\times10^{-28}$ |
| | | UK Biobank (70) | rs1389371 (pLoF) | 0.003 (0.002, 0.004) | $2.311\times10^{-10}$ |
| | | Astle WJ, *et al.* (71) | rs376261168 (inconclusive) | 0.024 (0.015, 0.032) | $1.57\times10^{-8}$ |
| | | Barton AR, *et al.* (72) | rs4500383 (pLoF) | 0.019 (0.015, 0.023) | $1.3\times10^{-21}$ |
| | | Vuckovic D, *et al.* (73) | rs6702347 (inconclusive) | 0.024 (0.019, 0.029) | $1\times10^{-23}$ |
| | | Sakaue S, *et al.* (74) | rs7542973 (inconclusive) | 0.018 (0.013, 0.022) | $3\times10^{-16}$ |
| | Gene Burden | Karczewski KJ, *et al.* (58) | Multiple variants (pLoF) | 0.004 (0.004, 0.005) | $1.792\times10^{-9}$ |
| Basophil count | OT Genetics | Chen MH, *et al.* (69) | rs1086893 (pLoF) | 0.03 (0.026, 0.034) | $2.41\times10^{-43}$ |
| | | | rs116522844 (pGoF) | -0.033 (-0.038, -0.028) | $5.54\times10^{-34}$ |
| | | UK Biobank (70) | rs116128776 (pGoF) | -0.021 (-0.026, -0.016) | $2\times10^{-15}$ |
| | | Astle WJ, *et al.* (71) | rs792448 (pLoF) | 0.028 (0.021, 0.035) | $2\times10^{-14}$ |
| | | Vuckovic D, *et al.* (73) | rs1086893 (pLoF) | 0.032 (0.028, 0.037) | $3.2\times10^{-43}$ |
| | | | rs12146141 (inconclusive) | -0.038 (-0.043, -0.032) | $3.8\times10^{-40}$ |
| | | Sakaue S, *et al.* (74) | rs12402642 (inconclusive) | 0.016 (0.012, 0.021) | $1\times10^{-14}$ |
| Basophil percentage of leukocytes | OT Genetics | UK Biobank (70) | rs903119 (inconclusive) | -0.017 (-0.021, -0.014) | $3.675\times10^{-20}$ |
| | | Astle WJ, *et al.* (71) | rs1774247 (inconclusive) | 0.028 (0.021, 0.035) | $8\times10^{-15}$ |
| | | Vuckovic D, *et al.* (73) | rs1086893 (pLoF) | 0.034 (0.03, 0.039) | $1.7\times10^{-48}$ |
| | Gene Burden | Karczewski KJ, *et al.* (58) | Multiple variants (pLoF) | 0.003 (0.003, 0.004) | $3.89\times10^{-10}$ |
| | | | Multiple variants (pLoF) | 0.005 (0.004, 0.005) | $1.919\times10^{-8}$ |
| Basophil % of granulocytes | OT Genetics | Astle WJ, *et al.* (71) | rs1774247 (inconclusive) | 0.03 (0.023, 0.037) | $1.223\times10^{-16}$ |
| Monocyte % of leukocytes | OT Genetics | UK Biobank (70) | rs1663626 (inconclusive) | 0.054 (0.041, 0.067) | $1.025\times10^{-16}$ |
| | | Astle WJ, *et al.* (71) | rs1663626 (inconclusive) | 0.022 (0.014, 0.029) | $3.538\times10^{-9}$ |
| | | Vuckovic D, *et al.* (73) | rs1663626 (inconclusive) | 0.026 (0.022, 0.031) | $8\times10^{-34}$ |
| Granulocyte % of myeloid white cells | OT Genetics | Astle WJ, *et al.* (71) | rs1663626 (inconclusive) | -0.023 (-0.03, -0.016) | $2.125\times10^{-10}$ |

[#] association p-value or burden test p-value; pGoF, predicted gain-of-function, (highlighted green); pLoF, predicted loss-of-function (highlighted red); inconclusive: inconclusive prediction based on available data on the effect of the variant on gene function. Data from Karczewski KJ, *et al.* (58) (Genebass) is from overall estimates from multiple pLoF variants. OT, Open Targets

**Table S2. Genotyping primers and single guide RNAs.**

| Strain | Target / Reaction | Primer | Sequence (5' ➞ 3') |
|---|---|---|---|
| ***Pacc1$^{-/-}$*** | Intron 1-2 | sgRNA 1 | GAAGGCGTGAACAGTTTAGCTGG |
| | Exon 2 | sgRNA 2 | CACTCACCTGGCACGACACCTGG |
| | PCR 1 | Forward 1 | TCTCCCTCACTGCTCATCGA |
| | | Reverse 1 | AGCTTTAGAACCGGGCAGAC |
| | PCR 2 | Forward 1 | TCTCCCTCACTGCTCATCGA |
| | | Reverse 2 | TCTCCCTCACTGCTCATCGA |
| ***Pacc1$^{fl/fl}$*** | Intron 2-3 | sgRNA 1 | TCCTTGCTAAGCACCCGACTTGG |
| | Intron 4-5 | sgRNA 2 | GGACCCTTATACACTGTGCACGG |
| | PCR 1 | Forward 1 | CTCTGCACTGTGTTCTGGTGCCATG |
| | | Reverse 1 | GTCCCTTGATGGCTAACATGACTGG |
| | PCR 2 | Forward 2 | TCTGGTAGCCACACACCCTGAGGAC |
| | | Reverse 2 | AACATACAACTGGGCCTAGCAAG |

**Table S3. Flow cytometry antibodies and reagents.**

| Antigen (or Viability) | Flurochrome (or Dye) | Target Species | Host Species | Isotype | Clone | Manufacturer | Catalogue # | Dilution |
|---|---|---|---|---|---|---|---|---|
| CD16/32 (TruStain fcX) | - | Mouse | Rat | IgG2a, λ | 93 | BioLegend | 101320 | 1:100 |
| B220 | V500 | Mouse/Human | Rat | IgG2a, κ | RA3-6B2 | BD Horizon | 561227 | 1:200 |
| CD11b | PECy7 | Mouse/Human | Rat | IgG2b, k | M1/70 | BioLegend | 101215 | 1:500 |
| CD11b | PB | Mouse/Human | Rat | IgG2b, κ | M1/70 | BioLegend | 101223 | 1:100 |
| CD11c | AF488 | Mouse | Hamster | IgG | N418 | BioLegend | 117313 | 1:100 |
| CD11c | PE-Cy7 | Mouse | Armenian Hamster | IgG | N418 | BioLegend | 117318 | 1:600 |
| CD19 | PerCP-Cy5.5 | Mouse | Rat | IgG2a, κ | J073E5 | BioLegend | 144515 | 1:100 |
| CD3 | BV510 | Mouse | Rat | IgG2b, κ | 17A2 | BioLegend | 100234 | 1:50 |
| CD3 | PB | Mouse | Rat | IgG2b, κ | 17A2 | BioLegend | 100214 | 1:50 |
| CD4 | PB | Mouse | Rat | IgG2a k | RM4-5 | BioLegend | 100534 | 1:200 |
| CD4 | FITC | Mouse | Rat | IgG2b, κ | RM4-5 | BioLegend | 100510 | 1:200 |
| CD49 | AF488 | Mouse | Rat | IgM, κ | DX5 | BioLegend | 108913 | 1:200 |
| CD8a | PECy7 | Mouse | Rat | IgG2a, κ | 53-6.7 | BioLegend | 100721 | 1:400 |
| CD8a | PE | Mouse | Rat | IgG2a, k | 53-6.7 | BioLegend | 100708 | 1:200 |
| F4/80 | PE | Mouse | Rat | IgG2a, κ | BM8 | BioLegend | 123109 | 1:200 |
| F4/80 | APC | Mouse | Rat | IgG2a, κ | BM8 | BioLegend | 123115 | 1:200 |
| Ly6C | BV510 | Mouse | Rat | IgG2c, κ | HK1.4 | BioLegend | 128033 | 1:100 |
| Ly6C | PE | Mouse | Rat | IgG2c, κ | HK1.4 | BioLegend | 128007 | 1:600 |
| Ly6C | BV510 | Mouse | Rat | IgG2c, κ | HK1.4 | BioLegend | 128033 | 1:100 |
| Ly6G | PECy7 | Mouse | Rat | IgG2a, κ | 1A8 | BioLegend | 127618 | 1:400 |
| Ly6G | APC | Mouse | Rat | IgG2a, κ | 1A8 | BioLegend | 127613 | 1:400 |
| Ly6G | PE | Mouse | Rat | IgG2a, κ | 1A8 | BioLegend | 127607 | 1:400 |
| MHC-II (I-A/I-E) | PerCP | Mouse | Rat | IgG2b, k | M5/114.15.2 | BioLegend | 107624 | 1:400 |
| NK1.1 | APC | Mouse | Mouse | IgG2a, κ | PK136 | BioLegend | 108709 | 1:50 |
| NKp46 | PE | Mouse | Rat | IgG2a, k | 29A1.4 | BioLegend | 137604 | 1:100 |
| *Pacc1*-RNA | APC | Mouse | - | - | - | Thermo Fisher | PF-210 | - |
| SiglecF | BV421 | Mouse | Rat | IgG2a, k | E50-2440 | BD Horizon | 562681 | 1:400 |
| SiglecF | APC | Mouse | Rat | IgG1 | ES22-10D8 | Miltenyi | 130-123-816 | 1:100 |
| Viability | DAPI | - | - | - | - | Thermo Fisher | R37606 | 1:25 |
| Viability | FVD eF780 | - | - | - | - | eBioscience | 65-0865-14 | 1:1000 |
| Viability | Propidium Iodide | - | - | - | - | Invitrogen | L34856 | 1:667 |
| Viability | Syto 9 | - | - | - | - | Invitrogen | L34856 | 1:667 |
| Viability | 7-AAD | - | - | - | - | BioLegend | 420403 | 1:1000 |

### Table S4. RT-qPCR primer sequences for *Gapdh* and *Pacc1*.

| qRT-PCR primers | | |
|---|---|---|
| **Primer** | **Forward Sequence (5' ➞ 3')** | **Reverse Sequence (5' ➞ 3')** |
| Mouse *Gapdh* | TACCCCCAATGTGTCCGTCGTG | CCTTCAGTGGGCCCTCAGATGC |
| Mouse *Pacc1 Exon 2/3* | CGAGGAGTTGGAGCAGGTGGTTG | TGTAAGACACGGACATGACAGGGTG |
| Mouse *Pacc1 Exon 3/4* | AGTCCGCCTCCAGCAGCATC | TGCAGTTCCTGTCTCCCGGCTG |
| Mouse *Pacc1 Exon 5/6* | TGTCCAGGGGCCCCAGGAAG | ACAGCTTCTCGCCCGTCCTC |
| Mouse *Pacc1 Exon 7/8* | AGCCGAGAGGAGTGCTCAGTTG | TGTCGCCTGGCCTCTCCG |

### References


1. S. Lykke-Andersen, T. H. Jensen, Nonsense-mediated mRNA decay: an intricate machinery that shapes transcriptomes. *Nature Reviews Molecular Cell Biology* **16**, 665-677 (2015).
2. J. P. Concordet, M. Haeussler, CRISPOR: intuitive guide selection for CRISPR/Cas9 genome editing experiments and screens. *Nucleic Acids Res* **46**, W242-w245 (2018).
3. A. Hodgkins *et al.*, WGE: a CRISPR database for genome engineering. *Bioinformatics* **31**, 3078-3080 (2015).
4. M. Stemmer, T. Thumberger, M. Del Sol Keyer, J. Wittbrodt, J. L. Mateo, CCTop: An Intuitive, Flexible and Reliable CRISPR/Cas9 Target Prediction Tool. *PLoS One* **10**, e0124633 (2015).
5. M. Bosmann *et al.*, Interruption of macrophage-derived IL-27(p28) production by IL-10 during sepsis requires STAT3 but not SOCS3. *J Immunol* **193**, 5668-5677 (2014).
6. J. Roewe *et al.*, Bacterial polyphosphates interfere with the innate host defense to infection. *Nat Commun* **11**, 4035 (2020).
7. B. Lindner, T. Burkard, M. Schuler, Phagocytosis assays with different pH-sensitive fluorescent particles and various readouts. *Biotechniques* **68**, 245-250 (2020).
8. A. Neaga, J. Lefor, K. E. Lich, S. F. Liparoto, Y. Q. Xiao, Development and validation of a flow cytometric method to evaluate phagocytosis of pHrodo™ BioParticles® by granulocytes in multiple species. *J Immunol Methods* **390**, 9-17 (2013).
9. S. Andrews (2010) FastQC: a quality control tool for high throughput sequence data. Available online at: http://www.bioinformatics.babraham.ac.uk/projects/fastqc.
10. F. Krueger *et al.* (2023) FelixKrueger/TrimGalore: v0.6.10 - add default decompression path (0.6.10). Zenodo. https://doi.org/10.5281/zenodo.7598955.
11. M. Martin, Cutadapt removes adapter sequences from high-throughput sequencing reads. *2011* **17**, 3 (2011).
12. O. Tange (2018) GNU Parallel 2018, Mar 2018, ISBN 9781387509881, DOI https://doi.org/10.5281/zenodo.1146014.
13. F. Mölder *et al.*, Sustainable data analysis with Snakemake. *F1000Research* **10**, 33 (2021).
14. A. Dobin *et al.*, STAR: ultrafast universal RNA-seq aligner. *Bioinformatics* **29**, 15-21 (2013).
15. P. Danecek *et al.*, Twelve years of SAMtools and BCFtools. *Gigascience* **10** (2021).
16. Y. Liao, G. K. Smyth, W. Shi, featureCounts: an efficient general purpose program for assigning sequence reads to genomic features. *Bioinformatics* **30**, 923-930 (2014).
17. P. Ewels, M. Magnusson, S. Lundin, M. Käller, MultiQC: summarize analysis results for multiple tools and samples in a single report. *Bioinformatics* **32**, 3047-3048 (2016).
18. M. I. Love, W. Huber, S. Anders, Moderated estimation of fold change and dispersion for RNA-seq data with DESeq2. *Genome Biol* **15**, 550 (2014).
19. S. Durinck, P. T. Spellman, E. Birney, W. Huber, Mapping identifiers for the integration of genomic datasets with the R/Bioconductor package biomaRt. *Nat Protoc* **4**, 1184-1191 (2009).
20. F. Marini, H. Binder, pcaExplorer: an R/Bioconductor package for interacting with RNA-seq principal components. *BMC Bioinformatics* **20**, 331 (2019).
21. A. Subramanian *et al.*, Gene set enrichment analysis: a knowledge-based approach for interpreting genome-wide expression profiles. *Proc Natl Acad Sci U S A* **102**, 15545-15550 (2005).
22. A. S. Castanza *et al.*, Extending support for mouse data in the Molecular Signatures Database (MSigDB). *Nat Methods* **20**, 1619-1620 (2023).
23. I. Dolgalev (2022) msigdbr: MSigDB Gene Sets for Multiple Organisms in a Tidy Data Format. R package version 7.5.1, <https://CRAN.R-project.org/package=msigdbr>.
24. T. Wu *et al.*, clusterProfiler 4.0: A universal enrichment tool for interpreting omics data. *Innovation (Camb)* **2**, 100141 (2021).

25. A. Ludt, A. Ustjanzew, H. Binder, K. Strauch, F. Marini, Interactive and Reproducible Workflows for Exploring and Modeling RNA-seq Data with pcaExplorer, Ideal, and GeneTonic. *Curr Protoc* **2**, e411 (2022).
26. K. Guo, B. McGregor (2024) VennDetail: A package for visualization and extract details. R package version 1.22.0, https://github.com/guokai8/VennDetail.
27. K. Blighe (2020) Tutorial:Clustering of DAVID gene enrichment results from gene expression studies. Available from: https://www.biostars.org/p/299161/.
28. Z. Gu, Complex heatmap visualization. *Imeta* **1**, e43 (2022).
29. H. Wickham, *ggplot2: Elegant Graphics for Data Analysis.* (Springer-Verlag New York, 2016), pp. 260.
30. Anonymous (Sino Biological. Other Enzymes Products Center. Retrieved June 27, 2025, from https://www.sinobiological.com/research/enzymes/other-enzymes-products-center.
31. A. Kramer, J. Green, J. Pollard, Jr., S. Tugendreich, Causal analysis approaches in Ingenuity Pathway Analysis. *Bioinformatics* **30**, 523-530 (2014).
32. I. Rusinova *et al.*, Interferome v2.0: an updated database of annotated interferon-regulated genes. *Nucleic Acids Res* **41**, D1040-1046 (2013).
33. A. J. Stagg, F. Burke, S. Hill, S. C. Knight, Isolation of mouse spleen dendritic cells. *Methods Mol Med* **64**, 9-22 (2001).
34. J. Roewe *et al.*, Bacterial polyphosphates induce CXCL4 and synergize with complement anaphylatoxin C5a in lung injury. *Front Immunol* **13**, 980733 (2022).
35. M. Dudek *et al.*, Lung epithelium and myeloid cells cooperate to clear acute pneumococcal infection. *Mucosal Immunol* **9**, 1288-1302 (2016).
36. M. Bosmann, N. F. Russkamp, P. A. Ward, Fingerprinting of the TLR4-induced acute inflammatory response. *Exp Mol Pathol* **93**, 319-323 (2012).
37. A. Sharma *et al.*, IL-27 Enhances gammadelta T Cell-Mediated Innate Resistance to Primary Hookworm Infection in the Lungs. *J Immunol* **208**, 2008-2018 (2022).
38. RCoreTeam, R: A language and environment for statistical computing. R Foundation for Statistical Computing, Vienna, Austria. URL https://www.R-project.org/. (2022).
39. RStudioTeam, RStudio: Integrated Development Environment for R. RStudio, PBC, Boston, MA URL http://www.rstudio.com/. (2022).
40. T. Stuart, A. Srivastava, S. Madad, C. A. Lareau, R. Satija, Single-cell chromatin state analysis with Signac. *Nat Methods* **18**, 1333-1341 (2021).
41. H. Wickham, *ggplot2: Elegant Graphics for Data Analysis* (Springer International Publishing, 2016).
42. Anonymous (PACC1: phenotype association scores from the Open Targets Platform. Available at: https://platform.opentargets.org/target/ENSG00000065600/associations . Accessed on: 13 March 2024
43. D. Ochoa *et al.*, The next-generation Open Targets Platform: reimagined, redesigned, rebuilt. *Nucleic Acids Res* **51**, D1353-d1359 (2023).
44. M. Ghoussaini *et al.*, Open Targets Genetics: systematic identification of trait-associated genes using large-scale genetics and functional genomics. *Nucleic Acids Research* **49**, D1311-D1320 (2020).
45. M. J. Landrum *et al.*, ClinVar: public archive of relationships among sequence variation and human phenotype. *Nucleic Acids Res* **42**, D980-985 (2014).
46. M. J. Landrum *et al.*, ClinVar: improvements to accessing data. *Nucleic Acids Res* **48**, D835-d844 (2020).
47. T. Cezard *et al.*, The European Variation Archive: a FAIR resource of genomic variation for all species. *Nucleic Acids Res* **50**, D1216-d1220 (2022).
48. A. Shen *et al.*, CMAT: ClinVar Mapping and Annotation Toolkit. *Bioinform Adv* **4**, vbae018 (2024).

49. Anonymous, Ultra-Rare Genetic Variation in the Epilepsies: A Whole-Exome Sequencing Study of 17,606 Individuals. *Am J Hum Genet* **105**, 267-282 (2019).
50. P. Akbari *et al.*, Multiancestry exome sequencing reveals INHBE mutations associated with favorable fat distribution and protection from diabetes. *Nat Commun* **13**, 4844 (2022).
51. J. D. Backman *et al.*, Exome sequencing and analysis of 454,787 UK Biobank participants. *Nature* **599**, 628-634 (2021).
52. L. Bomba *et al.*, Whole-exome sequencing identifies rare genetic variants associated with human plasma metabolites. *Am J Hum Genet* **109**, 1038-1054 (2022).
53. M. B. Makarious *et al.*, Large-scale rare variant burden testing in Parkinson's disease. *Brain* **146**, 4622-4632 (2023).
54. F. Riveros-Mckay *et al.*, The influence of rare variants in circulating metabolic biomarkers. *PLoS Genet* **16**, e1008605 (2020).
55. F. K. Satterstrom *et al.*, Large-Scale Exome Sequencing Study Implicates Both Developmental and Functional Changes in the Neurobiology of Autism. *Cell* **180**, 568-584.e523 (2020).
56. T. Singh *et al.*, Rare coding variants in ten genes confer substantial risk for schizophrenia. *Nature* **604**, 509-516 (2022).
57. X. Zhou *et al.*, Integrating de novo and inherited variants in 42,607 autism cases identifies mutations in new moderate-risk genes. *Nat Genet* **54**, 1305-1319 (2022).
58. K. J. Karczewski *et al.*, Systematic single-variant and gene-based association testing of thousands of phenotypes in 394,841 UK Biobank exomes. *Cell Genom* **2**, 100168 (2022).
59. S. J. Jurgens *et al.*, Rare coding variant analysis for human diseases across biobanks and ancestries. *Nat Genet* **56**, 1811-1820 (2024).
60. Anonymous (PACC1: predicted Loss-of-function exome-based association statistics from the Genebass database. Retrieved from: https://app.genebass.org/gene/ENSG00000065600?burdenSet=pLoF&phewasOpts=1&resultLayout=full . Accessed on 13 March 2024.
61. C. Bycroft *et al.*, The UK Biobank resource with deep phenotyping and genomic data. *Nature* **562**, 203-209 (2018).
62. G. Monaco *et al.*, RNA-Seq Signatures Normalized by mRNA Abundance Allow Absolute Deconvolution of Human Immune Cell Types. *Cell Rep* **26**, 1627-1640.e1627 (2019).
63. M. Karlsson *et al.*, A single-cell type transcriptomics map of human tissues. Further details available at: https://www.proteinatlas.org/about/assays+annotation#normalization_rna. *Sci Adv* **7** (2021).
64. C. Klijn *et al.*, A comprehensive transcriptional portrait of human cancer cell lines. *Nat Biotechnol* **33**, 306-312 (2015).
65. Anonymous (R Core Team (2024). R: A Language and Environment for Statistical Computing_. R Foundation for Statistical Computing, Vienna, Austria. <https://www.R-project.org/>.
66. J. Qie *et al.*, Integrated proteomic and transcriptomic landscape of macrophages in mouse tissues. *Nature Communications* **13**, 7389 (2022).
67. T. S. Heng, M. W. Painter, The Immunological Genome Project: networks of gene expression in immune cells. *Nat Immunol* **9**, 1091-1094 (2008).
68. K. Li *et al.*, Profiling phagosome proteins identifies PD-L1 as a fungal-binding receptor. *Nature* **630**, 736-743 (2024).
69. M.-H. Chen *et al.*, Trans-ethnic and Ancestry-Specific Blood-Cell Genetics in 746,667 Individuals from 5 Global Populations. *Cell* **182**, 1198-1213.e1114 (2020).
70. Anonymous (An updated GWAS analysis of the UK Biobank, 2018. Available from: http://www.nealelab.is/uk-biobank/
71. W. J. Astle *et al.*, The Allelic Landscape of Human Blood Cell Trait Variation and Links to Common Complex Disease. *Cell* **167**, 1415-1429.e1419 (2016).

72. A. R. Barton, M. A. Sherman, R. E. Mukamel, P.-R. Loh, Whole-exome imputation within UK Biobank powers rare coding variant association and fine-mapping analyses. *Nat Genet* **53**, 1260-1269 (2021).
73. D. Vuckovic *et al.*, The Polygenic and Monogenic Basis of Blood Traits and Diseases. *Cell* **182**, 1214-1231.e1211 (2020).
74. S. Sakaue *et al.*, A cross-population atlas of genetic associations for 220 human phenotypes. *Nat Genet* **53**, 1415-1424 (2021).

### Description of Supplementary Datasets

**Dataset S1. Normalized counts and differential expression data from bulk RNA sequencing analysis of *Pacc1*$^{-/-}$ and wildtype bone marrow-derived macrophages (BMDMs) stimulated with *Escherichia (E.) coli* BioParticles or left as unstimulated controls.**

**Dataset S2. Gene set enrichment analysis (GSEA) of differentially expressed genes in MSigDB Hallmark gene sets in *Pacc1*$^{-/-}$ and wildtype bone marrow-derived macrophages (BMDMs) stimulated with *Escherichia (E.) coli* BioParticles or left as unstimulated controls.**

**Dataset S3. Canonical pathways from Ingenuity Pathway Analysis (IPA) from significantly differentially expressed genes in *Pacc1*$^{-/-}$ vs. wildtype bone marrow-derived macrophages (BMDMs) stimulated with *Escherichia (E.) coli* BioParticles.**

**Dataset S4. Gene ontology biological process (GOBP) enrichment of significant differentially expressed genes in *Pacc1*$^{-/-}$ and wildtype bone marrow-derived macrophages (BMDMs) stimulated with *Escherichia (E.) coli* BioParticles or left as unstimulated controls.**
